# Regional and sub-regional microglial heterogeneity in the steady-state mouse brain and retina

**DOI:** 10.64898/2026.01.16.699866

**Authors:** Fazeleh Etebar, Paul Whatmore, Damien G. Harkin, Hazel Quek, Elisa Eme-Scolan, Paul G. McMenamin, Samantha J. Dando

## Abstract

CNS-resident immune cells are uniquely adapted to their microenvironment; however, the extent of their regional specialisation remains unclear. We combined morphometric and transcriptomic profiling of microglia across the healthy adult mouse CNS, including the olfactory bulbs, cortex, hippocampus, cerebellum and retina, to define their regional and sub-regional heterogeneity. Bulk RNA-sequencing revealed region-specific signatures, with retinal microglia showing the most divergent transcriptomes, and genes related to antigen presentation, phagocytosis and chemokine signalling among the top differentially expressed genes. Single-cell RNA sequencing identified predominantly homeostatic microglia across all examined regions, alongside smaller clusters of interferon-responsive, chemokine-enriched, apolipoprotein-enriched and proliferative microglia. Apolipoprotein-enriched microglia were restricted to the olfactory bulbs, whereas interferon-responsive microglia were most abundant in the retina. Single-cell profiling of human retinal microglia confirmed clusters enriched for interferon-stimulated genes. Together, this study reveals previously unrecognised microglial heterogeneity within the healthy brain and eye and provides a comparison of microglia transcriptomes across different neuroanatomical regions of the CNS.

## Background

Microglia are tissue-resident macrophages with roles in CNS development, homeostasis, inflammation, and injury repair (1). Microglia actively sense perturbations in the CNS environment through their highly ramified motile processes (2), and have specific functions in the non-diseased CNS including recognizing and phagocytosing apoptotic neurons, learning-related synapse formation, maintaining neurons, regulating neurogenesis, and contributing to synaptic structure and function (3–12).

In recent years, single cell ‘omics’ studies have revealed significant microglial plasticity during development and pathological conditions (13–15). Distinct microglia clusters, including proliferative region-associated microglia (PAM) (16) and specialized axon tract-associated microglia (ATM) (14), emerge during specific developmental periods, each exhibiting a unique transcriptional signature. Injury-responsive microglia (IRM) display a unique gene expression pattern in a spinal cord injury animal model, suggesting involvement in a tailored response to injury (14). In neurodegenerative disease, several microglial subpopulations have been described including disease-associated microglia (DAM) (17, 18), activated response microglia (ARM) (19), and microglia associated with neurodegenerative disease (MGnD) (20), each with distinct functions and gene expression profiles.

The CNS regional microenvironment also impacts microglial function, phenotype, and transcription profiles; with differences observed between microglia located in different CNS regions in pathological states (21–25). However, there are conflicting studies regarding the diversity of microglia within the healthy adult CNS (14, 26). Using scRNA-seq, Hammond *et al*. demonstrated heterogeneous subpopulations of microglia within the developing mouse brain and, to a lesser extent, the aged mouse brain (14). However, little microglial heterogeneity was observed in the juvenile and adult mouse brain. In contrast, using bulk RNA-seq Grabert *et al.* demonstrated distinct region-dependent transcriptional identities of microglia isolated from the healthy adult mouse brain (21). Furthermore, our lab also reported regional differences in the distribution of CD11c-expressing microglia in the healthy mouse CNS. We demonstrated that these cells accumulated primarily in the inner retina and brain regions lacking a blood-brain barrier (BBB) or vulnerable to microbial invasion (27). Recent studies identified functionally specialized microglia populations in the healthy adult mouse CNS, including capillary-associated microglia (28) and pericyte-associated microglia (29). Taken together, evidence suggests that there may be greater microglial diversity in the healthy adult CNS than previously thought.

The aim of the present study was to investigate the heterogeneity of microglia within five easily dissected discrete regions of the healthy adult mouse CNS, including the neural retina, olfactory bulbs, cerebral cortex, hippocampus, and cerebellum. We isolated microglia from each of these regions and performed gene expression profiling using bulk RNA-seq and scRNA-seq to generate a comprehensive resource of microglia gene transcriptomes and compare the regional- and sub-regional heterogeneity of microglia within defined neuroanatomical regions of the mouse brain and eye. The data demonstrated region-specific microglial gene expression profiles and identified small clusters of transcriptionally unique microglia within the investigated CNS regions. Our findings may serve as a base for future studies aimed at understanding the functions of microglia subpopulations in neurological and ocular diseases.

## Results

### Regional heterogeneity of microglia within the steady-state adult mouse CNS

We initially performed density and morphometric comparisons of microglia within brain sections and retinal wholemounts (Figure 1a) of healthy adult Cx3cr1-gfp/+ mice, in which microglia are readily identifiable by constitutive expression of GFP. In the brain, the density of microglia was significantly higher in the olfactory bulbs, cortex and hippocampus compared to the cerebellum (Figure 1b). Retinal microglial density varied according to tissue layer, with significantly higher densities observed in the inner plexiform layer (IPL) and outer plexiform layer (OPL) compared to the ganglion cell layer (GCL) (Figure 1b).

**Figure 1.**
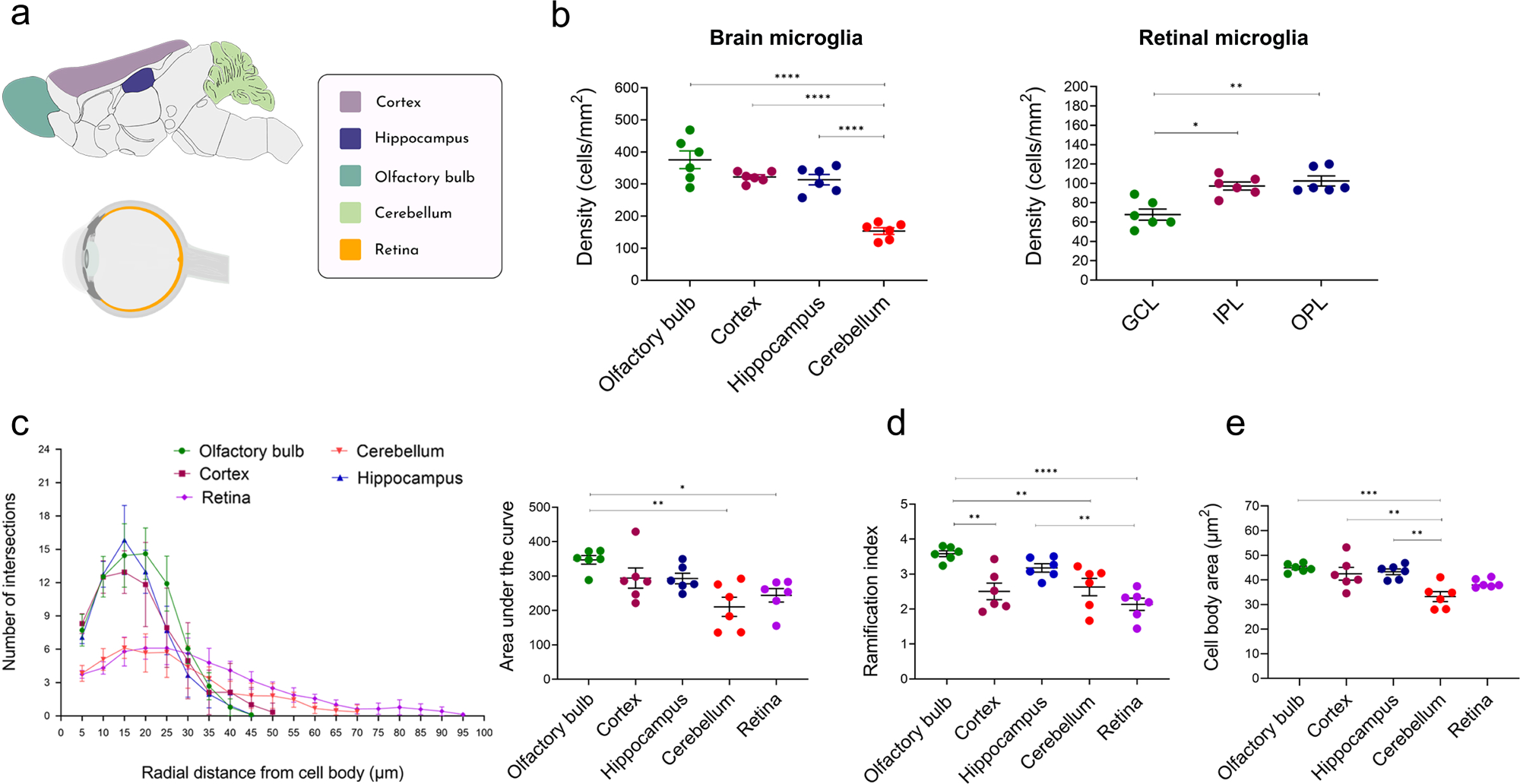
Regional differences in microglial density and morphometry in the brain and retina of healthy Cx3cr1^gfp/+^ mice. **(a)** Schematic of CNS regions analysed (olfactory bulb, cortex, hippocampus, cerebellum, and retina). **(b)** quantification of microglial density (cells/mm²) in the brain and laminar layers of the retina where microglia reside (ganglion cell layer [GCL], inner plexiform layer [IPL], outer plexiform layer [OPL]. **(c)** Sholl plot showing the radial distance of microglia processes from cell body and the number of intersections at each distance; and the area under the curve of the microglia Sholl profile. Comparison of microglial ramification index **(d)** and cell body area **(e)** across CNS regions. Data points in graphs represent averaged data for each mouse, obtained from analysis of n=6 microglia per brain region and n=18 retinal microglia per mouse (comprising n=6 microglia from the GCL, IPL and OPL). Data were analysed using one-way ANOVA with multiple comparisons test (* p<0.05, ** p<0.01, *** p<0.001 and **** p<0.0001). Mean ± SEM shown.

Significant morphometric differences were observed between steady-state microglia within different regions of the mouse CNS. Sholl analysis demonstrated that retinal and cerebellar microglia displayed fewer branch points (intersections) and longer processes compared to cortical, hippocampal and olfactory bulb microglia (Figure 1c). Olfactory bulb microglia displayed the highest ramification index, whereas retinal microglia were less ramified than brain microglia (Figure 1d). Additionally, cerebellar microglia had a significantly smaller cell body area compared to microglia in other brain regions (Figure 1e).

As microglial morphology is frequently used as an indicator of ‘activation’ status (30), we investigated whether the observed regional differences extended beyond morphology into gene expression. Bulk RNA-seq was performed to compare the transcriptome of microglia isolated from the olfactory bulbs, cortex, hippocampus, cerebellum and retina of the healthy adult C57Bl/6J mouse CNS (Figure S1). Microglia from each region expressed homeostatic microglia genes (such as *Tmem119*, *P2ry12*, *Scl2a5* and *Fcrls*) and myeloid cell genes (such as *Cx3cr1*, *Itgam*, *CD45*, *Aif1*). Of these genes, only *Fcrls* was differentially expressed (as determined by a false discovery rate (FDR) <0.05 and a log_2_ fold change (log_2_ FC) of >1) between regional microglia populations. Both cerebellar microglia (mean log_2_ FC −1.25) and retinal microglia (mean log_2_ FC −2.07) expressed lower levels of *Fcrls* compared to cortical microglia (Figure S2), suggesting that this gene may not be suitable as a universal homeostatic gene in microglia. A total of 747 differentially expressed genes (DEGs) were identified between microglial populations. Heat mapping of DEGs (Figure 2a) demonstrated that cortical and hippocampal microglia shared similar gene expression profiles. Cerebellar microglia had some unique DEGs compared to other brain microglia, whereas olfactory bulb microglia displayed an intermediate profile, sharing some DEGs with cortical and hippocampal microglia, and some with cerebellar microglia. Interestingly, retinal microglia were transcriptionally distinct from brain microglia (Figure 2a). Compared to cortical microglia, 103 genes were differentially expressed in hippocampal microglia, 222 and 230 genes were differentially expressed in cerebellar and olfactory bulb microglia respectively, and 447 genes were differentially expressed in retinal microglia. The greatest log_2_ fold changes in gene expression were observed when comparing retinal microglia to cortical microglia (Figure 2b).

**Figure 2.**
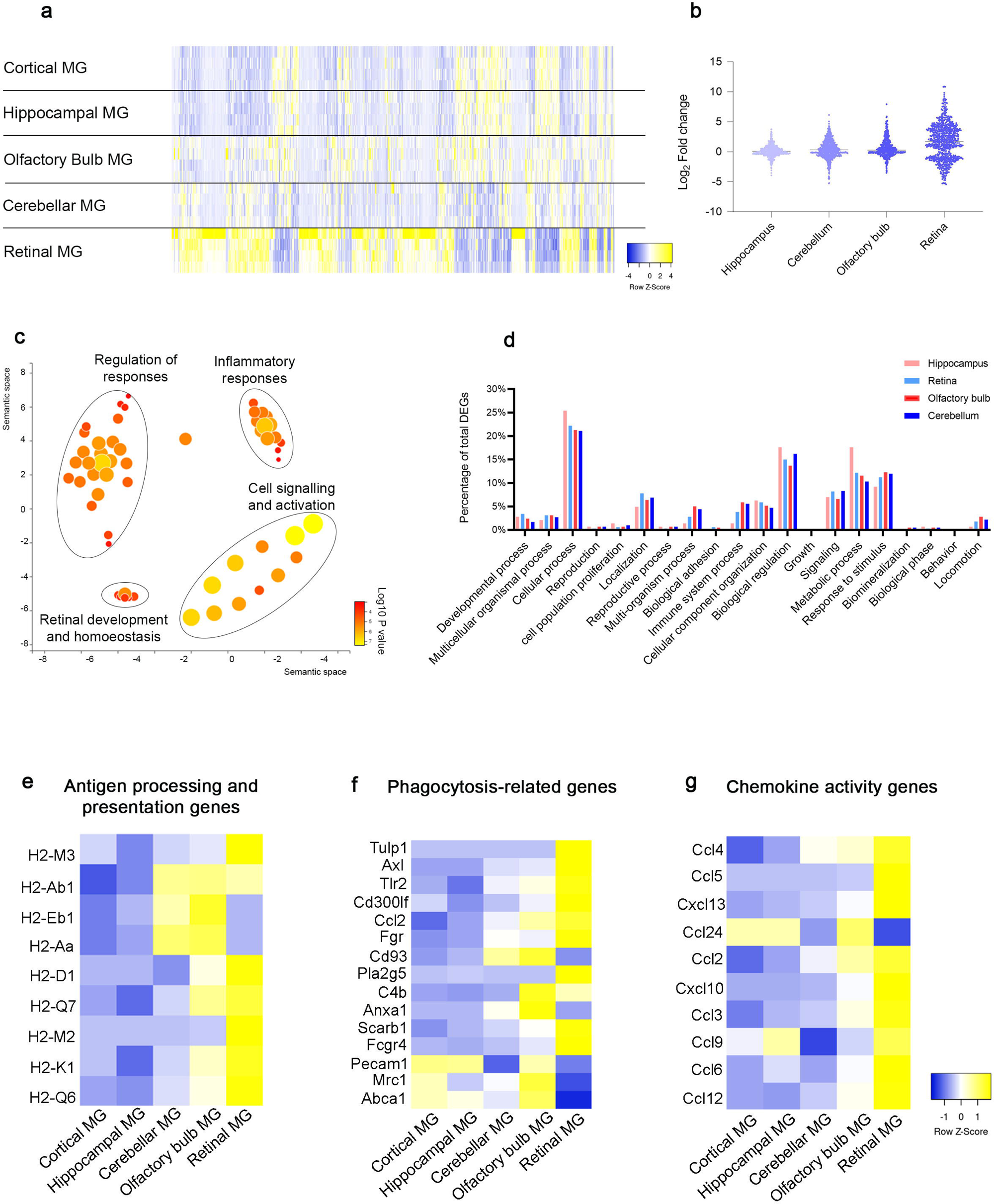
Bulk RNA-seq comparison of differentially expressed genes in microglia isolated from different regions of the healthy mouse CNS. **(a)** Heat map clustering of differentially expressed genes (DEGs, FDR <0.05, Log_2_ fold-change >1) based on scaled expression values (Z score). N= 4 biological replicates per CNS region. (**b)** Violin plot demonstrating the spread of log_2_ fold-change gene expression relative to cortical microglia. (**c)** Revigo clustering of GO terms overrepresented in DEGs. (**d)** Comparison of enriched GO terms for biological process in microglia isolated from different regions of the mouse CNS. Input genes for GO analysis were filtered according to FDR < 0.05 and log_2_ fold change of >1 compared to cortical microglia. **(e-g)** Heatmaps demonstrating differential expression of genes involved in: **(e)** antigen processing and presentation of exogenous and endogenous peptide antigens via MHC class II and class I; **(f)** phagocytosis functions; **(g)** chemokine activity. Heatmaps were generated based on scaled expression values (Z score).

DEGs were assessed for enriched terms and pathways using gene ontology (GO) to determine over-represented biological processes (Supplementary file 1). Revigo grouped the GO terms into 4 main clusters related to: (i) regulation of responses, (ii) inflammatory responses, (iii) cell signalling and activation, and (iv) retinal development and homoeostasis (Figure 2c). The GO terms ‘cellular process’, ‘biological regulation’, ‘metabolic process’, ‘response to stimulus’ and ‘signalling’ comprised the largest proportion of DEGs (Figure 2d).

The 747 DEGs were classified according to their biological process using the Uniprot database. Of these, 196/747 (26%) DEGs were involved in immune function (Supplementary file 2), and a large portion of these immune-related DEGs were upregulated in cerebellar, olfactory bulb and retinal microglia compared to cortical microglia. Immune-related DEGs were then categorized into groups based on GO classifications including: (i) genes involved in antigen processing and presentation of endogenous and exogenous peptide antigen via MHC class I and class II; (ii) genes with phagocytosis function, and (iii) genes with chemokine activity. Genes related to antigen presentation via MHC class II (including *H2-Aa*, *H2-Ab1* and *H2-Eb1*) were highly expressed in cerebellar and olfactory bulb microglia (but not hippocampal or retinal microglia). Retinal microglia had the highest expression of MHC class I genes (*H2-M2*, *H2-Q6*, *H2-Q7*, *H2-D1* and *H2-K1*) compared to brain microglia populations (Figure 2e). Genes involved in phagocytosis were also highly expressed in retinal microglia. For example, *Tulp1* (log_2_ FC= 8.7), which contributes to stimulation of phagocytosis of apoptotic cells (31), and *Axl* (log_2_ FC= 4.3), which regulates innate immune responses and phagocytosis (32, 33), were highly expressed in retinal microglia. In contrast, some phagocytosis related genes (e.g. *Anxa1* and *Cd93*) were upregulated in olfactory bulb and cerebellar microglia but downregulated in cortical, hippocampal and retinal microglia (Figure 2f). Consistent with the increased expression of antigen processing and presentation genes, a large number of chemokine-related genes were also highly expressed in retinal microglia. Notably, retinal microglia had a distinct expression profile for genes encoding the chemokines *Cxcl13* (log_2_ FC= 4.3), *Cxcl10* (log_2_ FC= 5.7), and *Ccl5* (log_2_ FC= 4.9) (Figure 2g).

These data demonstrate that microglia in the healthy adult mouse CNS have regionally distinct gene expression profiles. Our findings suggest that microglia in the retina, cerebellum and olfactory bulbs may be more equipped to respond to danger signals compared to cortical and hippocampal microglia. However, using this bulk RNA-seq approach it is not clear whether all microglia within these regions possess the same transcriptomic signature, or whether these findings are attributed to unique microglia subtypes within these CNS regions.

### scRNA-seq confirms microglia have region-dependent gene expression profiles

To develop a comprehensive understanding of intra-regional microglia diversity in healthy mice, scRNA-seq was used to determine the transcriptomic profile of microglia isolated from the olfactory bulbs, hippocampus, cortex, cerebellum, and retina (Figure S3). The total number of microglia sequenced from n=10 mice was 4,949 (olfactory bulbs), 4,563 (cortex), 6,855 (hippocampus), 5,724 (cerebellum) and 5,004 (retina). Cells were sequenced to a depth of ∼63,000-88,000 reads/cell and similar median numbers of genes per cell and median unique molecular identifiers (UMI) per cell were detected in each of the samples (Supplementary file 3).

In the first round of analysis, cells from each of the investigated CNS regions were pooled into a single dataset (aggregated data: 27,095 total cells). For the purpose of defining what constitutes a microglia cell in aggregated data, the expression of *Tmem119* and *P2ry12* was examined (Figure 3a) and cells that did not express *Tmem119* and *P2ry12* were filtered out and excluded from subsequent analyses. The majority of cells in the aggregated data (94%) expressed the microglia specific genes *Tmem119* and *P2ry12*, supporting the accuracy of the microglia enrichment method used in this study. When visualising the aggregated data according to tissue origin, retinal microglia had limited overlap with brain microglia, and a large portion of cerebellar microglia clustered separately from olfactory bulb, cortical and hippocampal microglia (Figure 3b). These findings are consistent with our bulk RNA-seq data findings and further demonstrate that microglia gene expression is region-dependent in the healthy CNS.

**Figure 3.**
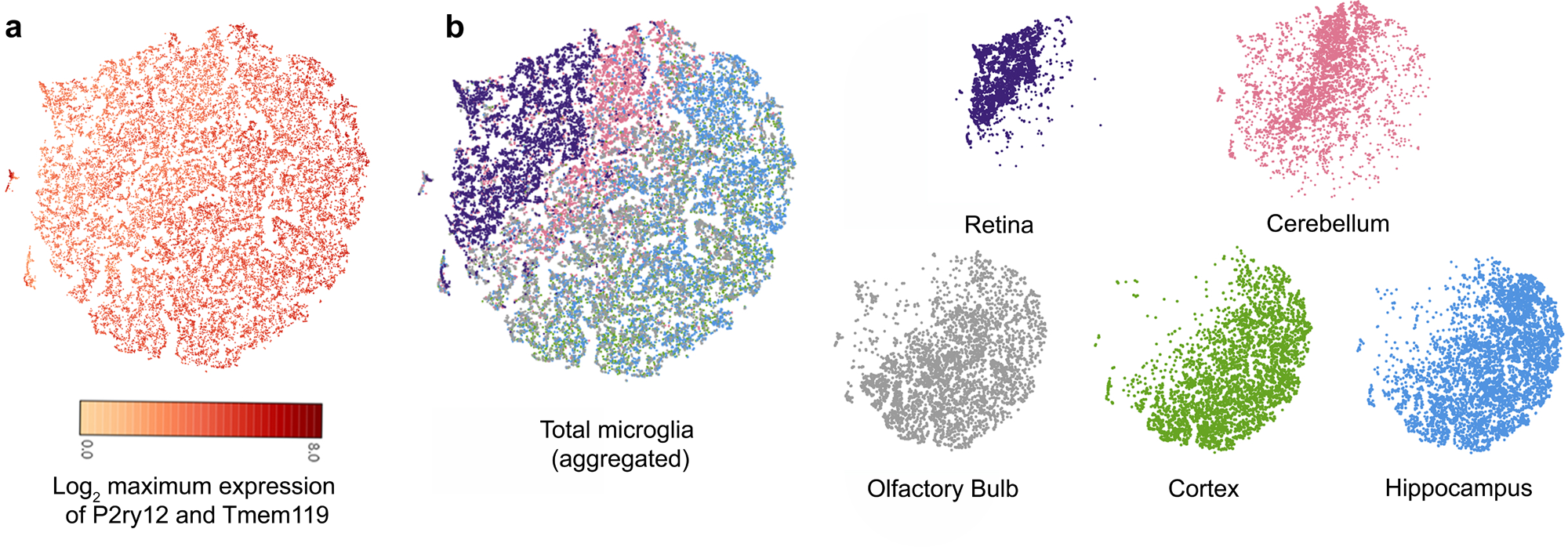
Single cell RNA-sequencing of microglia from different regions of the healthy adult mouse CNS. **(a)** tSNE plot of 27,095 cells aggregated from all five CNS regions, showing expression of the microglia homeostatic genes *Tmem119* and *P2ry12.* **(b)** tSNE plots of microglia, coloured according to tissue region of origin. A total of 25,465 microglia were analysed after removing cells that did not express *Tmem119* and *P2ry12*.

### Characterization of microglia clusters identified within CNS regions by scRNA sequencing

To further characterise the transcriptomic heterogeneity of microglia within different neuroanatomical regions of the healthy mouse CNS, dimensionality reduction and unsupervised clustering were performed separately on cells from each CNS region. After exclusion of cells that did not express *Tmem119* and *P2ry12*, the expression of selected microglia (Figure S4) and myeloid (Figure S5) signature genes was examined. As expected, the signature microglial genes (*Tmem119*, *P2ry12, Fcrls*, *Slc2a5*) and myeloid genes (*Cx3Cr1*, *CD45*, *Itgam*, and *Aif1*) were highly expressed by most of the analysed cells. Whilst *P2ry12* and *Tmem119* were uniformly expressed in microglial cells isolated from each CNS region, *Fcrls* was expressed by fewer retinal microglia compared to brain microglia. In addition, fewer cells expressed *Slc2a5* in all CNS regions compared with *P2ry12*, *Tmem119* and *Fcrls* (Figure S4).

Several microglia clusters were identified within each examined region of the mouse CNS: olfactory bulb: 6 clusters, cortex: 5 clusters, hippocampus: 6 clusters, cerebellum: 5 clusters, and retina: 6 clusters. In line with previous studies (14, 34), a large portion (66.5%-94.8%) of total microglia within each region were identified as being homeostatic microglia. Homeostatic microglia spanned several clusters within each CNS region (Figure 4a-e) that were not defined by any unique genes. However, unique microglia clusters with distinct gene expression signatures were also identified, including interferon-enriched microglia (IFN), chemokine-enriched microglia (CC), apolipoprotein-enriched microglia (APO) (Figure 4a-e). These clusters did not have significant enrichment of immediate early genes (*Fos*, *Jun*) or stress-induced genes (*Hspa1a*, *Dusp1*) relative to other clusters (Figure S6), suggesting they were unlikely an artefact as a result of *ex vivo* activation (35). The IFN cluster was observed in each CNS region, and this cluster was characterised by increased transcript levels of interferon-stimulated gene family members including *Ifit1*, *Ifit2*, *Ifit3*, *Ifit3b*, *Ifitm3*, *Oasl1*, *Oasl2*, *Ifi204*, *Ifi206*, *Ifi207*, *Ifi209*, *Ifi211*, *Ifi213* and *Ifi2014*. This cluster represented 0.8-3.1% of microglia within the examined brain regions and was found at the highest frequency within the retina (9.2% of total retinal microglia). A small proportion of cells within the retinal IFN cluster also expressed *Ccl3* and *Ccl4*, whereas a separate CC cluster characterised by unique expression of *Ccl2*, *Ccl3*, *Ccl4*, *Ccl6* and *Ccl9* and increased expression of *CD9*, *CD63*, *CD83*, *Ctsb*, *Ctsz* was identified in each of the brain regions examined (3.6-9.5% of total microglia within each region). This separate CC cluster was not present in retinal microglia. Interestingly, a unique APO microglia cluster (representing 19.5% of total microglia) was identified within the olfactory bulb (but not within other CNS regions). Genes enriched in the APO cluster included *Apoe* and *Lyz2*. Furthermore, cluster 5 within the olfactory bulb was a unique cluster restricted to olfactory bulb (1.17% of olfactory bulb microglia) and featured a small percentage of cells enriched in genes related to cellular proliferation, including *Mki67*, *Prc1*, *Top2a*, *Birc5, Hist1h1b*, *Knl1*, *Ckap* and *Cdca2*. Retina C5 microglial cluster, although relatively small in abundance, was characterized by elevated expression of lysosomal and antigen-processing genes, including *Lgmn*, *Psap*, *Hexb*, *Cd81*, *Cd37*, *Ctsd*, Ctss, *Ctsl*, and *Ctsz*. The transcriptional profile suggests a function specialized in phagocytosis and degradation of cellular debris, as well as antigen processing and immune surveillance.

**Figure 4.**
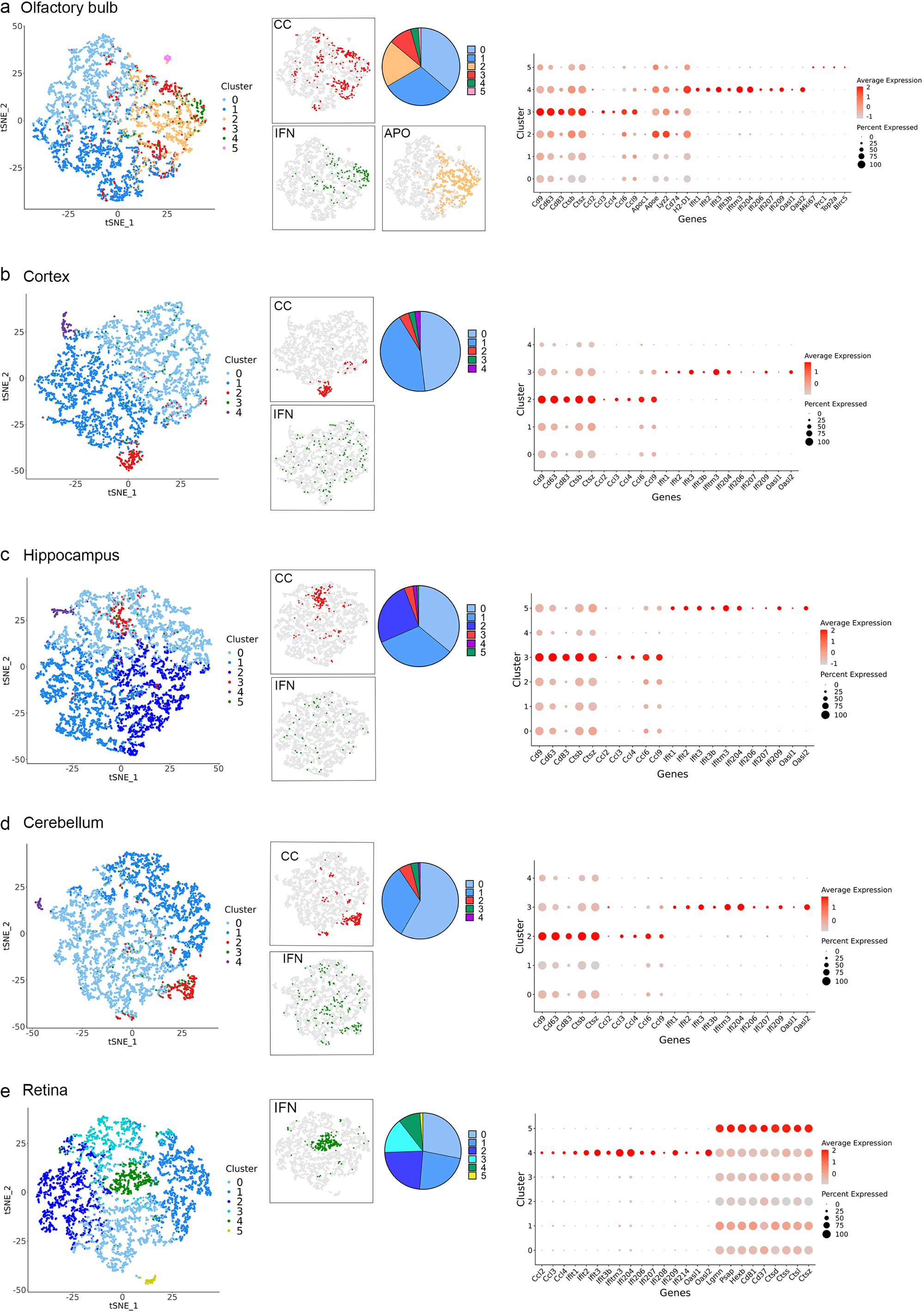
Identification of microglia clusters in different neuroanatomical regions of the healthy adult mouse CNS. **(a) – (e)** Identification of microglia clusters within the olfactory bulb (4,600 cells), cortex (4,341), hippocampus (6,539 cells), cerebellum (5,276 cells), and retina (4,709 cells) of 8-week-old female C57B6/6J mice. Left: tSNE plots showing microglia clusters within each region. Homeostatic microglia clusters: shades of blue and purple; chemokine-enriched microglia (CC): red; interferon-enriched microglia (IFN): green; apolipoprotein-enriched microglia (APO): light orange; C5 in olfactory bulb: pink; and C5 in retina: yellow. The CC, IFN and APO microglia clusters are displayed in individual tSNE plots (grey cells represent microglia not belonging to the specific cluster displayed). Pie charts showing the proportions of different clusters within each region. Right: Dot plots showing the expression of genes across microglia clusters, with the dot size representing the percentage of cells expressing the gene and the colour representing its average expression within a cluster.

These findings confirm that homeostatic microglia make up the majority of microglia within the healthy adult mouse CNS and additionally provide evidence of smaller diverse microglia populations in the brain and retina. The regional comparison of clusters provides new insights into the distribution of microglia clusters within anatomically distinct tissue regions.

### Microglia clusters: in situ validation

To investigate whether the IFN and CC clusters revealed by scRNAseq were relevant to microglial phenotypes displayed *in situ*, immunohistochemistry was performed on fixed frozen brain tissue sections from Cx3cr1-gfp/+ mice, using CCL4 and IFIT1 antibodies to identify interferon-enriched microglia and chemokine-enriched microglia respectively. Small numbers of Cx3cr1-GFP+ microglia expressed CCL4 (Figure 5) and IFIT1 (Figure 5). These observations are consistent with the data obtained by scRNA-seq and provide evidence for a CC microglia cluster (CCL4+) and an IFN microglia cluster (IFIT1+) within healthy the adult mouse CNS.

**Figure 5.**
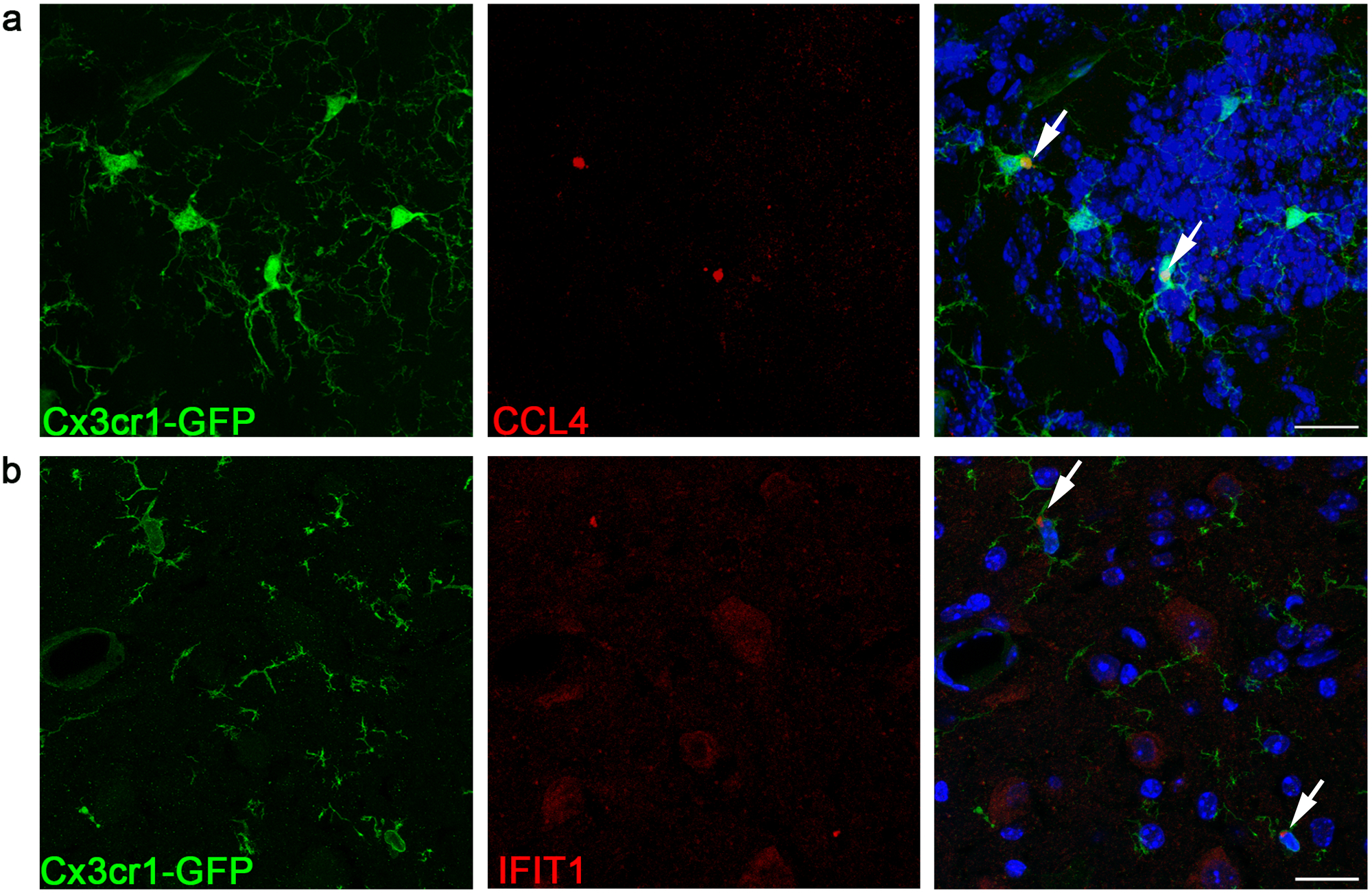
Identification of CCL4 and IFIT1 expressing microglia within healthy mouse brain using immunofluorescent staining and confocal microscopy. Immunostaining panels display co-expression of CCL4 **(a)** or IFIT1 **(b)** (red) and Cx3Cr1-GFP (green). Scale bars represent 20 µm.

### Predicted functions of identified microglia clusters

To study how the different transcriptomic profile of microglia clusters may relate to their functions, the DEGs were used to create gene ontology networks. DEGs common to the CC cluster within each region were related to functions including ‘cell motility and chemotaxis’, ‘regulation of cytokine production’, ‘regulation of vasculature development’, ‘metabolism’, ‘regulation of apoptotic process’, ‘response to external stimulus and ERK1/2 cascade’ (Figure 6a). When comparing the DEGs of CC clusters between different CNS regions some differences were observed. For example, the GO terms ‘protein folding and response to unfold protein’, and ‘leukocyte differentiation’ were identified in the CC cluster within the olfactory bulb, but not for other regions (Figure S7). DEGs shared by IFN clusters within each CNS region were linked to the GO terms ‘response and defence to virus and symbiont’, ‘regulation of innate immune response’, ‘response to molecule of bacterial origin and lipopolysaccharide’, ‘cellular response to interferon type I, alpha, beta and gamma’, ‘antigen processing and presentation’, ‘cell killing’, ‘regulation of response to cytokine stimulus and immune response’ (Figure 6b). The DEGs and associated GO terms for the IFN cluster were similar between CNS regions (Figure S8). The unique APO cluster identified in the olfactory bulb had DEGs that were involved in protein-lipid metabolism, bacteriolytic function and antigen processing and presentation particularly via MHC class II pathways (Figure 6c). The other unique olfactory bulb cluster (cluster 5) had enrichment of genes related to ribosome biogenesis and assembly, regulation of protein polymerization and actin filament and lamellipodium organization and metabolic process (Figure 6d).

**Figure 6.**
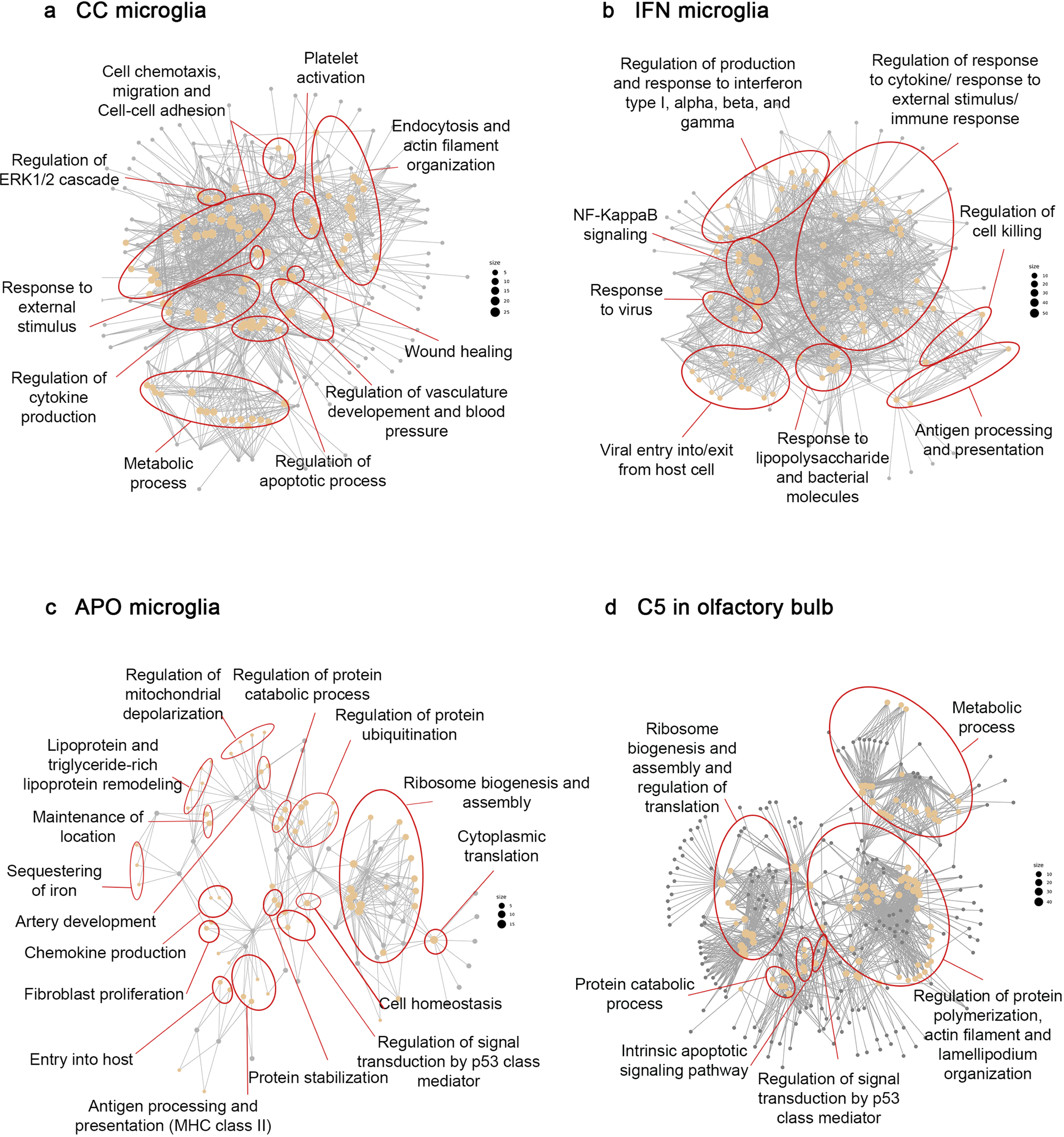
Gene ontology network of top 100 GO terms based on genes that are commonly differentially expressed in CC, IFN, Apo clusters in each CNS region, and in unique OB cluster 5. **(a-d)** Each node represents a gene ontology and the main themes within the data group in clusters. GO networks were generated from DEGs from all five CNS regions.

### IFN microglia signatures are conserved in human retinal microglia

One important observation from our analysis of healthy mouse microglia was the distinct transcriptional profile of retinal microglia compared to brain microglia. To assess whether similar region-specific features were conserved in the human retina, we performed single-cell RNA-sequencing on enriched human retinal microglia obtained post-mortem from a 45-year-old male tissue donor. This analysis identified microglia based on the expression of canonical myeloid markers, including *AIF1*, *PTPRC, ITGAM, C1QA*, *GPR34*, and *SLC2A5* (Figure 7a). A small population of natural killer cells (NK) and other retinal cells were also identified (Figure 7b and 7c). Microglia exhibited heterogeneous gene expression signatures and were classified into four subclusters (MG0–MG2 and MG4) (Figure 7b and 7c). Notably, IFN-stimulated genes such as *IFITM2* and *IFITM3* were among the enriched transcripts (particularly for cluster MG0 and MG4), suggesting the presence of IFN-responsive microglial subtypes in the human retina, similar to the mouse retina, which also contained a prominent IFN-responsive cluster with comparable gene expression patterns.

**Figure 7.**
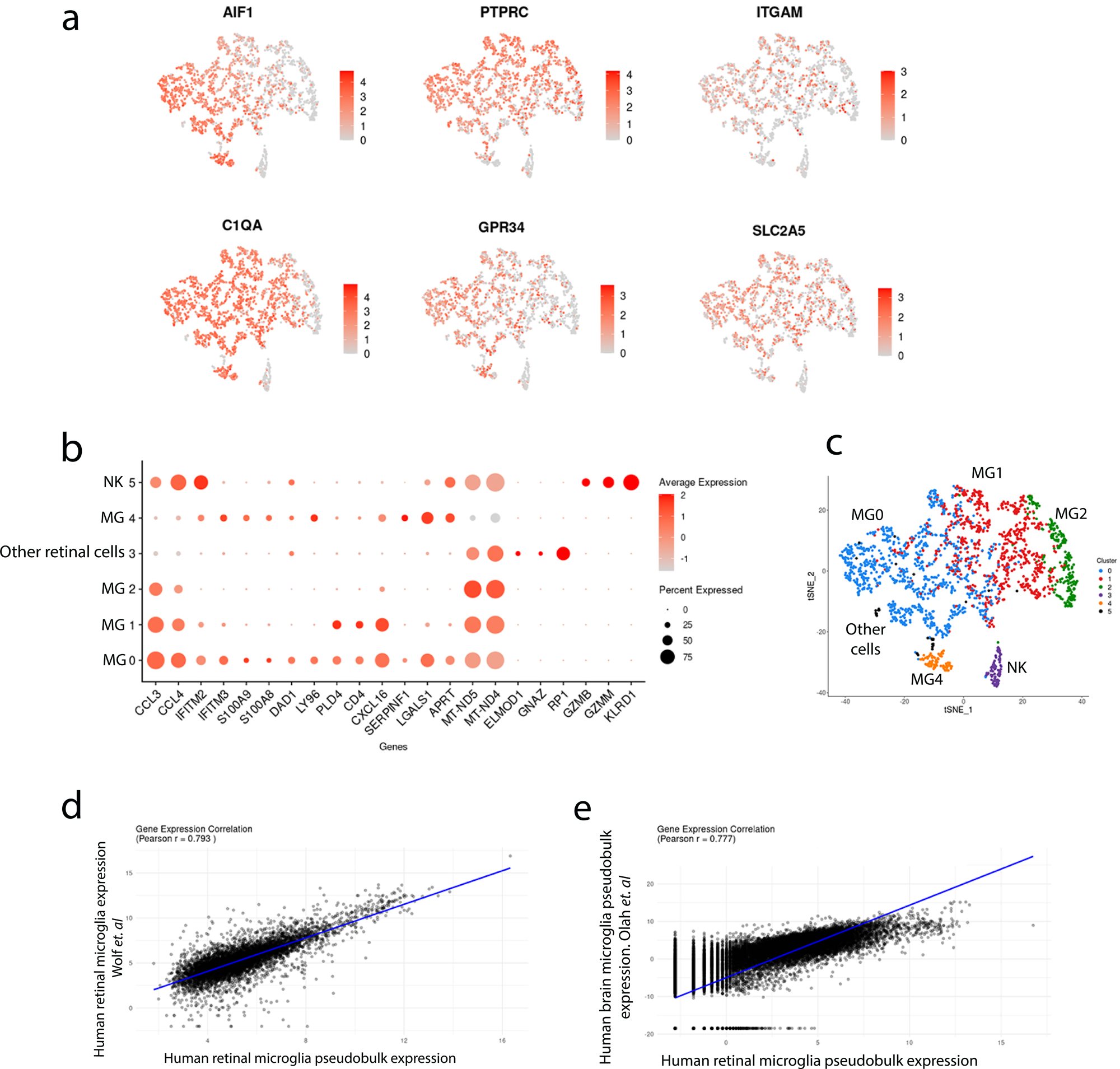
Characterisation of microglial populations in the human retina using single-cell RNA sequencing. **(a)** tSNE plots show the expression of canonical microglial and myeloid markers (*AIF1*, *PTPRC*, *ITGAM*, *C1QA*, *GPR34*, and *SLC2A5*) across sequenced human retinal cells (2,089 cells and 249,622 mean reads per cell). **(b)** Dot plot displays the average expression and percentage of cells expressing key DEGs across identified microglial subtypes (MG0–MG2 and MG4) and other retinal and immune cell types. **(c)** Correlation between human retinal microglia scRNA-seq-derived pseudobulk (this study) and bulk RNA-seq of human retinal microglia (36) shows high concordance (r= 0.79, *p* < 0.001). **(d)** Pseudobulk expression correlation between human retinal microglia scRNA-seq (this study) and a human brain microglia scRNA-seq dataset (37) reveals a strong positive correlation (r = 0.77, *p* < 0.001).

To assess the reliability of the human scRNA-seq data, we performed a pseudobulk expression analysis by aggregating gene expression profiles from human retinal microglia. This psuedobulk profile showed strong correlation with bulk RNA-seq data from isolated human retinal microglia reported by Wolf *et al.* (36) (r = 0.79, P < 0.001) (Figure 7d), validating the robustness of our single-cell dataset. Additionally, a similarly strong correlation was observed between pseudobulk expression of human retinal microglia (this study) and that of human brain microglia reported by Olah *et al.* (37) (r = 0.77, P < 0.001). Taken together, pseudobulk analysis supports the presence of conserved core microglial programs, while also highlighting distinct, retina-specific transcriptional features that may reflect tissue-specific adaptations (Figure 7e).

## Discussion

This study used transcriptomic approaches to investigate the regional and sub-regional heterogeneity of microglia in the steady state mouse CNS. Bulk RNA-seq demonstrated region-dependent microglial transcriptomic profiles, which aligns with recent research highlighting the impact of the CNS microenvironment on shaping microglial function, phenotype, immune response, and gene expression (21–24, 27, 38, 39).

Gene ontology analysis revealed that hippocampal microglia were the most similar to cortical microglia. The small number of DEGs identified in hippocampal microglia were mainly involved in ‘cellular process’, ‘biological regulation’, and ‘metabolic process’, which is consistent with Grabert *et al.*’s findings of higher energy production gene expression in hippocampal microglia (21). Previous studies have suggested that microglial responses to injury and pathological conditions vary according to neuroaxis locations (40, 41), although our data demonstrate higher complexity in the distribution of microglia immune states in the healthy CNS than previously thought. In this study, olfactory bulb and cerebellar microglia exhibited enrichment in ‘immune system processes,’ suggesting a more vigilant immune state in these brain regions compared to cortical microglia. One possible explanation for this is that the olfactory bulbs and cerebellum are more vulnerable to invasion from neurotropic pathogens due to their proximity to the olfactory and trigeminal nerves respectively. These nerves serve as a conduit for infectious agents to travel from the nasal cavity directly into the brain (42–44), hence microglia in the olfactory bulbs and cerebellum may exhibit heightened immune responsiveness as potential first responders. In support of this hypothesis, Abellanas *et al*. reported that midbrain microglia - located near the pons where the trigeminal nerve originates in the brainstem – exhibit enhanced expression of immune-related genes compared to striatal microglia, which reside in the forebrain and are anatomically rostral to the trigeminal nerve, in the adult steady state mouse brain (45, 46).

Our study also revealed significant differences in the transcriptome of retinal and brain microglia, highlighting enrichment of genes related to phagocytosis, antigen processing and presentation, and chemokines in retinal microglia when compared to cortical microglia. Notably, the phagocytosis-related genes *Tulp1* and *Axl* exhibited a log_2_ FC of 8.7 and 4.3 respectively in retinal microglia. These genes have known roles in apoptotic cell clearance and immune regulation (47–49). Additionally, *Axl*, one of the transcriptional signatures of DAM (50), is thought to play a dominant role in DAM activation in the retina (51). The unique gene profile of retinal microglia also included increased expression of MHC class I-related genes and chemokines including *Cxcl13*, *Cxcl10* and *Ccl5*, suggesting a specialized role in immune vigilance and viral defence in the retina (52, 53). This is an interesting finding, given that the environment of both the brain parenchyma and neural retina are regulated by the blood-brain and blood-retina barriers respectively. Our findings, coupled with the susceptibility of the retina to viral infections caused by Flaviviridae (54), Herpesviridae (55) and SARS-Cov-2 (56, 57), underscores the apparent readiness of retinal microglia to respond to viruses, warranting additional studies on their function during infection.

As microglia in the retina, cerebellum, and olfactory bulbs exhibited enhanced responsiveness to danger signals compared to cortical and hippocampal microglia, we next asked whether this immune-responsive profile was consistent across all microglia within these regions or limited to specific subtypes. To address this, we used single-cell RNA sequencing to profile microglia clusters within each CNS region. Consistent with previous studies by Hammond *et al.* and Marsh *et al.* (14, 34), the majority of microglia with the steady state adult mouse olfactory bulbs, cortex, hippocampus, cerebellum and retina displayed the classical homeostatic microglia transcriptomic signature. However, we additionally identified small populations of unique microglia clusters, including CC (chemokine-enriched), IFN (interferon-enriched) and APO (apolipoprotein-enriched) clusters.

The identification of a CC microglia cluster in each of the examined brain regions aligns with a previous study noting a small population of *Ccl4*-expressing microglia in the mouse brain, peaking during development and expanding in aging and injury (14). The chemokines expressed by this cluster are known to contribute to the pathogenesis of neuroinflammation by modifying BBB function (58), regulating the entry of peripheral immune cells into the brain (59) and recruiting immune cells to injured brain areas (60). The CC cluster also showed enrichment of genes associated with the regulation of vasculature development, which highlights the potential role of these cells in the proliferation, migration, and activation of endothelial cells. Although we did not investigate the interaction of CC microglia with blood vessels in this study, CC microglia are likely distinct from capillary-associated microglia described by Bisht *et al.* (61), as they comprise a much smaller proportion of total microglia. While additional work is required to determine the role of the CC cluster, we propose that these cells are a specialised subtype of microglia that are equipped to respond to external stimuli or pathological conditions by regulation of energy, chemokine production and angiogenesis.

An IFN responsive microglia cluster, characterised by upregulation of interferon-stimulated genes including members of *Ifit* and *Ifitm* gene families, was identified in all examined brain regions and the retina. Proteins encoded by interferon-stimulated genes (62, 63) are antiviral proteins (64), which are induced by IFN production from virus-infected cells. The *Ifitm3* gene, which is highly upregulated in the IFN cluster, is a component of the microglial “sensome” that functions to sense pathogen-associated molecular patterns and danger-associated molecular patterns (65). In addition, type I IFN signalling pathways are involved in regulation of microglia function during development and neuroinflammation (66), and alter microglial phenotype and function, including phagocytosis (67). The phagocytic role of a type I IFN responsive microglia subset was previously identified during topographic remapping, and in the pathological conditions such as AD and neurotropic viral infection (68). Based on these data, the IFN cluster identified in this study may be a specialised microglia subtype involved in antiviral responses and phagocytic function through and regulation of production and response to type I IFNs and the microglia sensome.

In the present study, the IFN cluster constituted 9.3% of microglia within the mouse retina while in the brain regions this population formed 0.8-3.1% of the total microglia population. This difference in frequency of the IFN cluster between brain and retinal microglia may explain why retinal microglia appeared to be distinct from brain microglia in our bulk RNA-seq analysis. The reason for the increased frequency of IFN responsive microglia within the retina is unclear; however, an explanation could be that the physiological and neurological microenvironment of the brain and neural retina differ in blood tissue barrier permeability (69), cellular composition, and neurotransmitter profiles (70–73). Another theory is that distribution and expression characteristics of endogenous retroviruses and other retrotransposons may be different in CNS regions (21, 74). Although these endogenous retroviruses are largely non-functional and inactive (75), stress signals in response to injury and infection activate certain endogenous retroviruses and virion production in a cell type specific manner (76). Recent studies have suggested that gene products of endogenous retroviruses and retrotransposons participate in various pathologic processes (77, 78). Therefore, it is possible that microglia in these CNS regions may express antiviral elements to maintain tissue homeostasis, although this requires more investigation and further studies are required to determine how endogenous retroviruses are differentially expressed throughout the CNS and if they play a role in shaping microglia phenotype and function.

Another interesting finding of the present study is that 33.5% of all olfactory bulb microglia (comprising CC, IFN, Apo, and OB cluster 5) had unique gene expression profiles distinct from homeostatic microglia. These findings are in line with previous reports demonstrating unique functions of olfactory bulb microglia (79, 80). Two microglia subtypes (Apo cluster and OB cluster 5) were uniquely found in the olfactory bulbs. The Apo cluster was enriched in genes involved in protein-lipid metabolism, which has been shown to be involved in microglia activation and phagocytosis (81). This suggests that the Apo cluster may be specialised for sensing and processing lipid antigens. The Apo cluster showed similarity with ‘Stage 1 DAM’ reactive microglia (characterised by the expression of a set of genes including *ApoE*, *Lyz2*, *Lpl*, *Ctsb*, *Fth1*, and *Ctsl*) that have been described in pathological conditions (17). Furthermore, upregulation of antigen processing and presentation genes (both MHC class I and MHC class II related genes) such as *H2-D1*, and *Cd74* in this cluster suggested the importance of an antigen presenting role in olfactory bulb microglia. This is consistent with our bulk RNA sequencing data, which demonstrated higher expression of MHC class II genes in olfactory bulb and cerebellar microglia compared to those in the cortex, hippocampus, and retina. Finally, the olfactory bulb also contained a unique microglia cluster (OB cluster 5), which was characterised by upregulation of genes associated with cellular proliferation. Previously, enrichment of proliferation pathways were found in developmental stages while only a small percentage of microglia after age P30 were demonstrated to be proliferative cells in the healthy mouse whole brain (14). Our study found that these cells are localised to the olfactory bulb. The spatial restriction of the Apo cluster and OB cluster 5, along with presence of the CC and IFN clusters in olfactory bulb, suggests that olfactory bulb microglia are more likely to monitor, and prime immune responses compared to microglia located in other regions of the brain, although further functional studies need to fully understand the role of these populations.

## Conclusion

By systematically comparing microglial transcriptomes across distinct neuroanatomical regions of the mouse CNS, we demonstrated that microglia exhibit both shared and region-specific gene expression profiles, reflecting local environmental cues and functional specialisation. These regional differences may be in part due to unique clusters of microglia that differed in distribution and density throughout the steady state CNS. We propose that the non-homeostatic microglia clusters may expand during neuroinflammation and drive local immune responses. Notably, the enrichment of IFN-responsive microglia in the retina and their presence in both mouse and human tissue suggests that conserved innate immune subtypes may contribute to neuroinflammatory processes. This study advances our understanding of microglial diversity across the CNS and provides a foundation for investigating region-specific microglial functions in disease contexts.

## Resource availability

The sequencing data presented in this publication have been deposited in NCBI’s Sequence Read Archive database (accession number PRJNA1345092). Differential gene expression data generated by bulk RNA-sequencing are included in the supplementary files of this publication.

## Materials and Methods

The manufacturers and catalogue numbers of the antibodies, chemicals and consumables used in this study are provided in the accompanying Key Resources Table.

### Ethics and biosafety approvals

Animal ethics approval for this study was obtained from QIMR Berghofer Medical Research Institute Animal Research Ethics Committee (approval A18613M); Monash Animal Research Platform (approval MARP-2018-017); QUT AEC administrative approval 1800001261. A notifiable low risk dealing for the use of genetically modified mice in this project was obtained from the QUT University Biosafety Committee (NLRD approval 1800000957). All animal experiments were performed in accordance with the Association for Research in Vision and Ophthalmology Statement for the Use of Animals in Ophthalmic and Vision Research and the National Health and Medical Research Council Australian Code for the Care and Use of Animals for Scientific Purposes. Human ethics approval was obtained from QUT HREC (ERM Project 5287) and Queensland Health, Metro South Hospital and Health Service (ERM Project 9461).

### Mice and tissue collection

Adult (6-16 weeks of age) female C57Bl/6J and Cx3cr1^gfp/+^ mice were used for this study. C57Bl/6J mice were purchased from the Animal Resources Centre (Canning Vale, Western Australia). Cx3cr1^gfp/+^ mice were generated by cross breeding Cx3cr1^gfp/gfp^ mice obtained from the QIMR Berghofer Medical Research Institute Animal Facility with C57Bl/6J wild type mice. Mice were maintained on a 12:12 hour light cycle with access to food and water ad libitum. Mice were deeply anaesthetised by an intraperitoneal injection of sodium pentobarbital (150 mg/kg). Mice were then perfused immediately through the left ventricle with cold phosphate buffered saline (PBS) to clear the vasculature of blood and circulating leukocytes. When tissues were being collected for histology and immunostaining, mice were also transcardially perfused with 4% paraformaldehyde (PFA) to deliver fixative systemically to the animal. Brains were collected and microdissected to obtain cerebral cortex, olfactory bulbs, hippocampus and cerebellum. The neural retina removed from the underlying RPE, choroid, and sclera (82).

### Mouse brain cryotomy and eye dissection

Brains and eyes were collected from perfusion fixed Cx3cr1^gfp/+^ mice and stored in 4% PFA at 4°C overnight. Retinae were dissected from the posterior eye cup and flat-mount preparations for wholemount immunostaining were prepared by creating radial incisions. Fixed brains were cryoprotected in 10% (w/v) sucrose in PBS for 48 h at 4° C, then 30% (w/v) sucrose in PBS for 48 h at 4° C. Cryoprotected brains were embedded in Tissue-Tek Optimal Cutting Temperature medium and rapidly frozen in chilled isopentane. Frozen brains were stored at −80 °C until the time of cryosectioning. Free-floating, 40 µm coronal brain sections were cut on a Cryostar NX70 cryostat (ThermoFisher) and stored in cryoprotectant solution (0.05 M phosphate buffer, 30% (w/v) sucrose, 1% (w/v) polyvinylpyrrolidone, 30% (v/v) ethylene glycol) in multi-well plates at −20 °C. Brains were sectioned in a rostral to caudal orientation, and every section was collected.

### Bulk RNA sequencing of mouse microglia and differential gene expression analysis

Adult female C57Bl/6J mice (8 weeks of age) were used for bulk RNA sequencing (n=20 mice in total). Following euthanasia, mice were transcardially perfused with ice cold PBS and brains and eyes were collected. Tissues were micro-dissected to obtain olfactory bulbs, cortex, hippocampus, cerebellum, and neural retina. The dissected tissues were pooled into 4 biological replicates (each replicate consisting of pooled tissue from five mice). The dissected tissues were transferred to 50 mL tubes containing dissection buffer (1x HBSS, no calcium, no magnesium, no phenol red; 5 mM glucose; 15 mM HEPES) and all processing was performed on ice. The tissues were passed through a 70 μm nylon cell strainer using the plunger of a 5 mL syringe to obtain single cell suspensions, and then pelleted by centrifugation at 400 *g* for 5 min at 4 °C. Brain tissue was resuspended in 30% (v/v) Percoll and then centrifuged at 700 *g* for 10 min at 4 °C without a brake being applied. The myelinated (i.e., hydrophobic) layer formed at the top of the tube was discarded prior to resuspending the cell pellet. The number of viable cells was determined by exclusion of Trypan blue stain as viewed by microscopy, assisted by use of a hemocytometer. Cells were subsequently resuspended in FACS buffer (3 mM EDTA, 0.1% (w/v) BSA, 1 x PBS, 100 μg/mL DNase I) and stained with fluorescent-conjugated antibodies (rat anti-mouse CD45-BUV395, and rat anti-mouse CD11b-APC-Cy7) for 30 min on ice, then centrifuged and resuspended in FACS buffer. Microglia (CD45^int^ CD11b^high^ cells) were isolated from stained cell suspensions using a BD Influx cell sorter (100 µm nozzle) at Monash FlowCore. Dead cells were identified and excluded from analysis by staining with propidium iodide (PI, 1 μg/ mL). RNA was isolated from sorted microglia using the miRNeasy Micro Kit (Qiagen 217084, as per manufacturer’s instructions) and stored at −80 °C. RNA was sent to Monash Health Translation Precinct Medical Genomics Facility for quality control, cDNA library construction and ultra-low input RNA-seq using paired-end sequencing (Illumina NextSeq550 V2 High output).

To identify differentially expressed genes between microglia isolated from different CNS regions, data QC and differential gene expression analysis was performed. Raw fastq. files were analysed using the RNAsik pipeline (83) and reference data was obtained from Ensembl. Degust (84) was used to perform differential gene expression analysis, using Limma-voom for statistical testing. Cortical microglia were set as the reference to which all other microglia datasets were compared for differential gene expression analysis. The bulk RNA-seq dataset was filtered for the differentially expressed genes with a false discovery rate (FDR) <0.05 and a log_2_ fold change of >1 compared to cortical microglia. Heatmaps were generated using Heatmapper (85). Pathway Analysis was performed using InnateDB and Panther (Protein ANalysis THrough Evolutionary Relationships) Classification System database to classify genes and determine biological pathways that are significantly different between microglia populations. Genes were classified according to biological process. The Uniprot database was used to determine gene molecular and biological function, family, and localisation of protein in the cell.

### Single cell RNA sequencing of mouse microglia

Microglia were isolated from the olfactory bulbs, cortex, hippocampus, cerebellum, and retinae of C57Bl/6J female mice (6 weeks of age). Freshly dissected tissues were pooled from n = 10 mice to obtain sufficient cell numbers. The dissected tissues were processed as described for bulk RNA-seq above. Cells were stained with a panel of fluorescent-conjugated antibodies: rat anti-mouse CD45-BV421, rat anti-mouse CD11b-APC-Cy7 and mouse anti-mouse Cx3cr1-FITC. Dead cells were excluded by PI staining (1 μg/mL). Microglia (CD45^int^ CD11b^high^ Cx3cr1^high^ cells) were sorted using a BD FACS Aria IIIu (100 µm nozzle) at QIMR Berghofer Flow Cytometry Facility. Sorted cells were encapsulated into nanoliter-scale droplets using the Chromium Controller (10x Genomics) and scRNA-seq libraries were prepared using the Chromium Single Cell 3′ Reagent Kit v3.1. Libraries were sequenced on an Illumina NovaSeq S1 platform (paired-end reads) at the University of Queensland Sequencing Facility.

### Mouse scRNAseq data analysis: Processing, QC metrics, removing non-target cells and clustering

Sequencing data were processed following the analytical pipeline described in our previous study of choroidal and meningeal immune cells (86), with adaptations for microglia. Raw base call files were demultiplexed using Cell Ranger v7.0.0 (10x Genomics) (87) via cellranger mkfastq, and gene expression quantification was performed using cellranger count against the mouse reference genome. For samples processed separately, outputs were aggregated using cellranger aggr, and gene expression levels were normalized based on relative library sizes. Gene-barcode matrices were imported into R v4.0.5 (88) using the Read10X function from the Seurat package (89). Data were converted into a Seurat object using CreateSeuratObject, and standard pre-processing steps were applied: log-normalization, identification of highly variable features, scaling, and dimensionality reduction via PCA. Clustering was performed using the Louvain algorithm (FindClusters), and data were visualized using UMAP and t-SNE. Quality control included filtering of cells with fewer than 200 or more than 6000 detected genes. Additionally, non-target cells were excluded based on the absence of homeostatic microglia genes (*Tmem119*, *P2ry12*). Clustering resolution was optimized using the Clustree package (90) to avoid over- or under-clustering. Once quality filtration was complete, cells were clustered by gene expression using Seurat’s ‘FindClusters’ function and clustering visualised with t-SNE, dot plots and heat maps.

### Mouse scRNAseq data Analysis: Identifying DEGs, cell subtypes, and functional enrichment

Differentially expressed genes (DEGs) between clusters were identified using Seurat’s FindMarkers function with thresholds of adjusted *p* < 0.05 and log fold change > 0.5. Cluster identities were assigned based on expression of known microglia subtype markers and functional annotations. Functional enrichment analysis for Gene Ontology (GO) terms was conducted using the clusterProfiler package (91). Gene identifier conversion (e.g., to Entrez or Ensembl IDs) was performed using the AnnotationHub package (92). Microglial clusters were characterized by both marker gene expression and enriched biological pathways. Homeostatic microglia were defined by expression of *Tmem119* and *P2ry12* and minimal transcriptional divergence (<6 DEGs) from the other microglial populations.

### Immunostaining, confocal microscopy and image analysis

Wholemount retinae and free-floating brain sections (40 µm) were permeabilised in 0.5% (v/v) Triton x-100 (Sigma Aldrich X100-100ML) in PBS at room temperature for 1 h. The tissues were then blocked in 3.0% (w/v) bovine serum albumin (Sigma) and 0.3% (v/v) Triton X-100 in PBS for 1 h at room temperature. Sections were incubated with primary antibodies (Key Resources Table) overnight at 4 °C. Tissues were washed three times (each 10 min) in PBS, and then incubated with fluorophore-labelled secondary antibodies (Key Resources Table) and Hoechst (20mM) for 2 h at room temperature. Tissues were washed three times (each 10 minutes) in PBS, mounted on microscope and coverslipped. Samples were imaged using an inverted SP5 5-channel confocal microscope (Leica Microsystems) using a 40X objective (numerical aperture 1.25). Z stacks were captured every 1 µm for brain sections and every 2 µm for retina, with a line averaging value of 3 applied. Retinal wholemount Z stacks were captured from the Ganglion Cell Layer (GCL) to the Outer Nuclear Layer (ONL). Brain regions for imaging were identified using the Mouse Brain in Stereotaxic Coordinates Atlas (93). Maximum intensity projection images were created by using FIJI (94).

Microglial density was calculated by counting the number of microglia in three random fields of view for each CNS region per mouse and dividing the number of counted cells by the total field of view area to obtain the number of microglia/mm^2^. To compare the morphometry of microglia in different regions of the mouse CNS, Sholl analysis was performed to determine microglia process complexity (calculated by area under the curve analysis of the number of branch intersections at increasing radial distances from the cell body), ramification index and cell body area as described previously (95–98). This morphometric analysis was performed on six microglia from each CNS region per mouse.

### scRNA-seq of human retinal microglia

Single-cell RNA sequencing was performed on cells isolated from human postmortem eyes. Fresh (unfixed) eye cups from a 45-year-old male donor with no history of eye disease or diabetes were obtained within 7 hours postmortem from the Queensland Eye Bank, Australia. The retinas were dissected and transferred to a 12-well plate for enzymatic digestion with Collagenase D (2.5 mg/mL in HBSS containing calcium and magnesium, without phenol red). Tissue was incubated at 37 °C for 15 minutes, followed by gentle trituration using a pipette and filtration through a 70 µm cell strainer to obtain a single-cell suspension. Cells were washed and resuspended in CryoStor CS10 freezing medium and placed in a Mr Frosty container to undergo controlled-rate freezing at −80 °C overnight prior to transfer to liquid nitrogen storage.

For scRNA-seq, frozen retinal cells were rapidly thawed and resuspended dropwise in pre-warmed DMEM/F12 containing 10% (w/v) BSA. Cell viability and yield were assessed using trypan blue staining and a hemocytometer. Thawed cell suspensions were incubated with anti-human Fc block for 15 minutes on ice, centrifuged at 400 × g for 5 minutes, and resuspended in FACS buffer containing BV421-conjugated mouse anti-human CD45 antibodies. After 30 minutes of incubation on ice, cells were centrifuged again and resuspended in FACS buffer for sorting. Live microglia (PI^neg^ CD45^int^) were sorted using a BD FACS Aria IIIu (100 µm nozzle). Sorted cells were encapsulated into nanoliter-scale droplets using the Chromium Controller (10x Genomics) and scRNA-seq libraries were prepared using the Chromium Single Cell 3′ Reagent Kit v3.1. Libraries were sequenced on an Illumina NovaSeq S1 platform (paired-end reads) at the University of Queensland Sequencing Facility.

Human retinal microglia datasets were processed and analysed using the same pipeline described above for mouse microglia, including Cell Ranger for initial processing and Seurat for downstream analysis. To compare transcriptomic profiles between single-cell and bulk RNA-sequencing datasets, we performed correlation analyses. Bulk RNA-seq data for retinal microglia (36), reported as FPKM-normalized values, were averaged across 10 donors and scaled by a factor of 1×10⁶ to approximate raw count distributions. These values were then rounded and processed using the edgeR package to compute log2 counts-per-million (logCPM). For single-cell RNA-seq, data from the Retina Seurat object were used, with microglial clusters 3 and 5 (non-microglial cells) excluded from the analysis. Microglial cells were aggregated by summing gene-level raw counts to generate a pseudobulk profile, followed by logCPM transformation using edgeR. Pearson correlation was computed between the logCPM values of the bulk and single-cell pseudobulk datasets to assess overall concordance. The same method was applied to compare the scRNA-seq data from the scRNA human retinal microglia to brain microglia scRNA-seq data from Olah *et al*. (37).

### Statistical methods

For transcriptomic studies of mouse and human cells, sample sizes were determined based on preliminary data regarding minimum numbers of cells required to obtain a sufficient quantity of RNA for bulk RNA-seq analysis, and a minimum number of cells required for single cell RNA-seq analysis. Microglia density and morphological data were analysed for normality using a normality and log normality test (Shapiro-Wilk test or Kolmogorov-Smirnov test). Following this, data were analysed using either a one-way ANOVA (for parametric data) or a Kruskal Wallis test (for non-parametric data) using Graph Pad Prism 9.4.1.

## Supporting information

Supplemental data description

## Key resources table

| Reagent/resource | Source | Product identifier | Starting concentration | Working concentration/dilution |
| --- | --- | --- | --- | --- |
| **Antibodies** |  |  |  |  |
| Anti-IFIT1 antibody | Abcam | ab236256 | - | 1/400 |
| Anti-mouse Tmem119 | Synaptic Systems | 400 004 | - | 1/1000 |
| CCL4 polyclonal antibody | Invitrogen | PA5-114961 | 1 mg/mL | 1/400 |
| Goat anti-Guinea Pig IgG (H+L) Highly Cross-Adsorbed Secondary Antibody, Alexa Fluor 594 | Life Technologies | A11076 | 2 mg/mL | 1/400 |
| Goat anti-rabbit IgG (H+L) Alexa Fluor 594 | Life Technologies | A11012 | 2 mg/mL | 1/400 |
| Hoechst 33342 Solution (20 mM) | Life Technologies | 62249 | 20 mM | 1/2000 |
| Human Fc block | BD Biosciences | 564220 | 0.5 mg/mL | 2.5 µg per million cells |
| Mouse anti-Human CD45 antibody BV421 | BD Biosciences | 563879 | - | 2.5 µL per million cells |
| Mouse anti-mouse Cx3cr1-FITC | Biolegend | 149020 | 0.5 mg/mL | 1 µg per million cells |
| Purified rat anti-mouse CD16/CD32 (Fc Block) | BD Biosciences | 553142 | 0.5 mg/mL | 0.5 µg per million cells |
| Rat anti- mouse CD11b-APC-Cy7 | BD Biosciences | 557657 | 0.2 mg/mL | 0.25 µg per million cells |
| Rat anti-mouse CD11b APC-Cy7 | Invitrogen | A15390 | 0.1 mg/mL | 0.25 µg per million cells |
| Rat anti- mouse CD45-BUV395 | BD Biosciences | 564279 | 0.2 mg/mL | 0.5 µg per million cells |
| Rat anti-mouse CD45 BV421 | BD Biosciences | 563890 | 0.2 mg/mL | 0.25 µg per million cells |
| **Chemicals and media** |  |  |  |  |
| Bovine serum albumin | Sigma Aldrich | A7906-500G |  |  |
| Collagenase D | Sigma Aldrich | 11088866001 |  | 2.5 mg/mL |
| Cryostor CS10 | Merck | C2874 |  |  |
| D (+)-Glucose | Merck | 1.08337.0250 |  |  |
| DMEM/F12 | Gibco | 11320033 |  |  |
| DNase I | Sigma Aldrich | 10104159001 |  |  |
| EDTA | Merck | MER_1084540250-500G |  |  |
| Ethylene glycol | Sigma Aldrich | 324558-1L |  |  |
| HBSS (calcium, magnesium, no phenol red) | Life Technologies | 14025092 |  |  |
| HEPES | Merck | H4034-100G |  |  |
| Isopentane | Merck | 1.06056.1000 |  |  |
| NaH<sub>2</sub>PO<sub>4</sub> | Merck | 1.06346.0500 |  |  |
| Paraformaldehyde (PFA) | Sigma Aldrich | P6148-500G |  |  |
| Percoll | GE Healthcare | 17089102 |  |  |
| Polyvinylpyrrolidone | Sigma Aldrich | PVP40-100G |  |  |
| Propidium iodide | BD Biosciences | 556463 |  | 1 µg/mL |
| Sucrose | Sigma Aldrich | S0389-1KG |  |  |
| Tissue-Tek Optimal Cutting Medium | Sakura Finetak | 4583 |  |  |
| Triton x-100 | Sigma Aldrich | X100-100ML |  |  |
| **Consumables** |  |  |  |  |
| 70 µm nylon cell strainer | Falcon | 352350 |  |  |
| Coverslip No. 1.5 | ProSciTech | G425-2240 |  |  |
| Superfrost™ Plus Microscope Slides | Fisher Scientific | 12-550-15 |  |  |

## Acknowledgements

The authors thank the facilities, and the scientific and technical assistance of QUT Central Analytical Research Facility, QUT eResearch, QIMR Berghofer Animal Facility, QIMR Berghofer Flow Cytometry Facility, Monash Animal Research Platform, Monash Flow Core, Monash Micro Imaging, Monash Health Translation Precinct Medical Genomics Facility, Monash Bioinformatics Platform, University of Queensland Sequencing Facility. We acknowledge the assistance of the Queensland Eye Bank, the donor and their family, for the generous donation of retinal tissue used in this study.

## Authors’ contributions

Conceptualisation: FE, SJD; Methodology: FE, PW, DGH, HQ, EES, PGM, SJD; Investigation: FE, PW, DGH, EES, SJD; Resources: SJD; Writing – original draft: FE, PW, SJD; Writing – review & editing: FE, PW, DGH, HQ, EES, PGM, SJD; Supervision: SJD, DGH; Funding acquisition: SJD, DGH.

## Funding

This study was supported by an Australian Research Council Discovery Early Career Research Award (DE180101075) to SJD, and a QUT inter-program collaborative grant to SJD and DGH.

## Notes

### Competing Interest Statement

The authors have declared no competing interest.

