## Supplemental data description for "Regional and sub-regional microglial heterogeneity in the steady-state mouse brain and retina"

**Supplementary Data**

**Figure S1. Gating strategy for isolation of microglia by FACS for bulk RNA sequencing.** Density dot plots display sorting regions for each tissue (cortex, hippocampus, olfactory bulb, cerebellum, and retina). From left to right: Gating of single cells; exclusion of dead cells by propidium iodide staining; gating of intact cells based on side scatter (SSC) and forward scatter (FSC) properties; gating microglia based on their $CD11b^{+}$ $CD45^{intermediate}$ expression.

**Figure S2. Bulk RNA-seq analysis of the expression of microglia homeostatic genes and myeloid genes in different regions of the healthy adult C57Bl/6J mouse CNS. (a)** Violin plots displaying gene expression fold change ($Log_2$ FC) of homeostatic markers from microglia isolated from different regions of CNS normalized to cortical microglia (dotted line). **(b)** Violin plots representing the $Log_2$ FC gene expression profiles of myeloid genes expressed by microglia regional populations. *Itgam* is the gene for CD11b; *AIF-1* gene product is Iba1. Each data point represents one biological replicate, which consists of pooled microglia sorted from n=5 mice. Total of n=4 biological replicates per CNS region.

**Figure S3. Gating strategy for isolation of microglia by FACS for single cell RNA sequencing.** Density dot plots show the gating strategy applied to isolate microglia from the mouse cortex, hippocampus, cerebellum, olfactory bulb, and retina. From left to right: Intact cells populations were refined to remove debris (P1). Dead cells were identified and excluded from the analysis using propidium iodide (PI) staining (P2). For each tissue, doublets were excluded by FSC-A versus FSC-H gating (P3). Microglia were sorted based on CD45+ CD11b+ gating (P4), and were verified to express Cx3Cr1 (final panel).

**Figure S4. Expression of microglia signature genes (*Tmem119*, *P2ry12*, *Fcrls*, and *Slc2a5*).** tSNE plots of microglia from each CNS region coloured for expression of *Tmem119*, *P2ry12*, *Fcrls* and *Slc2a5*. Grey: low expression; red: high expression.

**Figure S5. Expression of myeloid markers in microglia (*Cx3cr1*, *Ptprc*, *Itgam*, and *Aif1*).** tSNE plots of microglia from each CNS region coloured for expression of *Cx3cr1*, *Ptprc*, *Itgam*, *and Aif1*. Grey: low expression; red: high expression.

**Figure S6. Expression of *ex vivo* activation markers in microglia.** Dot plots showing the expression of microglia activation genes including *Fos*, *Jun*, *Hspala*, *Dusp1, Ccl3 and Ccl4*, with the dot size representing the percentage of cells expressing the gene and the colour representing its average expression within a cluster.

**Figure S7. Gene ontology network of top 100 GO terms based on genes that are differentially expressed between the CC cluster and other microglia clusters in each CNS region.** Each node represents a gene ontology and the main themes within the data group in clusters.

**Figure S8. Gene ontology network of top 100 GO terms based on genes that are differentially expressed between the IFN cluster and other microglia clusters in each CNS region.** Each node represents a gene ontology and the main themes within the data group in clusters.
