## Supplementary figures and images for "Regional and sub-regional microglial heterogeneity in the steady-state mouse brain and retina"

### Supplemental figure 1

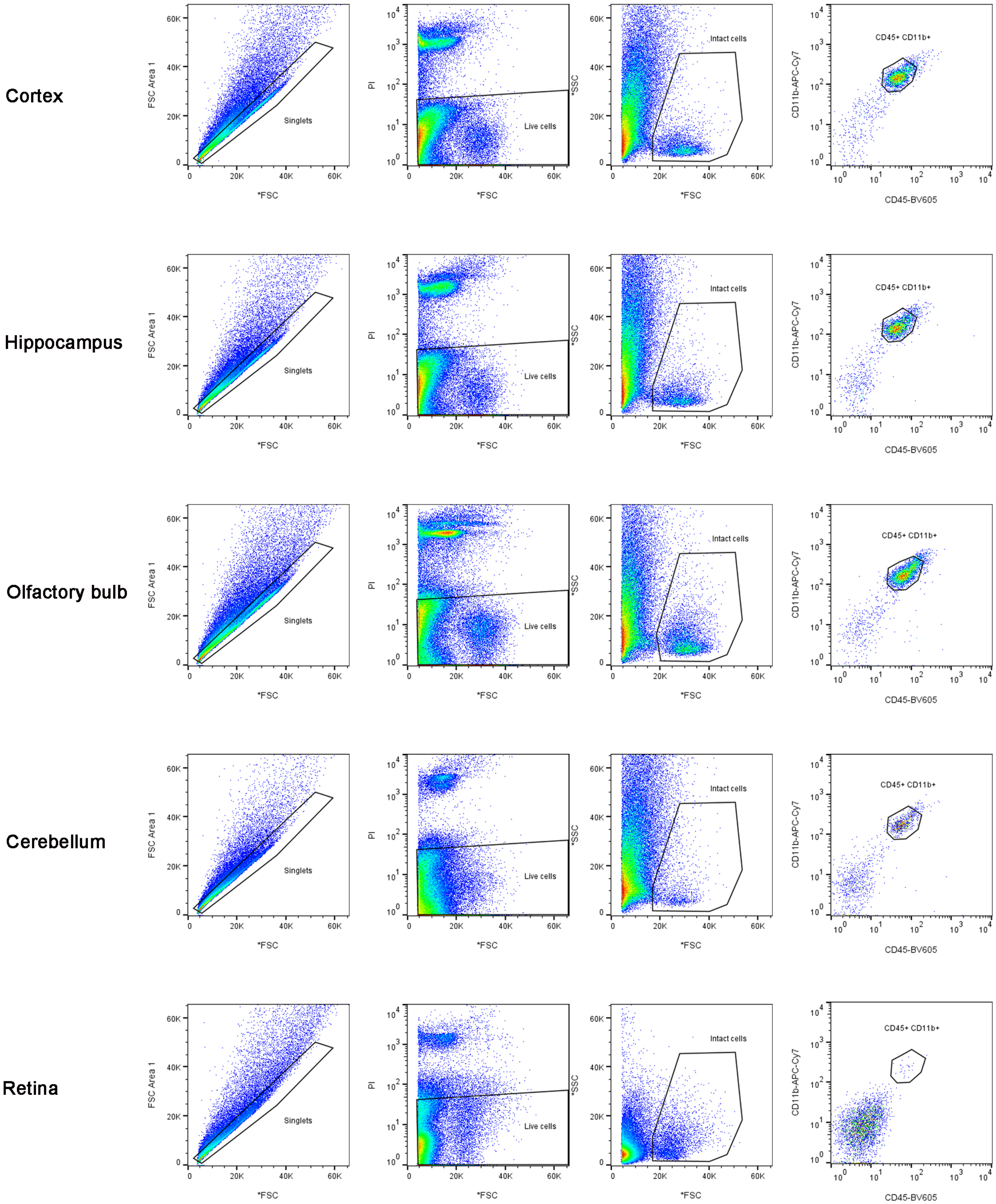

### Supplemental figure 2

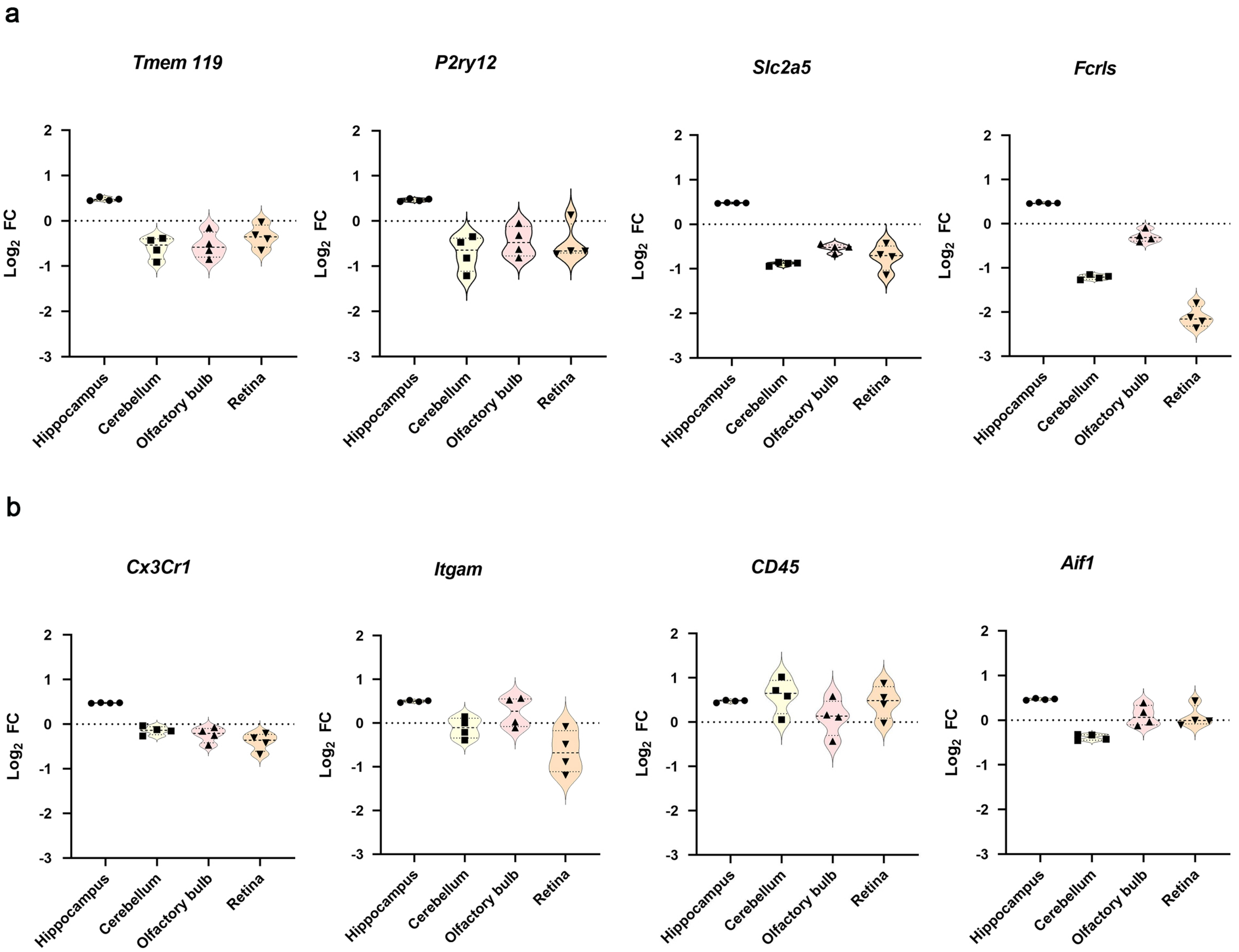

### Supplemental figure 3

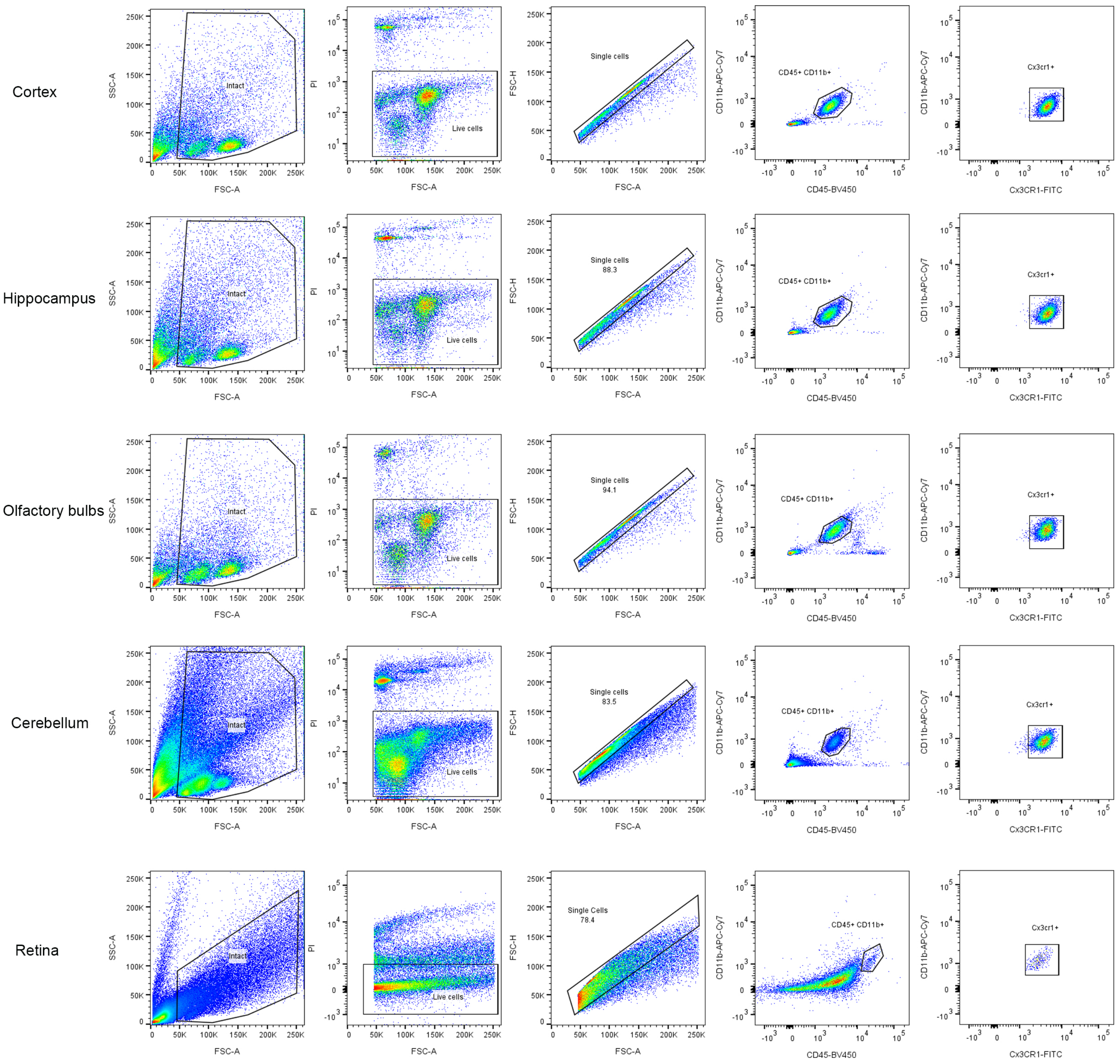

### Supplemental figure 4

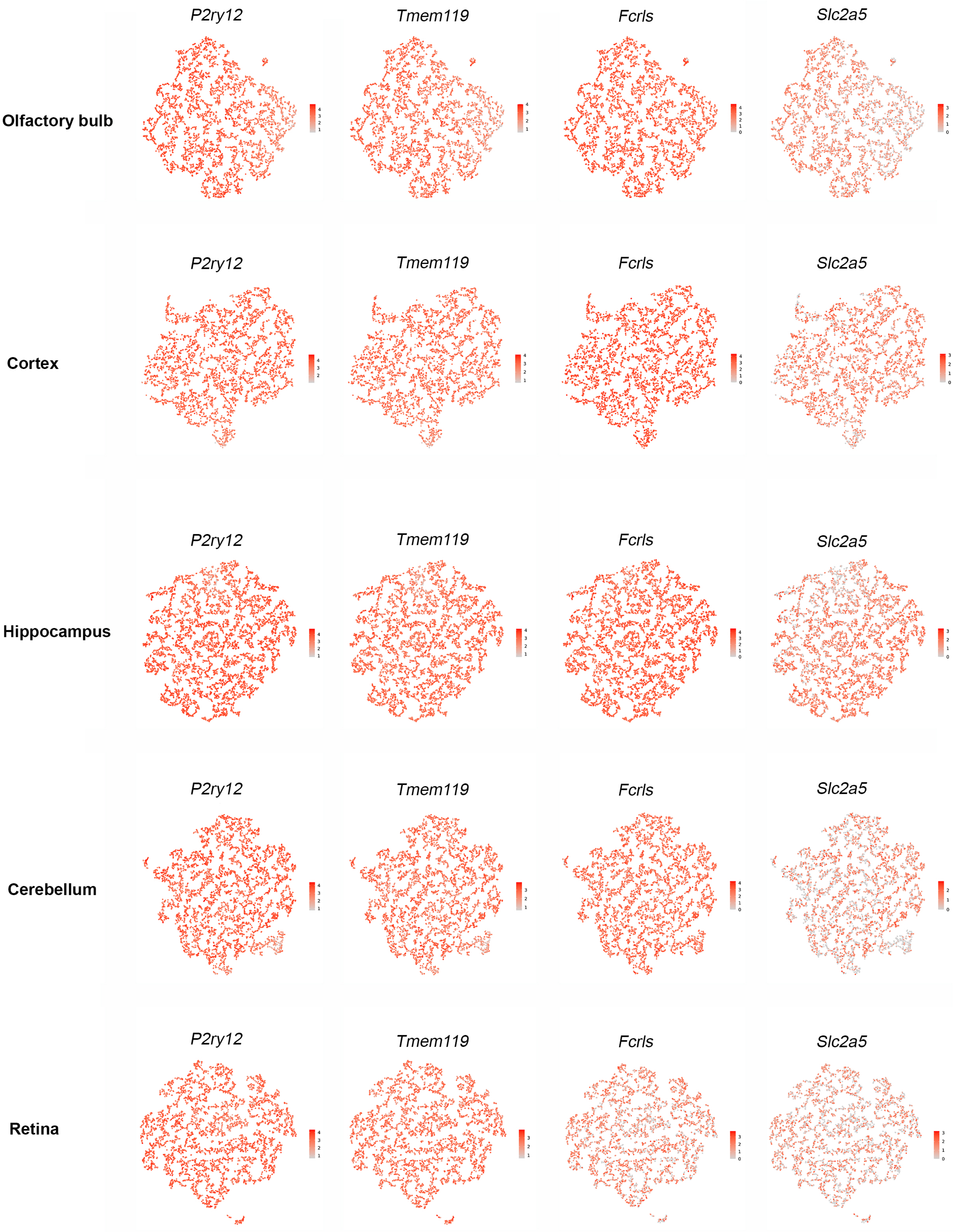

### Supplemental figure 5

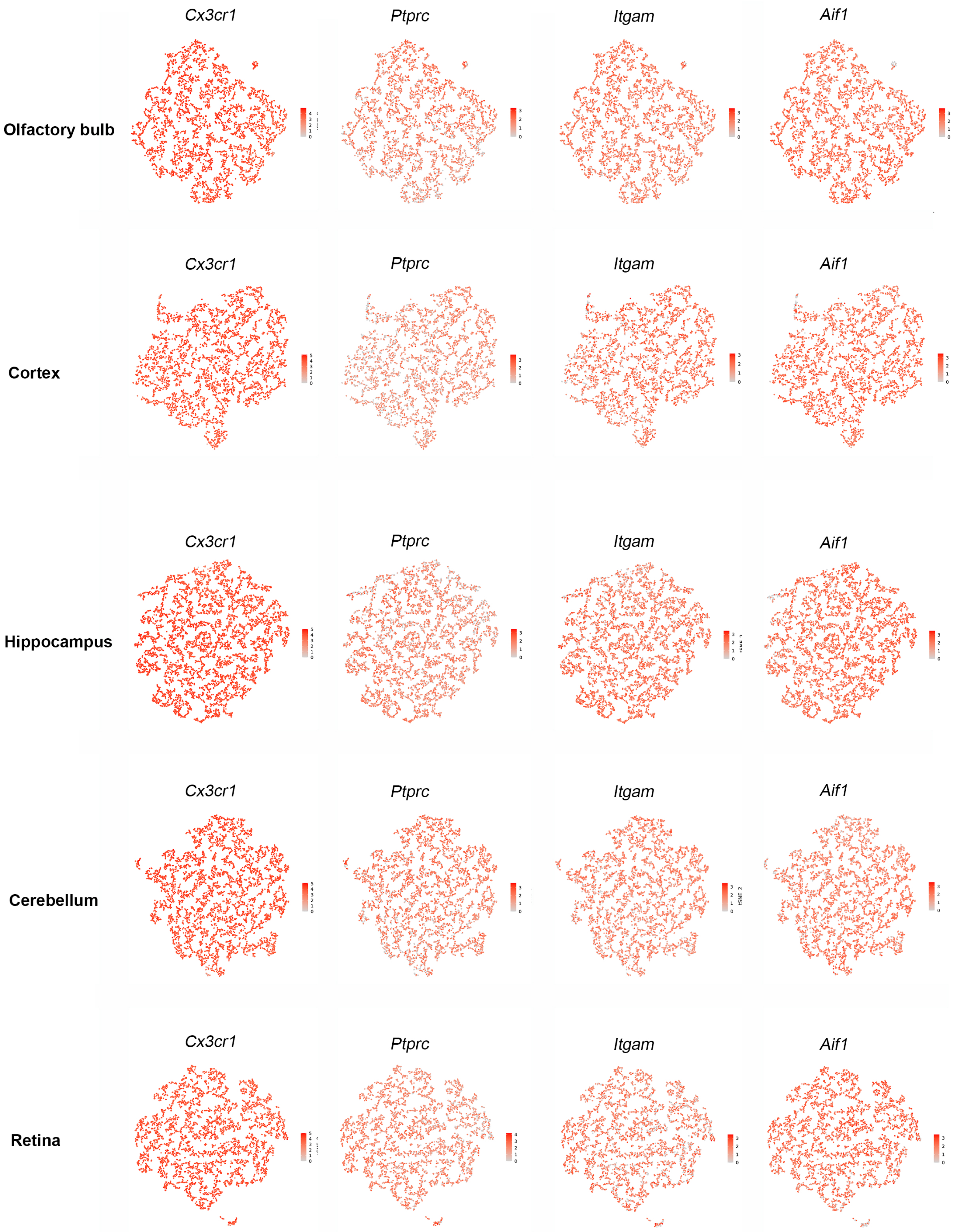

### Supplemental figure 6

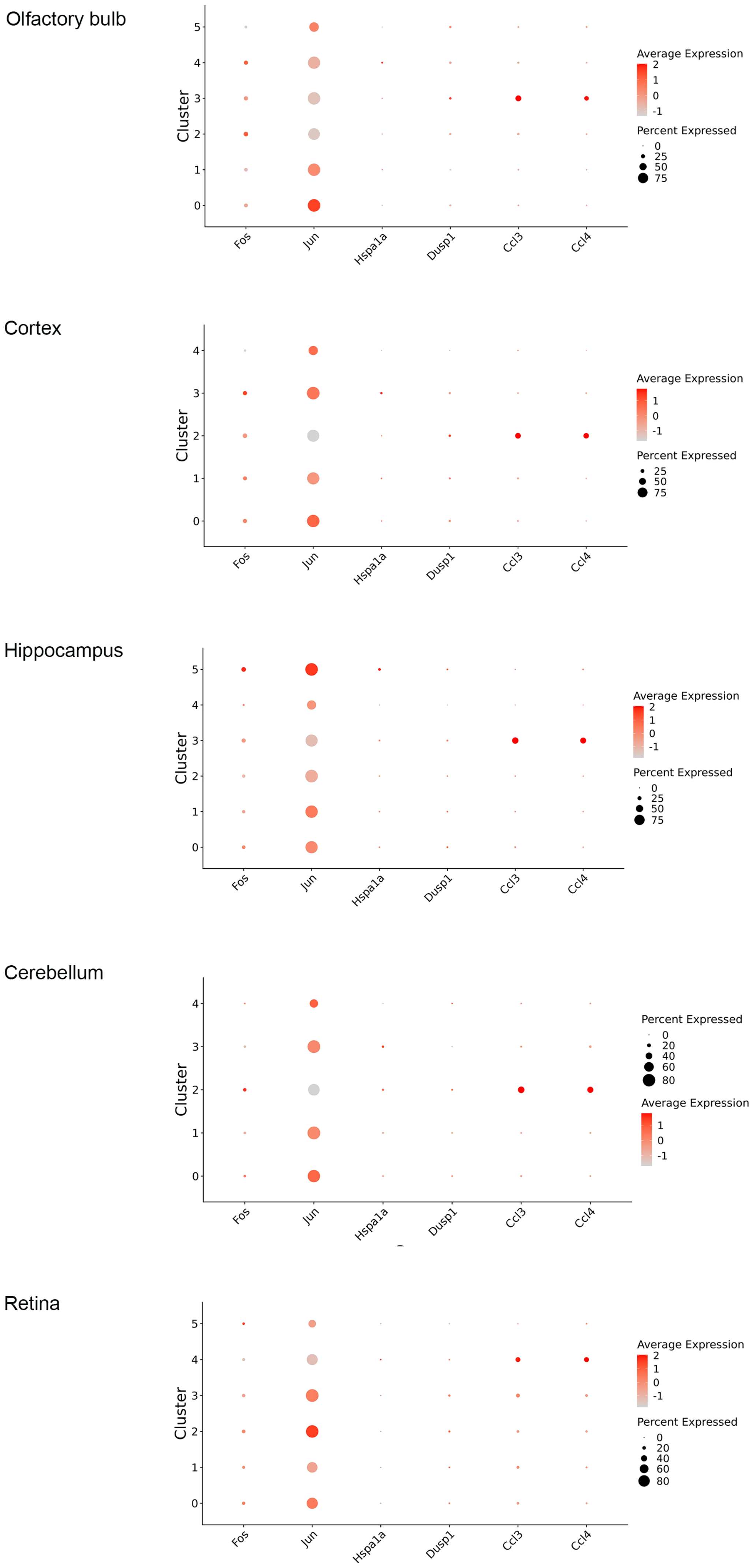

### Supplemental figure 7

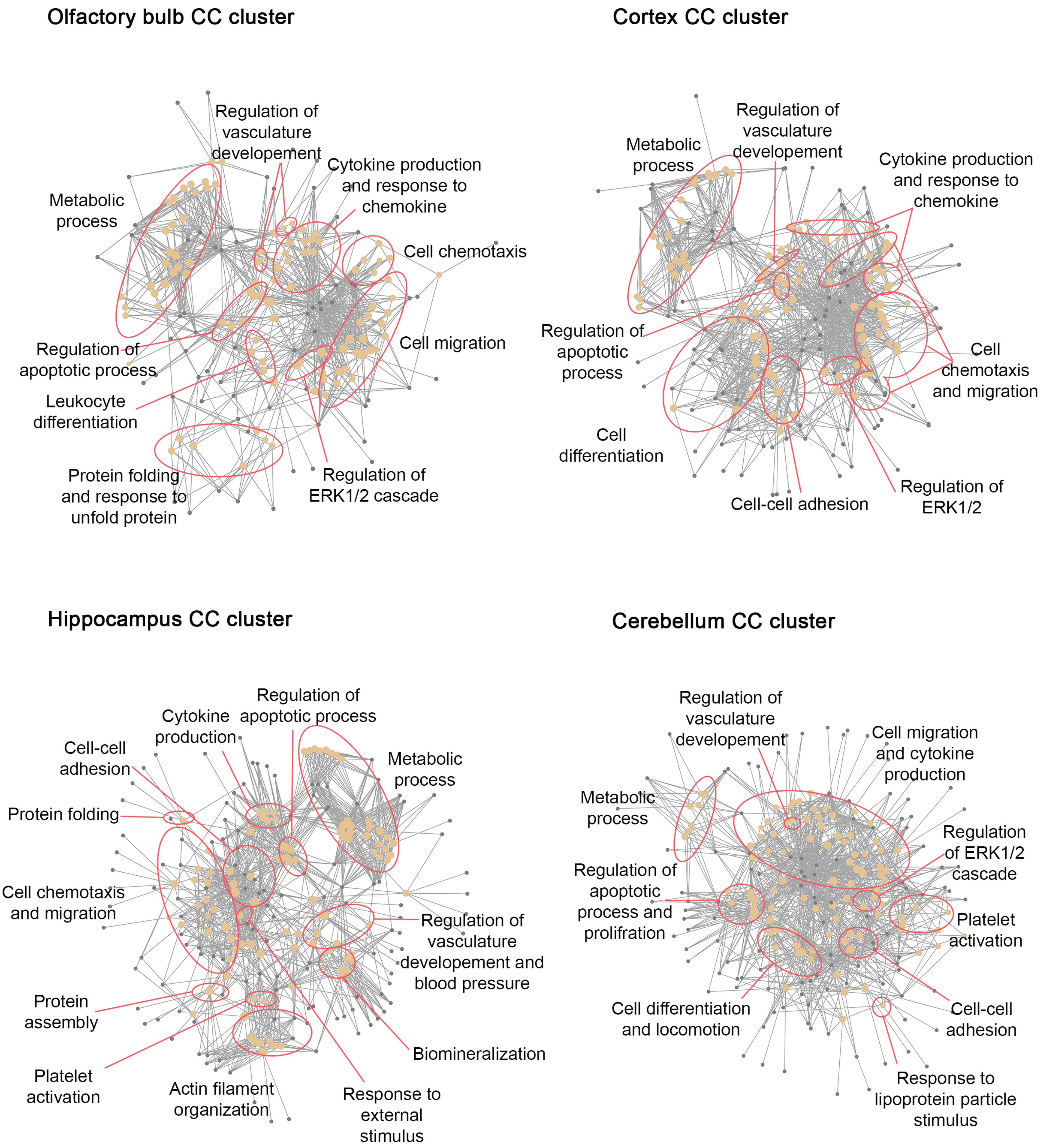

### Supplemental figure 8

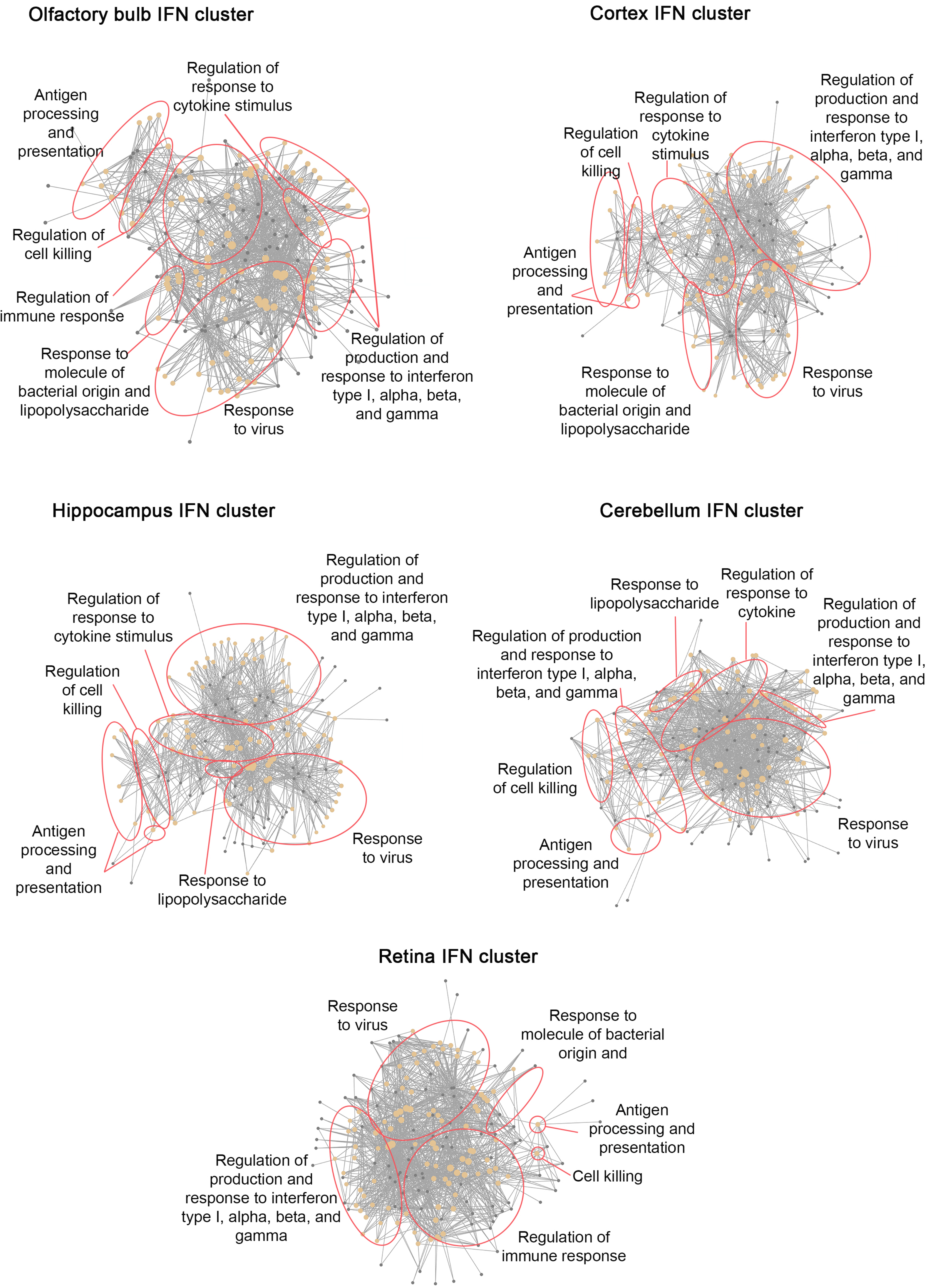
