## Supplemental file 1 for "Regional and sub-regional microglial heterogeneity in the steady-state mouse brain and retina"

Enriched GO terms for biological process in DEGs using PANTHER

| Biological process (Gene ontology accession) | Number of DEGs | Percentage of total DEGs |
|---|---|---|
| Cellular process (GO:0009987) | 261 | 22.30% |
| Biological regulation (GO:0065007) | 175 | 14.90% |
| Metabolic process (GO:0008152) | 148 | 12.60% |
| Response to stimulus (GO:0050896) | 127 | 10.80% |
| Localization (GO:0051179) | 94 | 8.00% |
| Signalling (GO:0023052) | 92 | 7.80% |
| Cellular component organization or biogenesis (GO:0071840) | 73 | 6.20% |
| Developmental process (GO:0032502) | 44 | 3.80% |
| Multicellular organismal process (GO:0032501) | 41 | 3.50% |
| Immune system process (GO:0002376) | 39 | 3.30% |
| Multi-organism process (GO:0051704) | 29 | 2.50% |
| Locomotion (GO:0040011) | 20 | 1.70% |
| Cell population proliferation (GO:0008283) | 6 | 0.50% |
| Growth (GO:0040007) | 6 | 0.50% |
| Biological adhesion (GO:0022610) | 5 | 0.40% |
| Reproduction (GO:0000003) | 4 | 0.30% |
| Reproductive process (GO:0022414) | 4 | 0.30% |
| Biomineralization (GO:0110148) | 2 | 0.20% |
| Biological phase (GO:0044848) | 2 | 0.20% |
| Behaviour (GO:0007610) | 1 | 0.10% |
| Total* | 1137 | 100% |

Note: the number of DEGs identified in the bulk RNA-seq dataset is 747. PANTHER analysis records the total number of DEGs as 1137, as some genes are involved in multiple biological processes and GO terms.
