## Supplemental file 2 for "Regional and sub-regional microglial heterogeneity in the steady-state mouse brain and retina"

DEGs (747 genes).

| Gene.ID | Chrom | Gene.Name | Biotype | cerebellum | cortex | hippocampus | ob | retina | FDR | P value |
|---|---|---|---|---|---|---|---|---|---|---|
| ENSMUSG00000029304 | 5 | Spp1 | protein_coding | 6.130871893 | 0 | 2.110631952 | 5.894457 | 8.079708 | 8.23E-08 | 1.33E-10 |
| ENSMUSG00000055148 | 8 | Klf2 | protein_coding | 5.04871599 | 0 | 1.44755254 | 2.851827 | 5.431431 | 3.76E-08 | 4.87E-11 |
| ENSMUSG00000030657 | 7 | Xylt1 | protein_coding | 4.686812561 | 0 | 2.80514007 | 2.310783 | 5.674133 | 2.16546E-05 | 2.24E-07 |
| ENSMUSG00000041261 | 4 | Car8 | protein_coding | 4.276518124 | 0 | 1.652721012 | -0.035895 | 1.959692 | 0.006114158 | 0.000236169 |
| ENSMUSG00000026822 | 2 | Lcn2 | protein_coding | 4.218347094 | 0 | 2.463422271 | 5.07716 | -0.792525 | 0.000220894 | 4.05728E-06 |
| ENSMUSG00000039313 | 9 | AF529169 | protein_coding | 4.143877333 | 0 | 1.480160431 | 4.158636 | 3.090858 | 0.014659706 | 0.000766692 |
| ENSMUSG00000050335 | 14 | Lgals3 | protein_coding | 4.128110423 | 0 | 1.28040307 | 3.426976 | 5.630613 | 3.60653E-06 | 1.87E-08 |
| ENSMUSG00000031722 | 8 | Hp | protein_coding | 3.951189841 | 0 | 3.044361538 | 5.169951 | 1.111205 | 0.026305779 | 0.001638626 |
| ENSMUSG00000044734 | 13 | Serpinb1a | protein_coding | 3.933228262 | 0 | 1.025648633 | 4.345552 | 0.972788 | 0.03112607 | 0.002076855 |
| ENSMUSG00000026728 | 2 | Vim | protein_coding | 3.874414689 | 0 | 2.13705168 | 2.516978 | 4.286879 | 0.010065091 | 0.00045317 |
| ENSMUSG00000028028 | 3 | Alpk1 | protein_coding | 3.778678259 | 0 | 3.257317856 | 3.34867 | 3.47122 | 0.007612838 | 0.000310146 |
| ENSMUSG00000059479 | 7 | B3gnt8 | protein_coding | 3.75816758 | 0 | 3.153035474 | 2.698413 | 4.688742 | 0.017035025 | 0.000932519 |
| ENSMUSG00000037849 | 1 | Ifi206 | protein_coding | 3.729584851 | 0 | 0.490824938 | 4.183982 | 4.260252 | 0.000102738 | 1.55452E-06 |
| ENSMUSG00000003032 | 4 | Klf4 | protein_coding | 3.657105665 | 0 | 2.820081303 | 3.022605 | 2.914722 | 0.012745836 | 0.000629602 |
| ENSMUSG00000028362 | 4 | Tnfsf8 | protein_coding | 3.649454062 | 0 | 0.981869116 | -0.742675 | 5.328515 | 1.06196E-06 | 3.35E-09 |
| ENSMUSG00000026835 | 2 | Fcnb | protein_coding | 3.590855302 | 0 | 2.568371157 | 4.999078 | 0.769484 | 0.011464118 | 0.000539368 |
| ENSMUSG00000033740 | 1 | St18 | protein_coding | 3.531775743 | 0 | 1.312896185 | -0.035895 | 3.07373 | 0.017477185 | 0.000960969 |
| ENSMUSG00000037995 | 1 | Igsf9 | protein_coding | 3.47499305 | 0 | 1.884161159 | 2.552665 | 4.378664 | 0.00300018 | 9.5722E-05 |
| ENSMUSG00000022584 | 15 | Ly6c2 | protein_coding | 3.457367713 | 0 | 1.755172302 | 5.072746 | 0.60716 | 0.002511287 | 7.58531E-05 |
| ENSMUSG00000020641 | 12 | Rsad2 | protein_coding | 3.434427312 | 0 | 0.959881235 | 3.13731 | 4.759717 | 4.46031E-05 | 5.76E-07 |
| ENSMUSG00000000673 | 17 | Haao | protein_coding | 3.411179597 | 0 | 1.205756557 | 0.785688 | 5.425677 | 1.99956E-06 | 7.77E-09 |
| ENSMUSG00000079445 | 1 | B3gnt7 | protein_coding | 3.386073223 | 0 | 0.819099989 | 2.51634 | 4.71227 | 0.005607648 | 0.000210701 |
| ENSMUSG00000022893 | 16 | Adamts1 | protein_coding | 3.344208329 | 0 | -0.547392689 | -1.979088 | 1.923514 | 0.002082491 | 5.98659E-05 |
| ENSMUSG00000003484 | 8 | Cyp4f18 | protein_coding | 3.256435633 | 0 | 0.354558377 | 1.021806 | 6.332474 | 2.14258E-06 | 9.02E-09 |
| ENSMUSG00000090877 | 17 | Hspa1b | protein_coding | 3.225907883 | 0 | 0.802603996 | -0.395809 | -0.405287 | 0.001293144 | 3.44517E-05 |
| ENSMUSG00000053279 | 19 | Aldh1a1 | protein_coding | 3.216619629 | 0 | 1.73323415 | -0.814396 | 1.114396 | 0.046460874 | 0.003698359 |
| ENSMUSG00000091971 | 17 | Hspa1a | protein_coding | 3.205567175 | 0 | -0.01094095 | -0.214826 | 1.008945 | 0.010304692 | 0.000467295 |
| ENSMUSG00000032496 | 9 | Ltf | protein_coding | 3.196058839 | 0 | 1.839351386 | 4.802103 | -1.74639 | 0.006169409 | 0.000239303 |
| ENSMUSG00000032690 | 5 | Oas2 | protein_coding | 3.188620211 | 0 | 1.555165453 | 4.717833 | 3.852152 | 0.001851768 | 5.24835E-05 |
| ENSMUSG00000040883 | 9 | Tmem205 | protein_coding | 3.08913085 | 0 | 1.680620631 | 2.107729 | 4.017104 | 0.000334876 | 6.7523E-06 |
| ENSMUSG00000004630 | 8 | Pcp2 | protein_coding | 3.05961991 | 0 | -0.336623544 | 0.342106 | 4.371969 | 0.002217581 | 6.44677E-05 |
| ENSMUSG00000027230 | 2 | Creb3l1 | protein_coding | 3.049834922 | 0 | 1.284879768 | 0.260387 | 3.859891 | 0.041273613 | 0.00314507 |
| ENSMUSG00000026068 | 1 | Il18rap | protein_coding | 3.032073139 | 0 | 0.171030813 | 0.676228 | 3.637547 | 0.00604449 | 0.000232989 |
| ENSMUSG00000056313 | 8 | 1810011O10Rik | protein_coding | 3.013310432 | 0 | 0.965462159 | -2.241433 | 4.188551 | 2.30E-08 | 2.42E-11 |
| ENSMUSG00000036875 | 10 | Dna2 | protein_coding | 2.977365819 | 0 | 1.767047686 | 0.592953 | 3.918154 | 0.000820599 | 1.98688E-05 |
| ENSMUSG00000030317 | 6 | Timp4 | protein_coding | 2.975342769 | 0 | -0.728289035 | 0.917303 | -1.473929 | 0.022519338 | 0.001333532 |
| ENSMUSG00000037580 | 14 | Gch1 | protein_coding | 2.955644753 | 0 | 1.399473051 | 3.112621 | 2.358734 | 0.008946785 | 0.000388329 |
| ENSMUSG00000021458 | 13 | 2010111I01Rik | protein_coding | 2.950252099 | 0 | 1.871331569 | 0.867312 | 3.28167 | 0.005131164 | 0.000191551 |
| ENSMUSG00000036067 | 2 | Slc2a6 | protein_coding | 2.923032705 | 0 | 1.722958662 | 2.79965 | 5.237169 | 0.000813811 | 1.96385E-05 |
| ENSMUSG00000010066 | 9 | Cacna2d2 | protein_coding | 2.885060464 | 0 | -1.05680879 | -0.75608 | 2.020937 | 0.006115128 | 0.000236702 |
| ENSMUSG00000015568 | 8 | Lpl | protein_coding | 2.878604358 | 0 | 1.00115449 | 2.521777 | 2.413033 | 3.00102E-06 | 1.43E-08 |
| ENSMUSG00000041754 | 17 | Trem3 | protein_coding | 2.877504377 | 0 | 1.362237272 | 5.050046 | 1.071254 | 0.012787272 | 0.000634756 |
| ENSMUSG00000020914 | 11 | Top2a | protein_coding | 2.833182506 | 0 | 3.040154165 | 4.698335 | 4.723934 | 0.000404733 | 8.55415E-06 |
| ENSMUSG00000036594 | 17 | H2-Aa | protein_coding | 2.763839343 | 0 | 0.767134826 | 2.745465 | 0.861272 | 0.026825733 | 0.001685064 |
| ENSMUSG00000021886 | 12 | Gpr65 | protein_coding | 2.754745397 | 0 | 1.004446867 | 2.044541 | 3.713108 | 1.75983E-06 | 6.41E-09 |
| ENSMUSG00000057191 | 9 | AB124611 | protein_coding | 2.7305726 | 0 | 1.730521329 | 3.492226 | 4.490252 | 0.003417808 | 0.000111537 |
| ENSMUSG00000048022 | 6 | Tmem229a | protein_coding | 2.721476297 | 0 | 1.023260157 | 2.517326 | 3.511284 | 0.036714809 | 0.002625247 |
| ENSMUSG00000039081 | 14 | Zfp503 | protein_coding | 2.68269915 | 0 | 1.565036678 | -0.83467 | 3.502929 | 0.005723298 | 0.000216437 |
| ENSMUSG00000074151 | 8 | Nlrc5 | protein_coding | 2.677741073 | 0 | -2.007085048 | 1.880921 | 3.386167 | 0.004918854 | 0.000178049 |
| ENSMUSG00000079017 | 12 | Ifi27l2a | protein_coding | 2.677370498 | 0 | 0.111119371 | 2.748787 | 3.705799 | 9.21E-07 | 2.83E-09 |
| ENSMUSG00000026832 | 2 | Cytip | protein_coding | 2.66527032 | 0 | 3.076647897 | 3.156113 | 4.888219 | 0.023174886 | 0.001388729 |
| ENSMUSG00000033208 | 10 | S100b | protein_coding | 2.656448216 | 0 | 1.292527193 | 1.994349 | -2.405363 | 0.026903732 | 0.001692785 |
| ENSMUSG00000090942 | 18 | F830016B08Rik | protein_coding | 2.651569971 | 0 | 0.138740687 | 1.673826 | -1.45007 | 0.007043891 | 0.000280067 |
| ENSMUSG00000037725 | 8 | Ckap2 | protein_coding | 2.647801053 | 0 | 0.49882726 | 3.558152 | 3.486743 | 0.010314277 | 0.000468565 |
| ENSMUSG00000040189 | 7 | Ccdc114 | protein_coding | 2.619610807 | 0 | -0.731544602 | 2.764558 | 0.939812 | 0.034958977 | 0.00244308 |
| ENSMUSG00000004709 | 1 | Cd244 | protein_coding | 2.589690726 | 0 | 0.383560156 | 1.041788 | 4.498612 | 0.000716403 | 1.70558E-05 |
| ENSMUSG00000024675 | 19 | Ms4a4c | protein_coding | 2.575877339 | 0 | 0.954983241 | 2.742857 | 3.997618 | 0.008926128 | 0.000384541 |
| ENSMUSG00000037474 | 1 | Dtl | protein_coding | 2.564443838 | 0 | 1.664185608 | 4.701923 | 5.128948 | 3.87527E-05 | 4.86E-07 |
| ENSMUSG00000073490 | 1 | Ifi207 | protein_coding | 2.561614701 | 0 | 0.839202023 | 2.890929 | 2.862795 | 0.019328055 | 0.001089345 |
| ENSMUSG00000060586 | 17 | H2-Eb1 | protein_coding | 2.549252212 | 0 | 0.554757167 | 3.064072 | 1.123164 | 0.008842676 | 0.000377366 |
| ENSMUSG00000041774 | 16 | Ydjc | protein_coding | 2.533414305 | 0 | 1.524570292 | 1.133859 | 3.867703 | 0.041279158 | 0.003148835 |
| ENSMUSG00000034701 | 2 | Neurod1 | protein_coding | 2.521702077 | 0 | 0.31714235 | -1.290041 | 5.803472 | 1.08968E-05 | 8.59E-08 |
| ENSMUSG00000020330 | 11 | Hmmr | protein_coding | 2.493540349 | 0 | 1.309889056 | 4.73921 | 2.671565 | 0.044566264 | 0.003482585 |
| ENSMUSG00000028518 | 4 | Prkaa2 | protein_coding | 2.485929412 | 0 | 0.557047195 | 0.578671 | 3.957235 | 0.032355799 | 0.002195656 |
| ENSMUSG00000031497 | 8 | Tnfsf13b | protein_coding | 2.485867573 | 0 | 0.736677785 | 0.631162 | 4.003227 | 1.91E-08 | 1.85E-11 |
| ENSMUSG00000027435 | 2 | Cd93 | protein_coding | 2.48559373 | 0 | 0.754709423 | 2.797001 | -0.867174 | 0.014627513 | 0.000761653 |
| ENSMUSG00000037217 | X | Syn1 | protein_coding | 2.471212698 | 0 | 0.650909273 | 3.785752 | 1.476698 | 0.041965017 | 0.003224941 |
| ENSMUSG00000006360 | 12 | Crip1 | protein_coding | 2.455001789 | 0 | -0.357025897 | 1.515539 | 1.743148 | 0.020035526 | 0.001142199 |
| ENSMUSG00000035458 | 7 | Tnni3 | protein_coding | 2.451246663 | 0 | 1.451282617 | 3.467853 | 2.823643 | 0.042964872 | 0.003329612 |
| ENSMUSG00000054206 | 10 | Gzmm | protein_coding | 2.427753326 | 0 | 0.589743301 | 0.18375 | 3.68567 | 0.049999429 | 0.004093402 |
| ENSMUSG00000040564 | 7 | Apoc1 | protein_coding | 2.424415568 | 0 | 2.012511782 | 4.565308 | 5.558918 | 0.000752472 | 1.80974E-05 |
| ENSMUSG00000041552 | X | Ptchd1 | protein_coding | 2.415953711 | 0 | -1.404035306 | 0.04136 | 3.199625 | 2.86603E-05 | 3.27E-07 |

| | | | | | | | | | | |
|---|---|---|---|---|---|---|---|---|---|---|
| ENSMUSG00000026285 | 1 | Pdcd1 | protein_coding | 2.380386121 | 0 | 0.217601256 | -0.914094 | 4.894188 | 2.67315E-05 | 3.03E-07 |
| ENSMUSG00000003585 | 11 | Sec14l2 | protein_coding | 2.368516963 | 0 | 2.536049643 | 2.431793 | 3.682244 | 0.012477565 | 0.000610288 |
| ENSMUSG00000102418 | 1 | Sh2d1b1 | protein_coding | 2.353961499 | 0 | 2.177736177 | 2.135681 | 4.311566 | 0.035778373 | 0.002519399 |
| ENSMUSG00000028874 | 4 | Fgr | protein_coding | 2.29192519 | 0 | 1.365197863 | 1.578893 | 3.242645 | 0.020077025 | 0.00114619 |
| ENSMUSG00000040212 | 7 | Emp3 | protein_coding | 2.286525349 | 0 | 0.448071301 | 1.048054 | 3.137795 | 1.29E-09 | 8.33E-13 |
| ENSMUSG00000009687 | 7 | Fxyd5 | protein_coding | 2.282071423 | 0 | -0.655950054 | 1.553429 | 4.6939 | 1.14E-09 | 6.47E-13 |
| ENSMUSG00000073489 | 1 | Ifi204 | protein_coding | 2.253637942 | 0 | -0.604461152 | 1.840358 | 3.289616 | 1.54E-07 | 2.99E-10 |
| ENSMUSG00000034593 | 9 | Myo5a | protein_coding | 2.220943784 | 0 | 0.985047033 | 1.767319 | 1.916671 | 8.87619E-05 | 1.30818E-06 |
| ENSMUSG00000002602 | 7 | Axl | protein_coding | 2.220411384 | 0 | -0.791637238 | 2.306578 | 4.338706 | 8.70E-10 | 2.94E-13 |
| ENSMUSG00000059108 | 7 | Ifitm6 | protein_coding | 2.203127999 | 0 | 1.646961178 | 3.573787 | -1.198609 | 0.049821775 | 0.004068914 |
| ENSMUSG00000026536 | 1 | Ifi211 | protein_coding | 2.20227544 | 0 | 0.020324276 | 1.353624 | 3.517917 | 0.016642727 | 0.000903396 |
| ENSMUSG00000040809 | 3 | Chil3 | protein_coding | 2.183138169 | 0 | 1.13645529 | 3.8735 | -2.456845 | 0.009154324 | 0.00040287 |
| ENSMUSG00000022033 | 14 | Pbk | protein_coding | 2.176702503 | 0 | 3.730357443 | 4.213626 | 4.36119 | 0.010674836 | 0.000486685 |
| ENSMUSG00000049744 | 2 | Arhgap15 | protein_coding | 2.157293628 | 0 | 1.041045356 | 2.258004 | 4.567842 | 7.41E-07 | 2.22E-09 |
| ENSMUSG00000073421 | 17 | H2-Ab1 | protein_coding | 2.146275182 | 0 | 0.613631593 | 2.015808 | 1.919613 | 0.001175172 | 3.08329E-05 |
| ENSMUSG00000040963 | 11 | Asgr2 | protein_coding | 2.138550345 | 0 | 0.517507738 | 1.934638 | -1.860172 | 0.034258474 | 0.002380255 |
| ENSMUSG00000024610 | 18 | Cd74 | protein_coding | 2.125364882 | 0 | 0.55932368 | 2.390668 | 1.489495 | 0.000529195 | 1.17418E-05 |
| ENSMUSG00000064065 | 10 | Ipcef1 | protein_coding | 2.113628919 | 0 | -0.041932909 | 0.431344 | 1.928196 | 1.04152E-05 | 8.10E-08 |
| ENSMUSG00000022246 | 15 | Rai14 | protein_coding | 2.078461036 | 0 | -0.365353787 | 2.52696 | 3.515773 | 0.018105236 | 0.001005765 |
| ENSMUSG00000070868 | 4 | Skint3 | protein_coding | 2.060287582 | 0 | -1.747842511 | -1.843973 | 0.717866 | 0.010185975 | 0.000460262 |
| ENSMUSG00000015340 | X | Cybb | protein_coding | 2.052838911 | 0 | 0.797768693 | 2.290361 | 3.073937 | 3.7748E-06 | 2.02E-08 |
| ENSMUSG00000034656 | 8 | Cacna1a | protein_coding | 2.047615995 | 0 | 0.839437896 | 0.656752 | -1.362494 | 0.001640958 | 4.57114E-05 |
| ENSMUSG00000056071 | 3 | S100a9 | protein_coding | 2.03991683 | 0 | 0.655996476 | 3.489083 | -3.671769 | 0.005116918 | 0.0001905 |
| ENSMUSG00000038026 | 1 | Kcnj9 | protein_coding | 2.012423438 | 0 | -0.470146445 | 0.831227 | 0.333769 | 0.03112607 | 0.00207692 |
| ENSMUSG00000069792 | 11 | Wfdc17 | protein_coding | 2.009809349 | 0 | 0.837093707 | 1.988155 | 2.953935 | 0.030639592 | 0.002020656 |
| ENSMUSG00000022864 | 16 | D16Ertd472e | protein_coding | 2.000382645 | 0 | 0.978377682 | 1.437919 | 3.551183 | 0.008867414 | 0.000380576 |
| ENSMUSG00000025854 | 5 | Fam20c | protein_coding | 1.998387165 | 0 | 0.72495249 | 1.993991 | 3.121473 | 0.004343105 | 0.000154043 |
| ENSMUSG00000057135 | 11 | Scimp | protein_coding | 1.989417757 | 0 | 0.184618755 | 1.347371 | 3.633955 | 0.019498696 | 0.001100542 |
| ENSMUSG00000040711 | 11 | Sh3pxd2b | protein_coding | 1.976816844 | 0 | -2.60390799 | 1.556016 | 2.185418 | 0.000597757 | 1.3754E-05 |
| ENSMUSG00000020085 | 10 | Aifm2 | protein_coding | 1.953665634 | 0 | 1.733643523 | 3.188737 | 4.616147 | 0.015406118 | 0.000817152 |
| ENSMUSG00000041840 | 18 | Haus1 | protein_coding | 1.94613444 | 0 | 0.295347436 | 2.635558 | 2.595181 | 0.024812761 | 0.001502951 |
| ENSMUSG00000096370 | 19 | Gm21992 | protein_coding | 1.944471658 | 0 | 0.629661524 | 3.978739 | 1.832562 | 0.04622608 | 0.003662199 |
| ENSMUSG00000028773 | 4 | Fabp3 | protein_coding | 1.937400149 | 0 | -1.415343371 | 0.697503 | 2.628317 | 0.013517343 | 0.000687415 |
| ENSMUSG00000005087 | 2 | Cd44 | protein_coding | 1.936389141 | 0 | -0.498217493 | 1.397863 | 3.006067 | 0.014659706 | 0.000766877 |
| ENSMUSG00000024253 | 17 | Dync2li1 | protein_coding | 1.924244933 | 0 | 0.134983131 | -0.889781 | 3.052832 | 0.047187386 | 0.003790581 |
| ENSMUSG00000018819 | 7 | Lsp1 | protein_coding | 1.919165727 | 0 | 0.187969374 | 0.610754 | 3.153445 | 1.68E-11 | 2.72E-15 |
| ENSMUSG00000047735 | 6 | Samd9l | protein_coding | 1.903119561 | 0 | -0.995982668 | 1.362458 | 3.205342 | 0.000446523 | 9.58204E-06 |
| ENSMUSG00000085111 | 10 | Ascl4 | protein_coding | 1.899016084 | 0 | -0.911591927 | -0.610863 | 3.041029 | 0.011979583 | 0.00056944 |
| ENSMUSG00000050711 | 1 | Scg2 | protein_coding | 1.879521662 | 0 | 2.848658398 | 4.185677 | 4.204894 | 0.005047864 | 0.000184354 |
| ENSMUSG00000051748 | 11 | Wfdc21 | protein_coding | 1.876234785 | 0 | -0.621607066 | 3.061403 | -2.461747 | 0.000345676 | 7.05404E-06 |
| ENSMUSG00000027712 | 3 | Anxa5 | protein_coding | 1.858962688 | 0 | 0.636789593 | 1.346439 | 2.551113 | 3.06894E-05 | 3.60E-07 |
| ENSMUSG00000052631 | 6 | Sh2d6 | protein_coding | 1.849344922 | 0 | 1.506442957 | 1.904673 | 3.514078 | 0.044077203 | 0.003440799 |
| ENSMUSG00000040907 | 7 | Atp1a3 | protein_coding | 1.847252672 | 0 | 0.898226006 | 2.085694 | 4.681409 | 0.000102738 | 1.55576E-06 |
| ENSMUSG00000003657 | 8 | Calb2 | protein_coding | 1.846606622 | 0 | 0.472680571 | 3.91177 | 4.05573 | 0.002520065 | 7.63223E-05 |
| ENSMUSG00000028402 | 4 | Mpdz | protein_coding | 1.842048275 | 0 | -0.247662898 | 2.245623 | 4.400459 | 0.003583261 | 0.000118098 |
| ENSMUSG00000032231 | 9 | Anxa2 | protein_coding | 1.841535957 | 0 | -0.875834907 | 1.870661 | 2.604779 | 0.017428105 | 0.000956859 |
| ENSMUSG00000024659 | 19 | Anxa1 | protein_coding | 1.81924572 | 0 | 0.342850856 | 3.304358 | -1.793764 | 0.006255499 | 0.000243655 |
| ENSMUSG00000027221 | 2 | Chst1 | protein_coding | 1.815490821 | 0 | 1.437993686 | 1.925774 | 3.818197 | 2.44623E-06 | 1.15E-08 |
| ENSMUSG00000029561 | 5 | Oasl2 | protein_coding | 1.795715342 | 0 | -2.173779349 | 1.554389 | 3.065798 | 4.22577E-06 | 2.41E-08 |
| ENSMUSG00000057315 | 5 | Arhgap24 | protein_coding | 1.790764884 | 0 | 0.574062879 | 0.687121 | 1.903462 | 0.000675332 | 1.58749E-05 |
| ENSMUSG00000027698 | 3 | Nceh1 | protein_coding | 1.773802764 | 0 | 0.41899876 | 0.633617 | 2.304464 | 1.70933E-06 | 6.09E-09 |
| ENSMUSG00000024044 | 17 | Epb41l3 | protein_coding | 1.755082238 | 0 | 0.735533746 | 1.432323 | 1.980963 | 4.7115E-05 | 6.18E-07 |
| ENSMUSG00000037148 | 8 | Arhgap10 | protein_coding | 1.72543893 | 0 | -1.319734555 | 1.49094 | 2.514594 | 0.030389223 | 0.001998222 |
| ENSMUSG00000035283 | 19 | Adrb1 | protein_coding | 1.711740414 | 0 | 0.870411304 | 1.001319 | 1.18719 | 4.71188E-05 | 6.22E-07 |
| ENSMUSG00000038576 | 1 | Susd4 | protein_coding | 1.694741153 | 0 | -0.336623544 | 3.143603 | 3.553417 | 0.013328406 | 0.00067241 |
| ENSMUSG00000045328 | 3 | Cenpe | protein_coding | 1.694135641 | 0 | 1.591471286 | 3.371563 | 3.703806 | 0.000585689 | 1.33747E-05 |
| ENSMUSG00000027276 | 2 | Jag1 | protein_coding | 1.683073011 | 0 | 0.71943573 | -0.075269 | 4.254852 | 0.013855695 | 0.000710972 |
| ENSMUSG00000026072 | 1 | Il1r1 | protein_coding | 1.673123303 | 0 | -2.119965163 | -1.188743 | 0.933983 | 0.028954707 | 0.001882795 |
| ENSMUSG00000001918 | 7 | Slc1a5 | protein_coding | 1.657608875 | 0 | -0.535577297 | 0.71844 | 2.414196 | 0.000367959 | 7.56835E-06 |
| ENSMUSG00000040710 | 1 | St8sia4 | protein_coding | 1.651808293 | 0 | 0.286232127 | 0.625584 | 0.386966 | 0.001717468 | 4.82599E-05 |
| ENSMUSG00000040528 | 11 | Milr1 | protein_coding | 1.648578416 | 0 | 1.016278025 | 1.772892 | 2.714059 | 0.0002754 | 5.32457E-06 |
| ENSMUSG00000020142 | 11 | Slc1a4 | protein_coding | 1.648463887 | 0 | 1.070609695 | 1.88399 | -2.067533 | 0.004192068 | 0.000145944 |
| ENSMUSG00000032258 | 9 | Lca5 | protein_coding | 1.62540187 | 0 | -1.581926459 | 1.109581 | 2.697122 | 0.027772435 | 0.001778929 |
| ENSMUSG00000002058 | 11 | Unc119 | protein_coding | 1.619491194 | 0 | 1.653796044 | 1.271929 | 7.48234 | 7.75474E-05 | 1.11778E-06 |
| ENSMUSG00000034855 | 5 | Cxcl10 | protein_coding | 1.618113008 | 0 | -0.606105827 | 3.036347 | 5.714292 | 5.49801E-05 | 7.52E-07 |
| ENSMUSG00000090223 | 16 | Pcp4 | protein_coding | 1.61112427 | 0 | 1.892558356 | 3.401235 | 2.444439 | 0.013275273 | 0.000668655 |
| ENSMUSG00000035356 | 16 | Nfkbiz | protein_coding | 1.605494279 | 0 | 0.708755375 | 1.342019 | 1.632144 | 0.002432255 | 7.28751E-05 |
| ENSMUSG00000043740 | 17 | B430306N03Rik | protein_coding | 1.602981139 | 0 | 0.800115623 | 1.586079 | 2.506014 | 0.007966472 | 0.000331587 |
| ENSMUSG00000032484 | 9 | Ngp | protein_coding | 1.596930891 | 0 | 0.709061081 | 3.281334 | -4.190131 | 0.000925047 | 2.29221E-05 |
| ENSMUSG00000113262 | 10 | Dohh | protein_coding | 1.591141677 | 0 | -1.269784203 | -0.890432 | 3.923102 | 0.006252671 | 0.000243038 |
| ENSMUSG00000020431 | 11 | Adcy1 | protein_coding | 1.588379871 | 0 | -0.859806731 | -3.135424 | 0.968 | 0.027640713 | 0.0017593 |
| ENSMUSG00000023495 | 9 | Pcbp4 | protein_coding | 1.580980794 | 0 | 2.261121692 | -1.576246 | 4.482905 | 0.001264893 | 3.35966E-05 |
| ENSMUSG00000027514 | 2 | Zbp1 | protein_coding | 1.579878847 | 0 | -1.087222155 | 2.380799 | 3.499346 | 0.002289585 | 6.7488E-05 |
| ENSMUSG00000023755 | 15 | Rhebl1 | protein_coding | 1.578393815 | 0 | 1.480187896 | -1.815545 | -0.842244 | 0.040235467 | 0.003033381 |

| | | | | | | | | | | |
|---|---|---|---|---|---|---|---|---|---|---|
| ENSMUSG00000020396 | 11 | Nefh | protein_coding | 1.576741291 | 0 | 0.057789566 | -0.83467 | 4.011005 | 0.005704513 | 0.000215265 |
| ENSMUSG00000041695 | 11 | Kcnj2 | protein_coding | 1.57662933 | 0 | 0.489156097 | 1.027172 | 1.546315 | 0.004129086 | 0.00014244 |
| ENSMUSG00000041488 | 19 | Stx3 | protein_coding | 1.559715346 | 0 | 0.394840878 | 0.502495 | 4.676043 | 0.000470714 | 1.02155E-05 |
| ENSMUSG00000043263 | 1 | Ifi209 | protein_coding | 1.553626858 | 0 | -0.775775013 | 0.683676 | 2.340976 | 7.24849E-05 | 1.03307E-06 |
| ENSMUSG00000028063 | 3 | Lmna | protein_coding | 1.550757124 | 0 | -0.478930262 | 1.156416 | 3.031863 | 0.013949393 | 0.000719553 |
| ENSMUSG00000035385 | 11 | Ccl2 | protein_coding | 1.523525246 | 0 | 0.665432576 | 1.767713 | 1.964214 | 0.008225206 | 0.000345693 |
| ENSMUSG00000019088 | X | Dnase1l1 | protein_coding | 1.522905714 | 0 | -0.1103039 | 0.211523 | 2.01654 | 0.001756637 | 4.95028E-05 |
| ENSMUSG00000041827 | 5 | Oasl1 | protein_coding | 1.47976849 | 0 | -0.781369304 | 1.201743 | 3.597087 | 0.032289886 | 0.002185954 |
| ENSMUSG00000043943 | 9 | Naalad2 | protein_coding | 1.43944121 | 0 | 0.472876973 | 1.272818 | 2.043925 | 3.06536E-06 | 1.49E-08 |
| ENSMUSG00000037242 | 4 | Clic4 | protein_coding | 1.425344747 | 0 | 0.28027084 | 1.357884 | 2.009177 | 0.039985127 | 0.002975652 |
| ENSMUSG00000073409 | 17 | H2-Q6 | protein_coding | 1.424454766 | 0 | -0.590746461 | 2.285191 | 2.683226 | 0.005930267 | 0.000227145 |
| ENSMUSG00000034452 | 9 | Slc24a1 | protein_coding | 1.419563854 | 0 | 0.473500905 | -0.035895 | 9.499346 | 1.67453E-05 | 1.52E-07 |
| ENSMUSG00000004891 | 3 | Nes | protein_coding | 1.402866462 | 0 | 0.068907885 | -0.429802 | 2.811808 | 0.001775513 | 5.01785E-05 |
| ENSMUSG00000030672 | 7 | Mylpf | protein_coding | 1.400308735 | 0 | 2.663944733 | 2.316952 | 4.215127 | 0.041582362 | 0.003178699 |
| ENSMUSG00000018930 | 11 | Ccl4 | protein_coding | 1.362406896 | 0 | 0.66604151 | 1.539092 | 1.835519 | 0.000575584 | 1.30816E-05 |
| ENSMUSG00000020846 | 11 | Rflnb | protein_coding | 1.362257587 | 0 | -0.336623544 | 3.452147 | 0.867041 | 0.036671215 | 0.00261916 |
| ENSMUSG00000000982 | 11 | Ccl3 | protein_coding | 1.35485491 | 0 | 0.880118146 | 1.736307 | 2.622812 | 0.000103311 | 1.58116E-06 |
| ENSMUSG00000038943 | 7 | Prc1 | protein_coding | 1.345504676 | 0 | 0.853059845 | 3.708975 | 4.023668 | 0.005751945 | 0.000218452 |
| ENSMUSG00000030717 | 7 | Nupr1 | protein_coding | 1.342441225 | 0 | 0.188954122 | 0.858082 | 1.035984 | 0.012254346 | 0.000598378 |
| ENSMUSG00000026222 | 1 | Sp100 | protein_coding | 1.337678102 | 0 | -0.799619011 | 1.019911 | 1.987718 | 3.78093E-05 | 4.69E-07 |
| ENSMUSG00000041736 | 15 | Tspo | protein_coding | 1.331900758 | 0 | 0.489616708 | 1.040222 | 2.412664 | 1.51367E-05 | 1.32E-07 |
| ENSMUSG00000027715 | 3 | Ccna2 | protein_coding | 1.318746837 | 0 | 1.81031138 | 1.846666 | 4.22765 | 0.035778373 | 0.002519828 |
| ENSMUSG00000035352 | 11 | Ccl12 | protein_coding | 1.316764421 | 0 | -0.234321432 | 2.163093 | 3.337355 | 1.05E-10 | 2.56E-14 |
| ENSMUSG00000028494 | 4 | Plin2 | protein_coding | 1.315340576 | 0 | 0.231548819 | 0.278026 | 1.242398 | 0.000549492 | 1.22811E-05 |
| ENSMUSG00000024378 | 18 | Stard4 | protein_coding | 1.311640822 | 0 | -1.442708415 | 1.652242 | 2.541265 | 0.010946847 | 0.000503507 |
| ENSMUSG00000034714 | 11 | Ttyh2 | protein_coding | 1.288416509 | 0 | 0.976665072 | 0.228794 | 2.171248 | 0.036738435 | 0.002629911 |
| ENSMUSG00000025395 | 10 | Prim1 | protein_coding | 1.278898731 | 0 | 0.740079348 | 1.656992 | 3.722816 | 0.024657731 | 0.001491564 |
| ENSMUSG00000031304 | X | Il2rg | protein_coding | 1.277908709 | 0 | 0.086697919 | 2.671912 | 3.134673 | 0.000515365 | 1.13515E-05 |
| ENSMUSG00000027849 | 3 | Syt6 | protein_coding | 1.276505245 | 0 | 2.797950603 | 3.030698 | -0.857593 | 0.039544139 | 0.002933228 |
| ENSMUSG00000000386 | 16 | Mx1 | protein_coding | 1.27431789 | 0 | -2.035413201 | 1.269264 | 1.883345 | 0.000674294 | 1.57803E-05 |
| ENSMUSG00000031004 | 7 | Mki67 | protein_coding | 1.254117841 | 0 | 2.866656627 | 4.564976 | 3.366079 | 0.037617181 | 0.00270919 |
| ENSMUSG00000038357 | 9 | Camp | protein_coding | 1.248168749 | 0 | 0.399393367 | 3.432306 | -4.053249 | 0.011052204 | 0.000510143 |
| ENSMUSG00000069516 | 10 | Lyz2 | protein_coding | 1.245187206 | 0 | 0.298421084 | 1.860094 | 1.192964 | 1.13316E-05 | 9.08E-08 |
| ENSMUSG00000056054 | 3 | S100a8 | protein_coding | 1.242062331 | 0 | 0.742322929 | 3.466899 | -4.353037 | 0.011269934 | 0.000526092 |
| ENSMUSG00000019876 | 10 | Pkib | protein_coding | 1.238783605 | 0 | 0.489241208 | 0.633901 | 1.739605 | 0.000210987 | 3.79021E-06 |
| ENSMUSG00000059326 | 19 | Csf2ra | protein_coding | 1.23037946 | 0 | 0.42888742 | 0.444896 | 1.632927 | 2.14044E-06 | 8.52E-09 |
| ENSMUSG00000038059 | 18 | Smim3 | protein_coding | 1.229654445 | 0 | -0.15772389 | 0.852077 | 2.421568 | 9.11088E-06 | 6.59E-08 |
| ENSMUSG00000022021 | 14 | Diaph3 | protein_coding | 1.204297277 | 0 | -1.08971953 | 2.170271 | 3.578938 | 0.008026635 | 0.000334741 |
| ENSMUSG00000024401 | 17 | Tnf | protein_coding | 1.203019381 | 0 | -0.08397666 | 0.722149 | 3.369352 | 0.025196717 | 0.001535915 |
| ENSMUSG00000029663 | 6 | Gngt1 | protein_coding | 1.19955318 | 0 | 1.074736081 | 0.746352 | 10.86395 | 0.00043447 | 9.2882E-06 |
| ENSMUSG00000030283 | 6 | St8sia1 | protein_coding | 1.199488346 | 0 | -1.296660195 | -0.530976 | 2.651546 | 0.021614035 | 0.001270693 |
| ENSMUSG00000032841 | 2 | Prr5l | protein_coding | 1.196516468 | 0 | -0.566318992 | 0.298529 | 1.264614 | 2.15781E-05 | 2.22E-07 |
| ENSMUSG00000044703 | 14 | Phf11a | protein_coding | 1.172637684 | 0 | 1.063429536 | 2.702794 | 6.534529 | 3.48396E-06 | 1.72E-08 |
| ENSMUSG00000034891 | 13 | Sncb | protein_coding | 1.171213235 | 0 | -1.303840829 | 1.145527 | 3.753211 | 0.007612838 | 0.000310703 |
| ENSMUSG00000001128 | X | Cfp | protein_coding | 1.169955319 | 0 | 0.956314205 | 1.359501 | -1.594824 | 0.01459084 | 0.000757367 |
| ENSMUSG00000029636 | 5 | Wasf3 | protein_coding | 1.168891911 | 0 | 0.893329614 | 2.503265 | 4.120123 | 0.02136753 | 0.001254471 |
| ENSMUSG00000026014 | 1 | Raph1 | protein_coding | 1.160787117 | 0 | 0.533057029 | 0.26675 | -0.471763 | 0.004927687 | 0.000178908 |
| ENSMUSG00000027306 | 2 | Nusap1 | protein_coding | 1.158862315 | 0 | 0.952268072 | 4.714937 | 3.739104 | 0.002995861 | 9.48564E-05 |
| ENSMUSG00000017754 | 2 | Pltp | protein_coding | 1.152724005 | 0 | 0.073749168 | 1.495796 | -1.783151 | 0.013358064 | 0.000674988 |
| ENSMUSG00000073491 | 1 | Ifi213 | protein_coding | 1.143327926 | 0 | 0.010841621 | 2.231345 | 2.40534 | 0.008225206 | 0.000347685 |
| ENSMUSG00000072620 | 11 | Slfn2 | protein_coding | 1.141011207 | 0 | -0.052820031 | 1.034402 | 1.530552 | 2.37421E-06 | 1.06E-08 |
| ENSMUSG00000015314 | 1 | Slamf6 | protein_coding | 1.139540744 | 0 | 0.015942786 | 0.845531 | 1.238045 | 0.000521444 | 1.15276E-05 |
| ENSMUSG00000029373 | 5 | Pf4 | protein_coding | 1.133953422 | 0 | 0.671735312 | 1.615624 | -0.790043 | 0.010871996 | 0.000498303 |
| ENSMUSG00000017716 | 11 | Birc5 | protein_coding | 1.132921151 | 0 | 0.649380466 | 3.305297 | 4.453524 | 0.020781576 | 0.001201558 |
| ENSMUSG00000013523 | 2 | Bcas1 | protein_coding | 1.121880731 | 0 | 1.69157353 | 0.719229 | -2.329324 | 0.015599758 | 0.000831212 |
| ENSMUSG00000054404 | 11 | Slfn5 | protein_coding | 1.119960649 | 0 | -0.357682101 | 1.572261 | 2.022226 | 1.20115E-05 | 9.82E-08 |
| ENSMUSG00000004707 | 1 | Ly9 | protein_coding | 1.115620493 | 0 | -1.315567587 | 0.91389 | 1.686722 | 0.000675332 | 1.5914E-05 |
| ENSMUSG00000036768 | 9 | Kif15 | protein_coding | 1.110268957 | 0 | -1.078341457 | 1.88435 | 1.204856 | 0.016962123 | 0.000927155 |
| ENSMUSG00000008845 | 6 | Cd163 | protein_coding | 1.106020813 | 0 | 0.39783313 | 1.580175 | -2.090052 | 0.012553548 | 0.000616038 |
| ENSMUSG00000021986 | 14 | Amer2 | protein_coding | 1.098199048 | 0 | -0.736851935 | -0.033588 | 1.609935 | 0.04014065 | 0.003004132 |
| ENSMUSG00000062488 | 19 | Ifit3b | protein_coding | 1.085227233 | 0 | -2.054939488 | 1.574942 | 3.565212 | 0.000113031 | 1.76654E-06 |
| ENSMUSG00000039942 | 15 | Ptger4 | protein_coding | 1.084989857 | 0 | 2.175186386 | 4.036334 | 2.534857 | 0.03248338 | 0.002206944 |
| ENSMUSG00000024197 | 17 | Plin3 | protein_coding | 1.076012878 | 0 | 0.492085417 | 0.624596 | 1.774328 | 0.002964727 | 9.36305E-05 |
| ENSMUSG00000056671 | 18 | Prelid2 | protein_coding | 1.071122658 | 0 | 0.94991868 | 0.568839 | 1.698104 | 0.030985782 | 0.002049995 |
| ENSMUSG00000025743 | 4 | Sdc3 | protein_coding | 1.063661022 | 0 | -0.457964997 | 1.114302 | 1.305996 | 0.000966647 | 2.41877E-05 |
| ENSMUSG00000070570 | 7 | Slc17a7 | protein_coding | 1.057448925 | 0 | 0.225872333 | -0.171755 | 4.864742 | 0.000450499 | 9.70383E-06 |
| ENSMUSG00000000318 | 11 | Clec10a | protein_coding | 1.047857592 | 0 | 0.084126464 | 1.185873 | -3.712845 | 0.000185852 | 3.23574E-06 |
| ENSMUSG00000028488 | 4 | Sh3gl2 | protein_coding | 1.04522904 | 0 | 1.539277468 | 2.033453 | 4.782134 | 0.011269934 | 0.000525332 |
| ENSMUSG00000074896 | 19 | Ifit3 | protein_coding | 1.045014111 | 0 | -0.872782344 | 0.181265 | 2.152417 | 0.000164785 | 2.77555E-06 |
| ENSMUSG00000028970 | 5 | Abcb1b | protein_coding | 1.03967233 | 0 | 0.30784721 | 0.394701 | 0.539063 | 0.01189262 | 0.00056338 |
| ENSMUSG00000040751 | 5 | Lat2 | protein_coding | 1.038443376 | 0 | 0.44965854 | 0.376814 | 1.846815 | 8.70E-10 | 3.52E-13 |
| ENSMUSG00000038775 | 9 | Vill | protein_coding | 1.037573583 | 0 | 3.767988328 | 1.982233 | 4.528463 | 0.002734908 | 8.41578E-05 |
| ENSMUSG00000091649 | 14 | Phf11b | protein_coding | 1.035347938 | 0 | 0.117291392 | 1.240817 | 2.661741 | 4.77804E-06 | 2.82E-08 |
| ENSMUSG00000004952 | 5 | Rasa4 | protein_coding | 1.023748698 | 0 | 0.558496401 | 0.649794 | 1.229783 | 0.031037555 | 0.002065987 |

| | | | | | | | | | | |
|---|---|---|---|---|---|---|---|---|---|---|
| ENSMUSG00000029163 | 5 | Emilin1 | protein_coding | 1.012446138 | 0 | 1.184717129 | 1.978568 | 3.594512 | 0.022650649 | 0.001349978 |
| ENSMUSG00000044199 | 10 | S1pr4 | protein_coding | 1.001386124 | 0 | 1.259438073 | 2.650417 | -1.857729 | 0.032677161 | 0.002222756 |
| ENSMUSG00000022844 | 16 | Pdia5 | protein_coding | 0.989686152 | 0 | 0.706124757 | 0.951925 | 2.148605 | 0.047395842 | 0.003822679 |
| ENSMUSG00000031328 | X | Flna | protein_coding | 0.96546977 | 0 | -0.37630059 | 0.07258 | 0.81951 | 0.002377874 | 7.08606E-05 |
| ENSMUSG00000022351 | 15 | Sqle | protein_coding | 0.962017494 | 0 | 1.251290581 | -0.206231 | 4.579106 | 0.001351896 | 3.61265E-05 |
| ENSMUSG00000038335 | 11 | Tsr1 | protein_coding | 0.959446471 | 0 | 1.293452735 | -0.407573 | 1.088331 | 0.040233467 | 0.003026714 |
| ENSMUSG00000036599 | 5 | Chst12 | protein_coding | 0.95822489 | 0 | -0.135296459 | 0.031016 | 1.201191 | 0.001656668 | 4.62831E-05 |
| ENSMUSG00000025492 | 7 | Ifitm3 | protein_coding | 0.957247399 | 0 | -0.547974772 | 2.031251 | 2.568357 | 6.95E-07 | 2.03E-09 |
| ENSMUSG00000018983 | 4 | E2f2 | protein_coding | 0.94309989 | 0 | 0.938050816 | 1.931322 | 3.454525 | 0.049487968 | 0.004023477 |
| ENSMUSG00000056737 | 6 | Capg | protein_coding | 0.931894598 | 0 | 0.584147386 | -0.374712 | 2.846975 | 4.71E-07 | 1.14E-09 |
| ENSMUSG00000036533 | 17 | Cdc42ep3 | protein_coding | 0.925134809 | 0 | -0.338623754 | 0.483397 | 1.386804 | 0.008639547 | 0.000367298 |
| ENSMUSG00000025498 | 7 | Irf7 | protein_coding | 0.919813938 | 0 | -0.341467784 | 1.076964 | 1.932754 | 0.047180577 | 0.003786214 |
| ENSMUSG00000040483 | 11 | Xaf1 | protein_coding | 0.90722977 | 0 | -0.301418616 | 0.952366 | 0.878173 | 0.006533416 | 0.000257654 |
| ENSMUSG00000070327 | 11 | Rnf213 | protein_coding | 0.900603873 | 0 | 0.125632023 | 0.860365 | 1.045643 | 0.031021216 | 0.002059658 |
| ENSMUSG00000038679 | 15 | Trps1 | protein_coding | 0.895456622 | 0 | -0.754901621 | -0.156629 | 1.629596 | 0.002453157 | 7.37E-05 |
| ENSMUSG00000037126 | 19 | Psd | protein_coding | 0.893435923 | 0 | 3.391396585 | 0.091544 | 2.275825 | 0.049487968 | 0.004023151 |
| ENSMUSG00000051456 | 13 | Hspb3 | protein_coding | 0.86923801 | 0 | 0.9695782 | 1.270025 | 1.522655 | 0.036102751 | 0.002561018 |
| ENSMUSG00000043510 | 5 | Hscb | protein_coding | 0.854080463 | 0 | 1.36003974 | 1.601515 | 1.640008 | 0.027772435 | 0.001777194 |
| ENSMUSG00000041329 | 11 | Atp1b2 | protein_coding | 0.852691313 | 0 | -0.00270631 | 1.715887 | 4.780074 | 0.008936393 | 0.000387154 |
| ENSMUSG00000030142 | 6 | Clec4e | protein_coding | 0.845707485 | 0 | 0.611680164 | -0.035895 | 4.45315 | 0.011464118 | 0.00053907 |
| ENSMUSG00000054580 | 2 | Pla2r1 | protein_coding | 0.845707485 | 0 | -0.336623544 | 0.578671 | 4.639159 | 0.0002754 | 5.35233E-06 |
| ENSMUSG00000024518 | 18 | Rax | protein_coding | 0.845707485 | 0 | -0.336623544 | -0.035895 | 4.765243 | 0.000150909 | 2.52961E-06 |
| ENSMUSG00000056055 | 1 | Sag | protein_coding | 0.832981308 | 0 | 0.299210171 | 0.090119 | 5.63379 | 3.59457E-05 | 4.40E-07 |
| ENSMUSG00000020601 | 12 | Trib2 | protein_coding | 0.830974957 | 0 | 0.576589904 | 0.961086 | 4.546102 | 0.012513423 | 0.000613055 |
| ENSMUSG00000022995 | 1 | Enah | protein_coding | 0.811517035 | 0 | 1.693780168 | 1.000347 | 3.454092 | 0.040966922 | 0.003113112 |
| ENSMUSG00000024672 | 19 | Ms4a7 | protein_coding | 0.811326776 | 0 | 0.115513042 | 1.599869 | 1.186973 | 0.020156626 | 0.001155574 |
| ENSMUSG00000068245 | 14 | Phf11d | protein_coding | 0.803753403 | 0 | 0.058266484 | 0.406721 | 2.100324 | 2.53726E-05 | 2.76E-07 |
| ENSMUSG00000073411 | 17 | H2-D1 | protein_coding | 0.79872045 | 0 | -0.235613737 | 1.016082 | 1.902674 | 1.89E-08 | 1.67E-11 |
| ENSMUSG00000029055 | 4 | Plch2 | protein_coding | 0.780801602 | 0 | -0.336623544 | 0.902737 | 4.047878 | 0.017575594 | 0.000968145 |
| ENSMUSG00000002985 | 7 | Apoe | protein_coding | 0.775731977 | 0 | 0.254956063 | 2.028773 | 1.254863 | 3.93E-08 | 5.41E-11 |
| ENSMUSG00000033355 | 16 | Rtp4 | protein_coding | 0.767389487 | 0 | -0.650812285 | 1.15184 | 1.862474 | 0.000147595 | 2.45016E-06 |
| ENSMUSG00000020053 | 10 | Igf1 | protein_coding | 0.762182846 | 0 | 0.611554142 | 1.72064 | 1.021184 | 0.045656676 | 0.003589978 |
| ENSMUSG00000068876 | 3 | Cgn | protein_coding | 0.757893951 | 0 | 0.665158554 | 0.900869 | 3.754096 | 0.033650566 | 0.002316218 |
| ENSMUSG00000022150 | 15 | Dab2 | protein_coding | 0.756335371 | 0 | 0.047595562 | 1.186844 | -0.741707 | 0.000103311 | 1.57935E-06 |
| ENSMUSG00000030577 | 7 | Cd22 | protein_coding | 0.751934928 | 0 | -0.574999916 | 1.744083 | 0.647736 | 0.038853506 | 0.002869414 |
| ENSMUSG00000024501 | 18 | Dpysl3 | protein_coding | 0.751275874 | 0 | -0.146562997 | -0.319623 | 4.943778 | 0.000955255 | 2.37479E-05 |
| ENSMUSG00000027168 | 2 | Pax6 | protein_coding | 0.747187761 | 0 | 0.825267536 | 1.903524 | 4.751747 | 0.011874403 | 0.000561556 |
| ENSMUSG00000029413 | 5 | Naaa | protein_coding | 0.737629678 | 0 | -0.240407445 | 0.324868 | 1.157233 | 0.000640874 | 1.48425E-05 |
| ENSMUSG00000020895 | 11 | Tmem107 | protein_coding | 0.734782622 | 0 | -0.475450239 | 0.345628 | 3.655059 | 0.013275273 | 0.000668651 |
| ENSMUSG00000049353 | 1 | Rd3 | protein_coding | 0.731105934 | 0 | 0.98568139 | 0.37436 | 5.728308 | 0.002134152 | 6.16967E-05 |
| ENSMUSG00000037851 | 13 | Iars | protein_coding | 0.722404098 | 0 | 0.451572879 | 1.060658 | 0.05093 | 0.020676141 | 0.001193788 |
| ENSMUSG00000025150 | 11 | Cbr2 | protein_coding | 0.721910184 | 0 | -0.249071652 | 0.838876 | -2.945142 | 0.02982775 | 0.001949226 |
| ENSMUSG00000027530 | 3 | Fabp12 | protein_coding | 0.713540544 | 0 | 0.05678378 | -0.035895 | 6.427615 | 0.000227825 | 4.22479E-06 |
| ENSMUSG00000028459 | 4 | Cd72 | protein_coding | 0.70542307 | 0 | -0.439783607 | 0.813552 | 3.41598 | 2.41E-07 | 5.26E-10 |
| ENSMUSG00000102037 | 9 | Bcl2a1a | protein_coding | 0.702951996 | 0 | -0.058413094 | 0.600727 | 1.672781 | 0.02764128 | 0.001763813 |
| ENSMUSG00000043008 | 16 | Klhl6 | protein_coding | 0.698811276 | 0 | 0.385204044 | 0.593132 | 1.117261 | 0.004192068 | 0.000146272 |
| ENSMUSG00000045932 | 19 | Ifit2 | protein_coding | 0.684880985 | 0 | -1.154984955 | 1.888731 | 2.6972 | 7.76915E-06 | 5.10E-08 |
| ENSMUSG00000000682 | 4 | Cd52 | protein_coding | 0.681097982 | 0 | 0.050644716 | 0.863661 | 2.016717 | 9.51E-08 | 1.74E-10 |
| ENSMUSG00000041607 | 18 | Mbp | protein_coding | 0.677748461 | 0 | 0.99818058 | 0.05094 | -1.633722 | 0.002913973 | 9.08478E-05 |
| ENSMUSG00000021360 | 13 | Gcnt2 | protein_coding | 0.672026737 | 0 | -0.406356133 | 0.414148 | 2.179134 | 3.7748E-06 | 2.04E-08 |
| ENSMUSG00000019823 | 10 | Mical1 | protein_coding | 0.668493104 | 0 | -0.015720788 | 0.35978 | 1.641319 | 0.035778373 | 0.002520624 |
| ENSMUSG00000067220 | 5 | Cnga1 | protein_coding | 0.656291893 | 0 | -0.512184051 | 0.188962 | 7.805889 | 1.04152E-05 | 8.05E-08 |
| ENSMUSG00000068129 | 2 | Cst7 | protein_coding | 0.654528805 | 0 | 0.289194958 | 2.325981 | 2.062429 | 0.000115854 | 1.8388E-06 |
| ENSMUSG00000031490 | 8 | Eif4ebp1 | protein_coding | 0.652354812 | 0 | -0.449328876 | 0.417196 | 0.542115 | 0.003026349 | 9.6802E-05 |
| ENSMUSG00000079339 | 19 | Ifit1bl1 | protein_coding | 0.649709867 | 0 | 0.229455678 | 1.078606 | 3.48942 | 0.040966922 | 0.003115065 |
| ENSMUSG00000047798 | 11 | Cd300lf | protein_coding | 0.642235207 | 0 | -0.013678992 | 1.175455 | 2.449149 | 0.038508643 | 0.002813072 |
| ENSMUSG00000020634 | 12 | Ubxn2a | protein_coding | 0.634250631 | 0 | -0.749290646 | -0.719656 | -0.454357 | 0.016113707 | 0.000870341 |
| ENSMUSG00000042111 | 1 | Ccdc115 | protein_coding | 0.633556021 | 0 | -0.032932861 | -0.008293 | 1.134484 | 0.009187008 | 0.000405451 |
| ENSMUSG00000021624 | 13 | Cd180 | protein_coding | 0.627737375 | 0 | 0.317508014 | 0.409941 | 1.165195 | 0.000138151 | 2.25981E-06 |
| ENSMUSG00000023078 | 5 | Cxcl13 | protein_coding | 0.627406106 | 0 | 0.425068795 | 2.545504 | 4.326694 | 0.034809016 | 0.002429782 |
| ENSMUSG00000026548 | 1 | Slamf9 | protein_coding | 0.622342982 | 0 | -0.337096275 | 0.112451 | 1.863134 | 1.89E-08 | 1.69E-11 |
| ENSMUSG00000039347 | 6 | Atp6v0e2 | protein_coding | 0.613337095 | 0 | 2.558977729 | 0.215389 | 4.53574 | 0.013079023 | 0.000653475 |
| ENSMUSG00000021880 | 14 | Rnase6 | protein_coding | 0.60285831 | 0 | -0.233127751 | 0.180905 | 1.542308 | 0.001537876 | 4.20927E-05 |
| ENSMUSG00000037979 | 5 | Ccdc92 | protein_coding | 0.602603786 | 0 | 0.229455678 | 0.705757 | 4.294802 | 0.006508351 | 0.000256139 |
| ENSMUSG00000028654 | 4 | Mycl | protein_coding | 0.595890382 | 0 | 0.408037068 | 0.393462 | -2.680879 | 0.013499696 | 0.000683238 |
| ENSMUSG00000034361 | 8 | Cpne2 | protein_coding | 0.593933413 | 0 | 0.750443799 | 1.001001 | -2.275527 | 0.041502018 | 0.003169196 |
| ENSMUSG00000021779 | 14 | Thrb | protein_coding | 0.587232659 | 0 | -1.603773617 | -1.298801 | 2.629909 | 0.007528323 | 0.000305425 |
| ENSMUSG00000058966 | 7 | Fam57b | protein_coding | 0.586130222 | 0 | 1.01052245 | -0.035895 | 7.576801 | 1.66829E-05 | 1.50E-07 |
| ENSMUSG00000002885 | 8 | Adgre5 | protein_coding | 0.583694419 | 0 | 0.752797055 | 1.917685 | 3.660748 | 0.038801406 | 0.002854267 |
| ENSMUSG00000073418 | 17 | C4b | protein_coding | 0.582579793 | 0 | 0.429990026 | 2.133183 | 1.470262 | 0.038501152 | 0.002809097 |
| ENSMUSG00000051906 | 8 | Cd209f | protein_coding | 0.577309024 | 0 | 0.225057646 | 1.574848 | -3.502502 | 0.000220894 | 4.06048E-06 |
| ENSMUSG00000030560 | 7 | Ctsc | protein_coding | 0.576015024 | 0 | -0.185359303 | 0.237967 | 0.964521 | 1.28663E-05 | 1.08E-07 |
| ENSMUSG00000031198 | X | Fundc2 | protein_coding | 0.574865743 | 0 | 0.79297219 | 1.046586 | 1.640004 | 0.025274875 | 0.001547236 |
| ENSMUSG00000058818 | 7 | Pirb | protein_coding | 0.573465051 | 0 | -0.329036235 | -0.355808 | 1.309689 | 0.000117611 | 1.88358E-06 |

| | | | | | | | | | | |
|---|---|---|---|---|---|---|---|---|---|---|
| ENSMUSG00000021730 | 13 | Hcn1 | protein_coding | 0.569656657 | 0 | 0.06097877 | 0.758548 | 5.929369 | 2.13098E-05 | 2.14E-07 |
| ENSMUSG00000020131 | 10 | Pcsk4 | protein_coding | 0.55883006 | 0 | -1.026164137 | -0.11087 | 3.471292 | 0.002941425 | 9.23119E-05 |
| ENSMUSG00000028977 | 4 | Casz1 | protein_coding | 0.551881074 | 0 | -0.945855053 | 0.963034 | 2.6524 | 0.002253834 | 6.62517E-05 |
| ENSMUSG00000029410 | 5 | Ppef2 | protein_coding | 0.545003742 | 0 | 0.613414105 | -0.052815 | 3.750621 | 0.046148204 | 0.003651048 |
| ENSMUSG00000027199 | 2 | Gatm | protein_coding | 0.530203572 | 0 | -0.252825922 | 0.237445 | 0.924041 | 2.13098E-05 | 2.13E-07 |
| ENSMUSG00000075602 | 15 | Ly6a | protein_coding | 0.520683348 | 0 | -0.152146786 | 1.405857 | 1.789074 | 0.022622803 | 0.001344654 |
| ENSMUSG00000034459 | 19 | Ifit1 | protein_coding | 0.517675269 | 0 | -0.121120074 | 2.663941 | 2.753053 | 0.001513146 | 4.11707E-05 |
| ENSMUSG00000029816 | 6 | Gpnmb | protein_coding | 0.515763083 | 0 | 0.05678378 | 0.578671 | 7.097569 | 7.52E-08 | 1.16E-10 |
| ENSMUSG00000049422 | 10 | Chchd10 | protein_coding | 0.508654783 | 0 | 0.083181575 | 1.458597 | 2.716231 | 0.021364278 | 0.00125255 |
| ENSMUSG00000016283 | 17 | H2-M2 | protein_coding | 0.507835985 | 0 | 0.05678378 | 0.578671 | 4.62052 | 0.000394679 | 8.27774E-06 |
| ENSMUSG00000026958 | 2 | Dpp7 | protein_coding | 0.507202354 | 0 | -0.448868034 | 0.114564 | 0.676876 | 0.037258493 | 0.002673174 |
| ENSMUSG00000035504 | 10 | Reep6 | protein_coding | 0.499069458 | 0 | 1.010623124 | 0.67103 | 8.503833 | 0.000147595 | 2.44721E-06 |
| ENSMUSG00000025927 | 1 | Tfap2b | protein_coding | 0.4958319 | 0 | -0.336623544 | 0.632164 | 3.725436 | 0.021015415 | 0.001217596 |
| ENSMUSG00000025888 | 9 | Casp1 | protein_coding | 0.47622703 | 0 | -0.287461567 | 0.318614 | 0.776138 | 0.04622608 | 0.003664696 |
| ENSMUSG00000024053 | 17 | Emilin2 | protein_coding | 0.474175159 | 0 | -0.911591927 | 2.925305 | 2.256392 | 0.026988906 | 0.001700329 |
| ENSMUSG00000024661 | 19 | Fth1 | protein_coding | 0.472328094 | 0 | 0.033033417 | 0.557483 | 1.562967 | 3.59E-07 | 8.44E-10 |
| ENSMUSG00000044339 | 5 | Alkbh2 | protein_coding | 0.458409093 | 0 | -0.013422332 | -0.185563 | 1.934817 | 0.002082491 | 5.9743E-05 |
| ENSMUSG00000039109 | 13 | F13a1 | protein_coding | 0.45541578 | 0 | -0.203701956 | 0.868543 | -2.655849 | 0.004152483 | 0.000143583 |
| ENSMUSG00000021123 | 12 | Rdh12 | protein_coding | 0.454754776 | 0 | 0.142468866 | -1.053546 | 5.34789 | 0.002253834 | 6.60369E-05 |
| ENSMUSG00000040265 | 1 | Dnm3 | protein_coding | 0.451551527 | 0 | -0.413348156 | -0.08225 | 2.091985 | 0.041607518 | 0.003183991 |
| ENSMUSG00000021423 | 13 | Ly86 | protein_coding | 0.427764806 | 0 | 0.313323432 | 0.499933 | 1.09809 | 3.60653E-06 | 1.84E-08 |
| ENSMUSG00000022090 | 14 | Pdlim2 | protein_coding | 0.427433648 | 0 | 0.548745403 | 0.784068 | 1.307147 | 0.044077203 | 0.003440248 |
| ENSMUSG00000024736 | 19 | Tmem132a | protein_coding | 0.417513081 | 0 | -0.354265006 | 1.215015 | -2.901907 | 0.019548391 | 0.00110493 |
| ENSMUSG00000030760 | 7 | Acer3 | protein_coding | 0.415399633 | 0 | 0.225065175 | 0.251045 | 1.144731 | 4.22577E-06 | 2.37E-08 |
| ENSMUSG00000019970 | 10 | Sgk1 | protein_coding | 0.411753542 | 0 | 0.357618208 | 0.000228 | 1.200727 | 1.32304E-06 | 4.50E-09 |
| ENSMUSG00000021263 | 12 | Degs2 | protein_coding | 0.408667672 | 0 | 2.191634148 | 3.485678 | -0.062516 | 0.01275594 | 0.000632167 |
| ENSMUSG00000029657 | 5 | Hsph1 | protein_coding | 0.404573921 | 0 | -0.216568392 | -0.475104 | -1.17204 | 0.000966647 | 2.41691E-05 |
| ENSMUSG00000001023 | 3 | S100a5 | protein_coding | 0.394533205 | 0 | 1.313111111 | 7.936736 | 2.081891 | 2.22048E-05 | 2.34E-07 |
| ENSMUSG00000027995 | 3 | Tlr2 | protein_coding | 0.393167344 | 0 | -0.281924404 | 0.438684 | 0.857982 | 0.001070356 | 2.78228E-05 |
| ENSMUSG00000020649 | 12 | Rrm2 | protein_coding | 0.375796659 | 0 | 1.771724012 | 4.032948 | 2.653822 | 0.015355484 | 0.000811979 |
| ENSMUSG00000034837 | 9 | Gnat1 | protein_coding | 0.374339491 | 0 | 0.58160659 | 0.650342 | 10.82664 | 0.00035538 | 7.28084E-06 |
| ENSMUSG00000027506 | 3 | Tpd52 | protein_coding | 0.370471123 | 0 | -0.286318917 | 0.330052 | 0.723325 | 0.004496535 | 0.000160942 |
| ENSMUSG00000059089 | 1 | Fcgr4 | protein_coding | 0.349477778 | 0 | -0.17116871 | 1.120745 | 2.748538 | 6.85E-07 | 1.84E-09 |
| ENSMUSG00000030055 | 6 | Rab43 | protein_coding | 0.346020709 | 0 | -0.573975852 | -0.909436 | -0.163881 | 0.041985595 | 0.003229923 |
| ENSMUSG00000019889 | 10 | Ptprk | protein_coding | 0.339937622 | 0 | -1.132971015 | 2.888314 | 0.877121 | 0.045997207 | 0.003635377 |
| ENSMUSG00000070354 | 11 | Evi2 | protein_coding | 0.33963025 | 0 | -1.411684819 | -0.236415 | 1.320355 | 0.048683986 | 0.003930515 |
| ENSMUSG00000045502 | 5 | Hcar2 | protein_coding | 0.336676113 | 0 | 1.039856169 | 1.058514 | 3.845916 | 0.012254346 | 0.000597837 |
| ENSMUSG00000040950 | 11 | Mgl2 | protein_coding | 0.327123415 | 0 | -0.016004941 | 0.460365 | -3.38056 | 0.009478275 | 0.000419841 |
| ENSMUSG00000015355 | 1 | Cd48 | protein_coding | 0.318499186 | 0 | 0.034334065 | 0.222058 | 1.40397 | 5.23807E-05 | 7.08E-07 |
| ENSMUSG00000033032 | 18 | Afap1l1 | protein_coding | 0.312568389 | 0 | 0.215036902 | -0.072567 | -1.598096 | 0.005860762 | 0.000223534 |
| ENSMUSG00000063234 | 15 | Gpr84 | protein_coding | 0.312460588 | 0 | 0.282537212 | 0.631659 | 1.72591 | 3.60653E-06 | 1.82E-08 |
| ENSMUSG00000093661 | 6 | Eif4e3 | protein_coding | 0.310241436 | 0 | -0.483744204 | -0.048378 | 0.528521 | 0.04255958 | 0.003291311 |
| ENSMUSG00000061232 | 17 | H2-K1 | protein_coding | 0.301667965 | 0 | -0.521808484 | 0.573946 | 1.046261 | 6.15E-07 | 1.54E-09 |
| ENSMUSG00000043687 | 8 | 1190005I06Rik | protein_coding | 0.297267829 | 0 | 1.628724272 | -0.035895 | 3.82778 | 0.033769094 | 0.002329846 |
| ENSMUSG00000044258 | 13 | Ctla2a | protein_coding | 0.297267829 | 0 | -0.336623544 | 1.144864 | 3.770672 | 0.009601266 | 0.000426747 |
| ENSMUSG00000040714 | 7 | Klc3 | protein_coding | 0.297267829 | 0 | -0.336623544 | -0.035895 | 5.242993 | 7.76915E-06 | 5.16E-08 |
| ENSMUSG00000021804 | 14 | Rgr | protein_coding | 0.297267829 | 0 | -0.336623544 | -0.035895 | 4.180454 | 0.002134152 | 6.15353E-05 |
| ENSMUSG00000091472 | 14 | Gm3739 | protein_coding | 0.295754798 | 0 | 0.330343222 | -0.308864 | -1.32014 | 0.031021216 | 0.002059875 |
| ENSMUSG00000024294 | 18 | Mib1 | protein_coding | 0.28480721 | 0 | 0.107915895 | 0.187983 | -0.914604 | 0.020156626 | 0.001156628 |
| ENSMUSG00000024140 | 17 | Epas1 | protein_coding | 0.284393185 | 0 | 1.678022174 | 1.277003 | 3.785984 | 0.015959245 | 0.000856829 |
| ENSMUSG00000030107 | 6 | Usp18 | protein_coding | 0.27746345 | 0 | -0.829992592 | 1.157908 | 1.863214 | 0.001231781 | 3.24179E-05 |
| ENSMUSG00000032193 | 9 | Ldlr | protein_coding | 0.273286774 | 0 | -0.242299548 | -0.086279 | 2.261462 | 1.16589E-05 | 9.44E-08 |
| ENSMUSG00000030701 | 7 | Plekhb1 | protein_coding | 0.256108692 | 0 | 1.074270087 | 0.213643 | 3.710911 | 0.033824387 | 0.002341878 |
| ENSMUSG00000020453 | 11 | Patz1 | protein_coding | 0.255602458 | 0 | 0.434276189 | -0.598493 | -1.264405 | 0.025899046 | 0.001598111 |
| ENSMUSG00000024575 | 18 | Pde6a | protein_coding | 0.255177957 | 0 | -0.022423556 | 0.186243 | 5.466846 | 1.28663E-05 | 1.08E-07 |
| ENSMUSG00000028572 | 4 | Hook1 | protein_coding | 0.246536486 | 0 | 1.252633645 | 0.812517 | 4.179874 | 0.025120228 | 0.001525643 |
| ENSMUSG00000029403 | 5 | Cdkl2 | protein_coding | 0.24115517 | 0 | -1.304459229 | -0.62573 | 2.744449 | 0.03328903 | 0.002280551 |
| ENSMUSG00000023236 | 2 | Scg5 | protein_coding | 0.234127933 | 0 | 1.284479996 | 0.632164 | 5.112316 | 0.002753513 | 8.51763E-05 |
| ENSMUSG00000027883 | 3 | Gpsm2 | protein_coding | 0.234127933 | 0 | 0.764803035 | 2.648833 | 5.269547 | 0.001411398 | 3.79451E-05 |
| ENSMUSG00000041578 | 7 | Crx | protein_coding | 0.234127933 | 0 | 0.448874683 | 0.578671 | 6.832023 | 2.53163E-05 | 2.73E-07 |
| ENSMUSG00000036395 | 9 | Glb1l2 | protein_coding | 0.234127933 | 0 | 0.428922771 | -0.035895 | 4.309983 | 0.003492608 | 0.000114261 |
| ENSMUSG00000020890 | 11 | Gucy2e | protein_coding | 0.234127933 | 0 | -0.336623544 | 0.342106 | 5.093107 | 0.00048284 | 1.05569E-05 |
| ENSMUSG00000031450 | 8 | Grk1 | protein_coding | 0.234127933 | 0 | -0.336623544 | -0.035895 | 6.854205 | 9.08001E-06 | 6.47E-08 |
| ENSMUSG00000056043 | 7 | Rgs9bp | protein_coding | 0.234127933 | 0 | -0.336623544 | -0.035895 | 6.077132 | 1.65141E-06 | 5.75E-09 |
| ENSMUSG00000021255 | 12 | Esrrb | protein_coding | 0.234127933 | 0 | -0.336623544 | -0.035895 | 5.715155 | 1.41801E-05 | 1.22E-07 |
| ENSMUSG00000032725 | 7 | Folr2 | protein_coding | 0.233275367 | 0 | 0.137631678 | 1.914351 | -2.694243 | 0.000907569 | 2.23952E-05 |
| ENSMUSG00000016494 | 1 | Cd34 | protein_coding | 0.233080064 | 0 | 0.133968749 | 0.071852 | 1.037873 | 0.000105419 | 1.62197E-06 |
| ENSMUSG00000059412 | 9 | Fxyd2 | protein_coding | 0.227619951 | 0 | -0.323280314 | 2.644091 | 2.904575 | 0.020342238 | 0.001171215 |
| ENSMUSG00000052837 | 8 | Junb | protein_coding | 0.225824849 | 0 | -0.108248532 | -0.365831 | 0.85744 | 0.000727191 | 1.73716E-05 |
| ENSMUSG00000037161 | 3 | Mgarp | protein_coding | 0.220601535 | 0 | -0.075169953 | -2.008582 | 5.698902 | 6.81309E-05 | 9.65E-07 |
| ENSMUSG00000042757 | 9 | Tmem108 | protein_coding | 0.215968291 | 0 | -1.848755819 | -1.548027 | 4.210213 | 4.97577E-05 | 6.69E-07 |
| ENSMUSG00000034248 | 14 | Slc25a37 | protein_coding | 0.202121537 | 0 | 0.505155948 | 0.205811 | -0.530236 | 0.00885202 | 0.000378481 |
| ENSMUSG00000032265 | 9 | Fam46a | protein_coding | 0.195691125 | 0 | -0.404378794 | -0.569112 | -1.21054 | 2.13098E-05 | 2.14E-07 |
| ENSMUSG00000027273 | 2 | Snap25 | protein_coding | 0.195107054 | 0 | -0.895959565 | -0.17134 | 2.091605 | 0.000895779 | 2.19792E-05 |

| | | | | | | | | | | |
|---|---|---|---|---|---|---|---|---|---|---|
| ENSMUSG00000043384 | X | Gprasp1 | protein_coding | 0.185438342 | 0 | -0.04482721 | -0.131992 | -0.87209 | 0.04379575 | 0.003408158 |
| ENSMUSG00000031216 | X | Stard8 | protein_coding | 0.183549465 | 0 | 0.339030308 | 0.755272 | -0.649657 | 0.000385208 | 8.01672E-06 |
| ENSMUSG00000016206 | 17 | H2-M3 | protein_coding | 0.179648385 | 0 | -0.271894115 | 0.045861 | 0.819357 | 0.010219925 | 0.000462624 |
| ENSMUSG00000036617 | 2 | Etl4 | protein_coding | 0.173013189 | 0 | -0.661933879 | -0.025118 | 2.806668 | 0.035863091 | 0.002532401 |
| ENSMUSG00000029804 | 6 | Herc3 | protein_coding | 0.170317188 | 0 | -0.2390926 | -1.192319 | 2.301924 | 0.013177436 | 0.000659459 |
| ENSMUSG00000035042 | 11 | Ccl5 | protein_coding | 0.167008878 | 0 | 0.030109871 | 1.415699 | 4.942751 | 2.17E-07 | 4.39E-10 |
| ENSMUSG00000006800 | 2 | Sulf2 | protein_coding | 0.160662748 | 0 | 0.069060952 | 1.057873 | -1.24406 | 0.038984354 | 0.002882235 |
| ENSMUSG00000049939 | 6 | Lrrc4 | protein_coding | 0.158566203 | 0 | 1.620354559 | 1.139702 | -3.156384 | 0.007659656 | 0.000314475 |
| ENSMUSG00000031431 | X | Tsc22d3 | protein_coding | 0.155586266 | 0 | -0.580080354 | -0.838 | 0.173534 | 6.65016E-05 | 9.26E-07 |
| ENSMUSG00000001521 | 6 | Tulp3 | protein_coding | 0.154117479 | 0 | -0.038259061 | -0.536808 | 0.682685 | 0.003731065 | 0.00012448 |
| ENSMUSG00000037936 | 5 | Scarb1 | protein_coding | 0.140533447 | 0 | 0.013377229 | 0.571991 | 1.279092 | 0.00546082 | 0.000204742 |
| ENSMUSG00000020435 | 11 | Osbp2 | protein_coding | 0.133928187 | 0 | -1.069167278 | -1.053129 | 3.854264 | 0.002751577 | 8.48936E-05 |
| ENSMUSG00000033880 | 11 | Lgals3bp | protein_coding | 0.128810758 | 0 | -0.641760447 | 0.41642 | 0.836254 | 0.00016896 | 2.88693E-06 |
| ENSMUSG00000039470 | 8 | Zdhhc2 | protein_coding | 0.108363468 | 0 | -1.217707305 | -0.005514 | 2.747619 | 0.029877359 | 0.001954888 |
| ENSMUSG00000006007 | 1 | Pdc | protein_coding | 0.100910842 | 0 | 0.157472865 | -0.430816 | 9.154045 | 0.00058637 | 1.34378E-05 |
| ENSMUSG00000025386 | 11 | Pde6g | protein_coding | 0.099456521 | 0 | 0.536790328 | -1.213751 | 8.733364 | 0.001264893 | 3.35618E-05 |
| ENSMUSG00000042476 | 5 | Abcb4 | protein_coding | 0.092981835 | 0 | 0.440324404 | -0.048744 | -0.856438 | 0.015430549 | 0.000819697 |
| ENSMUSG00000030789 | 7 | Itgax | protein_coding | 0.089890996 | 0 | -0.085712028 | 1.562794 | 1.307967 | 0.046372556 | 0.003680064 |
| ENSMUSG00000060802 | 2 | B2m | protein_coding | 0.08324408 | 0 | -0.203467387 | 0.574726 | 1.028936 | 1.87874E-05 | 1.78E-07 |
| ENSMUSG00000028514 | 4 | Usp24 | protein_coding | 0.082899007 | 0 | 0.104426679 | -0.118944 | -1.079194 | 0.002941425 | 9.24255E-05 |
| ENSMUSG00000032262 | 9 | Elovl4 | protein_coding | 0.068398544 | 0 | -0.36499287 | -0.192959 | 6.958284 | 4.19007E-05 | 5.29E-07 |
| ENSMUSG00000032064 | 9 | Dixdc1 | protein_coding | 0.066894288 | 0 | -0.548341864 | 0.933567 | 3.990514 | 0.020156626 | 0.001157264 |
| ENSMUSG00000049130 | 7 | C5ar1 | protein_coding | 0.062302046 | 0 | -0.182550591 | -0.152001 | 0.932074 | 0.000575584 | 1.30401E-05 |
| ENSMUSG00000043467 | 1 | Zbtb37 | protein_coding | 0.060819728 | 0 | 0.403214529 | 0.24202 | -1.06042 | 0.027938448 | 0.00179635 |
| ENSMUSG00000045658 | 1 | Pid1 | protein_coding | 0.052855068 | 0 | -0.15706789 | -0.145492 | -1.078899 | 2.20586E-05 | 2.30E-07 |
| ENSMUSG00000054256 | 5 | Msi1 | protein_coding | 0.044538182 | 0 | -0.435701099 | -0.186082 | 3.806689 | 0.012254346 | 0.000598368 |
| ENSMUSG00000018428 | 11 | Akap1 | protein_coding | 0.040104953 | 0 | -2.268644736 | -0.616881 | 0.537586 | 0.045836113 | 0.003607798 |
| ENSMUSG00000018927 | 11 | Ccl6 | protein_coding | 0.038924659 | 0 | 0.449401429 | 0.64 | 1.540994 | 1.91225E-06 | 7.28E-09 |
| ENSMUSG00000036334 | 3 | Igsf10 | protein_coding | 0.038200715 | 0 | 1.807919327 | 1.429005 | 0.541763 | 0.025107579 | 0.001522842 |
| ENSMUSG00000044365 | 3 | Cxxc4 | protein_coding | 0.025729975 | 0 | -1.185632793 | 0.074862 | 2.282304 | 0.025274875 | 0.001547316 |
| ENSMUSG00000064023 | 7 | Klk8 | protein_coding | 0.023320182 | 0 | 0.083986018 | 0.38099 | 1.043305 | 0.010946847 | 0.000503183 |
| ENSMUSG00000004837 | 11 | Grap | protein_coding | 0.021001139 | 0 | 0.916008018 | -0.511934 | 0.119749 | 0.01271895 | 0.000627244 |
| ENSMUSG00000074505 | 9 | Fat3 | protein_coding | 0.019692778 | 0 | 0.324588406 | -0.371112 | -1.393823 | 0.012809442 | 0.000636893 |
| ENSMUSG00000020212 | 10 | Mdm1 | protein_coding | 0.018987023 | 0 | -0.172293055 | 0.183557 | 1.764 | 0.033650566 | 0.002315827 |
| ENSMUSG00000032609 | 9 | Klhdc8b | protein_coding | 0.016643174 | 0 | 0.861708151 | -0.100484 | -0.327124 | 0.0002754 | 5.34395E-06 |
| ENSMUSG00000021928 | 14 | Ebpl | protein_coding | 0.014630119 | 0 | -0.039813804 | -0.543902 | 0.97859 | 0.013949393 | 0.000719509 |
| ENSMUSG00000038473 | 1 | Nos1ap | protein_coding | -0.010049864 | 0 | 0.462739615 | -0.130872 | -0.930112 | 0.00780816 | 0.000322481 |
| ENSMUSG00000091537 | 9 | Tma7 | protein_coding | -0.013981513 | 0 | 0.12558575 | 0.163073 | 1.621876 | 0.006403478 | 0.000250974 |
| ENSMUSG00000045095 | 6 | Magi1 | protein_coding | -0.015155681 | 0 | 0.495394278 | -0.011255 | -0.702787 | 0.046696489 | 0.003723448 |
| ENSMUSG00000020697 | 11 | Lig3 | protein_coding | -0.015347985 | 0 | 1.074994799 | 0.5271 | -0.329469 | 0.025195821 | 0.001532275 |
| ENSMUSG00000020526 | 11 | Znhit3 | protein_coding | -0.017102767 | 0 | 0.504914583 | -0.994472 | 1.315759 | 0.012254346 | 0.000597093 |
| ENSMUSG00000042015 | 13 | Wdr41 | protein_coding | -0.024897575 | 0 | 0.262793625 | 0.675006 | -0.444642 | 0.014206841 | 0.000735134 |
| ENSMUSG00000035692 | 4 | Isg15 | protein_coding | -0.02504931 | 0 | -0.980244131 | 1.396788 | 2.194882 | 0.000570134 | 1.28348E-05 |
| ENSMUSG00000061080 | 16 | Lsamp | protein_coding | -0.034069734 | 0 | 0.121659585 | -0.449196 | 3.397481 | 0.032936792 | 0.002245751 |
| ENSMUSG00000022311 | 15 | Csmd3 | protein_coding | -0.050023933 | 0 | 0.49465163 | -0.33246 | -2.230185 | 9.51E-08 | 1.77E-10 |
| ENSMUSG00000039899 | 5 | Fgl2 | protein_coding | -0.054346679 | 0 | -0.721130258 | 1.92159 | 2.508752 | 0.000335255 | 6.78708E-06 |
| ENSMUSG00000023044 | 15 | Csad | protein_coding | -0.063916533 | 0 | -0.052968414 | -0.044709 | -1.252321 | 4.36631E-05 | 5.59E-07 |
| ENSMUSG00000030342 | 6 | Cd9 | protein_coding | -0.065588009 | 0 | -0.126196089 | -0.06827 | 1.049212 | 2.13098E-05 | 2.13E-07 |
| ENSMUSG00000028691 | 4 | Prdx1 | protein_coding | -0.0740433 | 0 | -0.129899574 | 0.375362 | 0.995949 | 0.004192068 | 0.000146649 |
| ENSMUSG00000013663 | 19 | Pten | protein_coding | -0.074828381 | 0 | -0.126461626 | -0.275561 | -1.115479 | 0.004221514 | 0.000148021 |
| ENSMUSG00000026712 | 2 | Mrc1 | protein_coding | -0.075354294 | 0 | -0.524356777 | 0.571175 | -2.612383 | 0.000500119 | 1.09752E-05 |
| ENSMUSG00000054619 | 15 | Mettl7a1 | protein_coding | -0.08340858 | 0 | 0.57820946 | 0.123298 | -0.951134 | 0.01966127 | 0.001113766 |
| ENSMUSG00000032517 | 9 | Mobp | protein_coding | -0.085967058 | 0 | 1.043788746 | 0.320741 | -5.264441 | 3.78093E-05 | 4.72E-07 |
| ENSMUSG00000051586 | 6 | Mical3 | protein_coding | -0.091144867 | 0 | 0.507362248 | 0.277144 | -0.870341 | 0.016845537 | 0.000918054 |
| ENSMUSG00000024074 | 17 | Crim1 | protein_coding | -0.097018372 | 0 | 0.679521917 | 0.117026 | -0.97512 | 0.013275273 | 0.000665947 |
| ENSMUSG00000025900 | 1 | Rp1 | protein_coding | -0.09765323 | 0 | 0.246589418 | -0.430816 | 8.003096 | 2.14258E-06 | 8.91E-09 |
| ENSMUSG00000029086 | 5 | Prom1 | protein_coding | -0.09765323 | 0 | -0.334684859 | 0.457442 | 6.387672 | 1.51367E-05 | 1.34E-07 |
| ENSMUSG00000040253 | 3 | Gbp7 | protein_coding | -0.097860128 | 0 | -0.153205672 | 0.065278 | -1.166214 | 0.000296682 | 5.86205E-06 |
| ENSMUSG00000029070 | 4 | Mxra8 | protein_coding | -0.101243797 | 0 | 0.208355044 | 0.394797 | 3.580575 | 0.022436043 | 0.001320836 |
| ENSMUSG00000086322 | 4 | E130218I03Rik | protein_coding | -0.101243797 | 0 | -0.735135169 | -0.434406 | 5.879223 | 1.7766E-05 | 1.63E-07 |
| ENSMUSG00000045763 | 15 | Basp1 | protein_coding | -0.119003101 | 0 | 0.298206191 | -0.149791 | -0.811592 | 0.001630055 | 4.51436E-05 |
| ENSMUSG00000032192 | 9 | Gnb5 | protein_coding | -0.11939383 | 0 | -0.060711292 | 0.490335 | 3.109045 | 0.035876795 | 0.002536274 |
| ENSMUSG00000024579 | 18 | Pcyox1l | protein_coding | -0.120016904 | 0 | 0.510731009 | 0.013178 | -1.631948 | 0.026019459 | 0.001613969 |
| ENSMUSG00000039396 | 8 | Neil3 | protein_coding | -0.126991789 | 0 | 3.436447119 | 2.974849 | 4.037354 | 0.014768822 | 0.000774977 |
| ENSMUSG00000042282 | X | Gucy2f | protein_coding | -0.126991789 | 0 | 1.848485827 | 1.569195 | 5.262243 | 0.000375458 | 7.78341E-06 |
| ENSMUSG00000054752 | 4 | Fsd1l | protein_coding | -0.126991789 | 0 | 0.474321239 | 0.620909 | 4.421854 | 0.012254346 | 0.000589295 |
| ENSMUSG00000035403 | 2 | Crb2 | protein_coding | -0.126991789 | 0 | 0.473500905 | 0.512767 | 3.597556 | 0.019001397 | 0.001067857 |
| ENSMUSG00000039375 | 8 | Wdr17 | protein_coding | -0.126991789 | 0 | 0.380169782 | -0.035895 | 4.931953 | 0.000261475 | 4.97585E-06 |
| ENSMUSG00000040554 | 11 | Aipl1 | protein_coding | -0.126991789 | 0 | 0.340009217 | -0.035895 | 6.662079 | 5.39822E-05 | 7.34E-07 |
| ENSMUSG00000021803 | 14 | Cdhr1 | protein_coding | -0.126991789 | 0 | 0.251714957 | 1.548179 | 7.092857 | 2.29653E-05 | 2.45E-07 |
| ENSMUSG00000037446 | 17 | Tulp1 | protein_coding | -0.126991789 | 0 | 0.251714957 | 0.342106 | 8.706932 | 2.14044E-06 | 8.67E-09 |
| ENSMUSG00000039194 | 7 | Rlbp1 | protein_coding | -0.126991789 | 0 | 0.0602362 | 0.818492 | 5.795902 | 7.76915E-06 | 5.12E-08 |
| ENSMUSG00000049122 | 4 | Frmd3 | protein_coding | -0.126991789 | 0 | 0.0602362 | 0.818492 | 3.249982 | 0.025403531 | 0.001557249 |
| ENSMUSG00000029491 | 5 | Pde6b | protein_coding | -0.126991789 | 0 | 0.0602362 | -0.035895 | 8.823633 | 2.3897E-06 | 1.10E-08 |

| | | | | | | | | | | |
|---|---|---|---|---|---|---|---|---|---|---|
| ENSMUSG00000045776 | 14 | Lrtm1 | protein_coding | -0.126991789 | 0 | 0.0602362 | -0.035895 | 4.506154 | 0.00032277 | 6.48206E-06 |
| ENSMUSG00000074006 | 7 | Omp | protein_coding | -0.126991789 | 0 | -0.336623544 | 4.609623 | 0.139844 | 3.22606E-05 | 3.84E-07 |
| ENSMUSG00000027744 | 3 | Stoml3 | protein_coding | -0.126991789 | 0 | -0.336623544 | 3.471173 | 0.964244 | 0.031021216 | 0.002058193 |
| ENSMUSG00000048617 | 8 | Rtbdn | protein_coding | -0.126991789 | 0 | -0.336623544 | 1.503063 | 7.484689 | 2.13098E-05 | 2.08E-07 |
| ENSMUSG00000048988 | 5 | Elfn1 | protein_coding | -0.126991789 | 0 | -0.336623544 | 1.165191 | 3.682214 | 0.0228063 | 0.001361102 |
| ENSMUSG00000022126 | 14 | Acod1 | protein_coding | -0.126991789 | 0 | -0.336623544 | 0.900869 | 3.436773 | 0.022485444 | 0.001327386 |
| ENSMUSG00000005696 | X | Sh2d1a | protein_coding | -0.126991789 | 0 | -0.336623544 | 0.818492 | 3.902507 | 0.006398041 | 0.000250243 |
| ENSMUSG00000041534 | 14 | Rbp3 | protein_coding | -0.126991789 | 0 | -0.336623544 | 0.37436 | 7.317412 | 8.88E-08 | 1.51E-10 |
| ENSMUSG00000021396 | 13 | Nxnl2 | protein_coding | -0.126991789 | 0 | -0.336623544 | 0.342106 | 6.452934 | 1.36705E-05 | 1.16E-07 |
| ENSMUSG00000047034 | 15 | Ankrd33 | protein_coding | -0.126991789 | 0 | -0.336623544 | 0.342106 | 5.7599 | 2.13098E-05 | 2.07E-07 |
| ENSMUSG00000058831 | 6 | Opn1sw | protein_coding | -0.126991789 | 0 | -0.336623544 | 0.342106 | 4.359058 | 0.000597757 | 1.37955E-05 |
| ENSMUSG00000075410 | 11 | Prcd | protein_coding | -0.126991789 | 0 | -0.336623544 | -0.035895 | 8.217105 | 8.65066E-05 | 1.26093E-06 |
| ENSMUSG00000031293 | X | Rs1 | protein_coding | -0.126991789 | 0 | -0.336623544 | -0.035895 | 8.17708 | 7.99961E-06 | 5.38E-08 |
| ENSMUSG00000034829 | 8 | Nxnl1 | protein_coding | -0.126991789 | 0 | -0.336623544 | -0.035895 | 7.502637 | 9.72945E-06 | 7.31E-08 |
| ENSMUSG00000032292 | 9 | Nr2e3 | protein_coding | -0.126991789 | 0 | -0.336623544 | -0.035895 | 7.111723 | 1.26838E-05 | 1.05E-07 |
| ENSMUSG00000075330 | 16 | A930003A15Rik | protein_coding | -0.126991789 | 0 | -0.336623544 | -0.035895 | 6.920872 | 3.04938E-05 | 3.56E-07 |
| ENSMUSG00000025496 | 7 | Drd4 | protein_coding | -0.126991789 | 0 | -0.336623544 | -0.035895 | 6.178673 | 1.84664E-05 | 1.73E-07 |
| ENSMUSG00000063681 | 1 | Crb1 | protein_coding | -0.126991789 | 0 | -0.336623544 | -0.035895 | 6.135309 | 7.25816E-06 | 4.64E-08 |
| ENSMUSG00000041460 | 6 | Cacna2d4 | protein_coding | -0.126991789 | 0 | -0.336623544 | -0.035895 | 5.899712 | 2.01873E-05 | 1.93E-07 |
| ENSMUSG00000043418 | 14 | Lrit2 | protein_coding | -0.126991789 | 0 | -0.336623544 | -0.035895 | 5.790562 | 8.4183E-06 | 5.89E-08 |
| ENSMUSG00000046049 | 14 | Rp1l1 | protein_coding | -0.126991789 | 0 | -0.336623544 | -0.035895 | 5.677697 | 4.73789E-06 | 2.76E-08 |
| ENSMUSG00000047298 | 19 | Kcnv2 | protein_coding | -0.126991789 | 0 | -0.336623544 | -0.035895 | 5.622479 | 1.04152E-05 | 8.10E-08 |
| ENSMUSG00000021363 | 13 | Mak | protein_coding | -0.126991789 | 0 | -0.336623544 | -0.035895 | 5.414052 | 2.64894E-05 | 2.98E-07 |
| ENSMUSG00000039714 | 9 | Cplx3 | protein_coding | -0.126991789 | 0 | -0.336623544 | -0.035895 | 5.276222 | 8.4183E-06 | 5.93E-08 |
| ENSMUSG00000024842 | 19 | Cabp4 | protein_coding | -0.126991789 | 0 | -0.336623544 | -0.035895 | 5.126927 | 4.91175E-06 | 2.98E-08 |
| ENSMUSG00000042961 | 15 | Egflam | protein_coding | -0.126991789 | 0 | -0.336623544 | -0.035895 | 5.034055 | 2.53726E-05 | 2.77E-07 |
| ENSMUSG00000020599 | 11 | Rgs9 | protein_coding | -0.126991789 | 0 | -0.336623544 | -0.035895 | 4.810703 | 0.000149999 | 2.50222E-06 |
| ENSMUSG00000030523 | 7 | Trpm1 | protein_coding | -0.126991789 | 0 | -0.336623544 | -0.035895 | 4.627695 | 8.79236E-05 | 1.2887E-06 |
| ENSMUSG00000041044 | 14 | Lrit1 | protein_coding | -0.126991789 | 0 | -0.336623544 | -0.035895 | 4.53097 | 0.000298452 | 5.94535E-06 |
| ENSMUSG00000038115 | 6 | Ano2 | protein_coding | -0.126991789 | 0 | -0.336623544 | -0.035895 | 4.415485 | 7.27271E-05 | 1.04241E-06 |
| ENSMUSG00000032343 | 9 | Impg1 | protein_coding | -0.126991789 | 0 | -0.336623544 | -0.035895 | 4.178007 | 0.001470572 | 3.97035E-05 |
| ENSMUSG00000026989 | 2 | Dapl1 | protein_coding | -0.126991789 | 0 | -0.336623544 | -0.035895 | 4.162673 | 0.000210987 | 3.79295E-06 |
| ENSMUSG00000106379 | 5 | Lhfpl3 | protein_coding | -0.126991789 | 0 | -0.336623544 | -0.035895 | 3.997742 | 0.000689641 | 1.6307E-05 |
| ENSMUSG00000041193 | 4 | Pla2g5 | protein_coding | -0.126991789 | 0 | -0.336623544 | -0.035895 | 3.837718 | 0.003277453 | 0.000105896 |
| ENSMUSG00000051860 | 3 | Samd7 | protein_coding | -0.126991789 | 0 | -0.336623544 | -0.035895 | 3.795792 | 0.007528323 | 0.000304715 |
| ENSMUSG00000030201 | 6 | Lrp6 | protein_coding | -0.136731463 | 0 | -0.393748258 | -0.087849 | -1.328514 | 0.040171417 | 0.00301554 |
| ENSMUSG00000043940 | 5 | Wdfy3 | protein_coding | -0.13876089 | 0 | -0.040938006 | 0.234696 | -0.780141 | 0.01526606 | 0.000804778 |
| ENSMUSG00000039137 | 4 | Whrn | protein_coding | -0.139434395 | 0 | 0.069361636 | -0.288658 | -0.987533 | 4.5444E-05 | 5.92E-07 |
| ENSMUSG00000037706 | 7 | Cd81 | protein_coding | -0.144840008 | 0 | -0.132772424 | -0.175774 | 0.878565 | 9.56315E-05 | 1.42491E-06 |
| ENSMUSG00000055435 | 8 | Maf | protein_coding | -0.14582738 | 0 | 0.062024007 | 0.172535 | -0.913367 | 8.4183E-06 | 5.81E-08 |
| ENSMUSG00000001120 | 10 | Pcbp3 | protein_coding | -0.150092248 | 0 | 0.087224288 | 1.202701 | 3.687321 | 0.027541599 | 0.001750762 |
| ENSMUSG00000042429 | 1 | Adora1 | protein_coding | -0.151941938 | 0 | 0.256231611 | 0.83927 | 3.40565 | 0.026079353 | 0.001619796 |
| ENSMUSG00000027562 | 3 | Car2 | protein_coding | -0.154823103 | 0 | 0.198764216 | 0.619946 | 3.43667 | 0.002766677 | 8.58713E-05 |
| ENSMUSG00000024597 | 18 | Slc12a2 | protein_coding | -0.157180509 | 0 | -0.294506587 | 0.272393 | -1.083071 | 0.000230279 | 4.28895E-06 |
| ENSMUSG00000020300 | 11 | Cpeb4 | protein_coding | -0.165251435 | 0 | 0.154009972 | 0.253377 | -0.785179 | 0.04208381 | 0.003242379 |
| ENSMUSG00000056602 | 5 | Fry | protein_coding | -0.168585399 | 0 | 0.098366417 | -0.348039 | -1.237703 | 0.000187471 | 3.28591E-06 |
| ENSMUSG00000038543 | 3 | BC028528 | protein_coding | -0.175585808 | 0 | -0.24073347 | -0.196185 | 0.86655 | 0.005086256 | 0.000186168 |
| ENSMUSG00000030324 | 6 | Rho | protein_coding | -0.178404914 | 0 | -1.247642425 | 1.23281 | 10.83007 | 0.000243895 | 4.62154E-06 |
| ENSMUSG00000039115 | 9 | Itga9 | protein_coding | -0.183911758 | 0 | 0.096385878 | 0.074736 | -0.923192 | 0.003999509 | 0.000134731 |
| ENSMUSG00000079164 | 1 | Tlr5 | protein_coding | -0.204779762 | 0 | 0.155867205 | -0.990227 | -3.382831 | 0.011464118 | 0.000538307 |
| ENSMUSG00000000244 | 7 | Tspan32 | protein_coding | -0.208327982 | 0 | -0.315328226 | 0.288123 | 2.043549 | 2.90E-07 | 6.57E-10 |
| ENSMUSG00000023940 | 17 | Sgo1 | protein_coding | -0.208356764 | 0 | 0.697487502 | 2.938432 | 3.53746 | 0.036489027 | 0.002603193 |
| ENSMUSG00000000594 | 11 | Gm2a | protein_coding | -0.222222451 | 0 | 0.545382295 | 0.613194 | 1.13054 | 0.001065043 | 2.75985E-05 |
| ENSMUSG00000024975 | 19 | Pdcd4 | protein_coding | -0.224090913 | 0 | 0.528872616 | 0.204602 | -0.586976 | 0.012105977 | 0.00057854 |
| ENSMUSG00000031210 | X | Gpr165 | protein_coding | -0.228036981 | 0 | 0.594050909 | -0.316745 | -4.149159 | 2.62229E-05 | 2.89E-07 |
| ENSMUSG00000030691 | 7 | Fchsd2 | protein_coding | -0.233933327 | 0 | 0.109219293 | -0.090682 | -1.35233 | 1.45762E-05 | 1.26E-07 |
| ENSMUSG00000042286 | 14 | Stab1 | protein_coding | -0.24430581 | 0 | 0.008143334 | -0.065516 | -1.514465 | 5.33716E-06 | 3.28E-08 |
| ENSMUSG00000032679 | 2 | Cd59a | protein_coding | -0.245947561 | 0 | 1.057156061 | 0.404428 | 3.526814 | 0.014627513 | 0.000762824 |
| ENSMUSG00000028565 | 4 | Nfia | protein_coding | -0.247552546 | 0 | 0.548883437 | -0.101867 | -0.532339 | 0.002561191 | 7.81901E-05 |
| ENSMUSG00000002741 | 11 | Ykt6 | protein_coding | -0.270950347 | 0 | 0.52350826 | -0.32792 | -0.663944 | 0.004290674 | 0.00015174 |
| ENSMUSG00000045216 | 1 | Hs6st1 | protein_coding | -0.275669921 | 0 | 0.056977207 | -0.222535 | -1.043047 | 0.035853546 | 0.002528823 |
| ENSMUSG00000021221 | 12 | Dpf3 | protein_coding | -0.277700555 | 0 | -0.100647145 | -0.610863 | 3.906054 | 0.00224933 | 6.55728E-05 |
| ENSMUSG00000021846 | 14 | Peli2 | protein_coding | -0.278491212 | 0 | -0.579846064 | -0.354691 | -1.177019 | 0.02128011 | 0.001242446 |
| ENSMUSG00000025451 | 13 | Paip1 | protein_coding | -0.28690937 | 0 | 0.978034053 | 0.557741 | -0.168872 | 0.002198072 | 6.37225E-05 |
| ENSMUSG00000003206 | 17 | Ebi3 | protein_coding | -0.287936006 | 0 | -0.783571238 | -0.145767 | 0.420567 | 0.028425344 | 0.001839165 |
| ENSMUSG00000052727 | 13 | Map1b | protein_coding | -0.288697991 | 0 | -0.159133743 | -0.056282 | 2.324607 | 0.000105524 | 1.63213E-06 |
| ENSMUSG00000022706 | 16 | Mrpl40 | protein_coding | -0.288828567 | 0 | -0.293388804 | -0.922292 | 0.528164 | 0.034809016 | 0.002424231 |
| ENSMUSG00000032294 | 9 | Pkm | protein_coding | -0.292542855 | 0 | -0.091168031 | -0.083767 | 1.005809 | 0.001055247 | 2.69319E-05 |
| ENSMUSG00000024529 | 18 | Lox | protein_coding | -0.297010564 | 0 | 0.369671341 | 3.588737 | 2.658184 | 0.01966127 | 0.001114494 |
| ENSMUSG00000032245 | 9 | Cln6 | protein_coding | -0.297890626 | 0 | -2.115566945 | 0.621759 | 0.2619 | 0.009121838 | 0.000399457 |
| ENSMUSG00000021241 | 12 | Isca2 | protein_coding | -0.314543019 | 0 | -0.157674752 | 0.064179 | 0.696748 | 0.047355196 | 0.003811731 |
| ENSMUSG00000020077 | 10 | Srgn | protein_coding | -0.321064633 | 0 | -0.161382087 | -0.100511 | 0.829235 | 0.001050026 | 2.66992E-05 |
| ENSMUSG00000029632 | 6 | Ndufa4 | protein_coding | -0.322212368 | 0 | -0.226606039 | -0.034114 | 0.783067 | 0.02673235 | 0.001671178 |
| ENSMUSG00000042363 | 11 | Lgalsl | protein_coding | -0.325078135 | 0 | 0.181094988 | 0.841911 | 2.662642 | 0.040768044 | 0.003086737 |

| | | | | | | | | | | |
|---|---|---|---|---|---|---|---|---|---|---|
| ENSMUSG00000005233 | 2 | Spc25 | protein_coding | -0.329815294 | 0 | -1.741249258 | 0.487946 | 2.571267 | 0.009340965 | 0.000413002 |
| ENSMUSG00000005469 | 8 | Prkaca | protein_coding | -0.331133654 | 0 | 0.925866124 | 0.519092 | 0.238201 | 0.036489027 | 0.002601173 |
| ENSMUSG00000019874 | 10 | Fabp7 | protein_coding | -0.340840451 | 0 | 1.156077467 | 5.748325 | -0.054598 | 2.25586E-05 | 2.39E-07 |
| ENSMUSG00000015243 | 4 | Abca1 | protein_coding | -0.348148565 | 0 | -0.084122307 | 0.295101 | -2.143921 | 1.02E-09 | 4.94E-13 |
| ENSMUSG00000003500 | 6 | Impdh1 | protein_coding | -0.358988652 | 0 | -1.560840912 | -1.456801 | 4.494811 | 0.000182325 | 3.15957E-06 |
| ENSMUSG00000031604 | 8 | Msmo1 | protein_coding | -0.361021977 | 0 | -0.296744635 | -0.162787 | 1.042027 | 0.00257021 | 7.86735E-05 |
| ENSMUSG00000063229 | 7 | Ldha | protein_coding | -0.362439878 | 0 | -0.304266047 | -0.132022 | 0.715456 | 0.015250558 | 0.000802726 |
| ENSMUSG00000026473 | 1 | Glul | protein_coding | -0.373434419 | 0 | 0.206668013 | -0.085053 | -0.837677 | 2.94076E-05 | 3.41E-07 |
| ENSMUSG00000005360 | 15 | Slc1a3 | protein_coding | -0.379847195 | 0 | -0.074170569 | 0.067001 | -1.482362 | 5.47E-08 | 7.97E-11 |
| ENSMUSG00000076431 | 13 | Sox4 | protein_coding | -0.382624879 | 0 | 0.309135504 | 0.12139 | -1.449149 | 6.85E-07 | 1.89E-09 |
| ENSMUSG00000060206 | 4 | Zfp462 | protein_coding | -0.391712885 | 0 | -0.336551073 | 0.484102 | 4.183223 | 0.004129086 | 0.000141267 |
| ENSMUSG00000014956 | 5 | Ppp1cb | protein_coding | -0.392635549 | 0 | 0.130173641 | -0.080674 | -0.909499 | 0.027240648 | 0.001729425 |
| ENSMUSG00000044626 | 16 | Liph | protein_coding | -0.401076299 | 0 | -0.309421338 | -0.069185 | 0.654108 | 0.003686528 | 0.000122695 |
| ENSMUSG00000018378 | 11 | Cuedc1 | protein_coding | -0.402150283 | 0 | 0.294463734 | -0.575808 | 3.71396 | 0.007649596 | 0.000313313 |
| ENSMUSG00000031540 | 8 | Kat6a | protein_coding | -0.430105182 | 0 | -0.173747908 | -0.298837 | -1.13429 | 0.018578489 | 0.001035064 |
| ENSMUSG00000038502 | 7 | Ptov1 | protein_coding | -0.432860961 | 0 | -0.169234934 | -0.854491 | -1.018954 | 0.04208381 | 0.003245725 |
| ENSMUSG00000029096 | 5 | Htra3 | protein_coding | -0.43428544 | 0 | 0.42620796 | -0.019319 | 3.979532 | 0.007528323 | 0.000305145 |
| ENSMUSG00000018476 | 11 | Kdm6b | protein_coding | -0.441030231 | 0 | 0.741606812 | 0.076136 | -0.804422 | 0.00230382 | 6.80941E-05 |
| ENSMUSG00000030541 | 7 | Idh2 | protein_coding | -0.448721693 | 0 | 0.154776597 | 0.071779 | -0.994052 | 0.002523772 | 7.6639E-05 |
| ENSMUSG00000032643 | 4 | Fhl3 | protein_coding | -0.453715722 | 0 | 0.799443277 | 0.126676 | -0.999521 | 0.032289886 | 0.002184696 |
| ENSMUSG00000021719 | 13 | Rgs7bp | protein_coding | -0.45458574 | 0 | 0.036606804 | 0.222727 | -3.334483 | 1.80983E-05 | 1.67E-07 |
| ENSMUSG00000021951 | 14 | Eef1akmt1 | protein_coding | -0.455884033 | 0 | -0.253186077 | -0.179199 | 1.078844 | 0.049821775 | 0.004070789 |
| ENSMUSG00000057132 | 14 | Rpgrip1 | protein_coding | -0.457864972 | 0 | 0.428834333 | 0.095945 | 1.71768 | 0.028434853 | 0.001842083 |
| ENSMUSG00000034413 | 17 | Neurl1b | protein_coding | -0.459999701 | 0 | -0.200966432 | 3.075536 | 2.078432 | 0.034666794 | 0.002411432 |
| ENSMUSG00000012819 | 10 | Cdh23 | protein_coding | -0.465161888 | 0 | 0.259380401 | -0.76711 | -1.03701 | 0.031322382 | 0.002100165 |
| ENSMUSG00000032470 | 9 | Mras | protein_coding | -0.46612578 | 0 | -0.052283455 | -0.089417 | -1.587029 | 0.004290674 | 0.000151836 |
| ENSMUSG00000045268 | 4 | Zfp691 | protein_coding | -0.467293382 | 0 | 0.417563564 | -0.205245 | -0.71118 | 0.035385269 | 0.002475737 |
| ENSMUSG00000024924 | 19 | Vldlr | protein_coding | -0.47265169 | 0 | -2.764835041 | -1.481429 | 4.155588 | 5.59839E-05 | 7.71E-07 |
| ENSMUSG00000035778 | 2 | Ggta1 | protein_coding | -0.477805445 | 0 | -0.352987158 | -0.380762 | -1.349748 | 0.008927942 | 0.000385342 |
| ENSMUSG00000031938 | 9 | 4931406C07Rik | protein_coding | -0.479224121 | 0 | -0.303344328 | -0.140225 | -1.052646 | 0.016592864 | 0.000898909 |
| ENSMUSG00000020907 | 11 | Rcvrn | protein_coding | -0.485105807 | 0 | -1.11899718 | 0.186889 | 8.497318 | 0.001539621 | 4.22651E-05 |
| ENSMUSG00000041817 | 13 | Fam169a | protein_coding | -0.48897826 | 0 | 0.491867687 | -0.44414 | 4.920069 | 0.000168857 | 2.87149E-06 |
| ENSMUSG00000070462 | 7 | Tlnrd1 | protein_coding | -0.494192954 | 0 | -0.513638484 | 0.299129 | -2.749683 | 0.01459084 | 0.000757253 |
| ENSMUSG00000037940 | 8 | Inpp4b | protein_coding | -0.497685325 | 0 | -0.011893225 | -0.181006 | -1.280433 | 5.39916E-06 | 3.37E-08 |
| ENSMUSG00000019189 | 11 | Rnf145 | protein_coding | -0.498828353 | 0 | -0.444467151 | -0.368523 | -1.012895 | 0.022649402 | 0.00134807 |
| ENSMUSG00000020333 | 11 | Acsl6 | protein_coding | -0.501862678 | 0 | 0.877454495 | 1.089129 | 3.460256 | 0.049247044 | 0.003987938 |
| ENSMUSG00000043391 | 16 | 2510009E07Rik | protein_coding | -0.503937804 | 0 | -0.568807845 | 0.064175 | 0.78883 | 0.001055247 | 2.70883E-05 |
| ENSMUSG00000035847 | X | Ids | protein_coding | -0.504462964 | 0 | 0.129988189 | -0.147134 | -1.043862 | 0.049340891 | 0.003999533 |
| ENSMUSG00000026094 | 1 | Stk17b | protein_coding | -0.517245747 | 0 | -0.820602948 | -0.571382 | -1.551028 | 0.000847919 | 2.06676E-05 |
| ENSMUSG00000048503 | 9 | Tmem136 | protein_coding | -0.521912848 | 0 | 0.918190052 | -0.430816 | 4.210175 | 0.007328988 | 0.00029437 |
| ENSMUSG00000040632 | 14 | Nrl | protein_coding | -0.521912848 | 0 | 0.510845663 | -0.052815 | 8.118235 | 0.00020858 | 3.7159E-06 |
| ENSMUSG00000023978 | 17 | Prph2 | protein_coding | -0.521912848 | 0 | -0.338137279 | 0.117846 | 9.821305 | 6.54803E-05 | 9.07E-07 |
| ENSMUSG00000024519 | 18 | Cplx4 | protein_coding | -0.521912848 | 0 | -0.731544602 | -0.430816 | 7.185401 | 3.7748E-06 | 2.08E-08 |
| ENSMUSG00000096351 | 4 | Samd11 | protein_coding | -0.523377479 | 0 | 1.31223584 | -0.104252 | 4.425188 | 0.001090697 | 2.84399E-05 |
| ENSMUSG00000031789 | 8 | Cngb1 | protein_coding | -0.525503415 | 0 | -0.735135169 | 0.363613 | 8.775912 | 2.37421E-06 | 1.08E-08 |
| ENSMUSG00000044147 | 12 | Arf6 | protein_coding | -0.532706648 | 0 | 0.772318649 | 0.067876 | -0.032959 | 0.033981399 | 0.002358252 |
| ENSMUSG00000025151 | X | Maged1 | protein_coding | -0.533773255 | 0 | 0.250171966 | -0.221522 | -0.870756 | 0.013668512 | 0.000697317 |
| ENSMUSG00000038963 | 2 | Slco4a1 | protein_coding | -0.542290366 | 0 | -0.020371114 | -1.279835 | -0.666693 | 0.011269934 | 0.00052493 |
| ENSMUSG00000024985 | 19 | Tcf7l2 | protein_coding | -0.549087521 | 0 | 0.335634044 | -0.085263 | -0.68897 | 0.013855695 | 0.000710187 |
| ENSMUSG00000039202 | 7 | Abhd2 | protein_coding | -0.549869355 | 0 | 0.143642489 | -0.053311 | -1.082676 | 0.049530511 | 0.004030947 |
| ENSMUSG00000007670 | 17 | Khsrp | protein_coding | -0.551024885 | 0 | 0.487634423 | 0.097204 | -0.611158 | 0.003970755 | 0.000133441 |
| ENSMUSG00000026303 | 1 | Mlph | protein_coding | -0.55214855 | 0 | 0.117649457 | -0.160636 | -2.337872 | 9.72945E-06 | 7.31E-08 |
| ENSMUSG00000049866 | 1 | Arl4c | protein_coding | -0.554531475 | 0 | -0.145061464 | -0.338282 | -1.177644 | 0.003544709 | 0.00011654 |
| ENSMUSG00000029064 | 4 | Gnb1 | protein_coding | -0.559257997 | 0 | -0.16418392 | -0.13203 | 1.998733 | 4.22577E-06 | 2.43E-08 |
| ENSMUSG00000060002 | 10 | Chpt1 | protein_coding | -0.562823027 | 0 | -0.50329853 | -0.412944 | -1.282523 | 0.033738255 | 0.002324986 |
| ENSMUSG00000031714 | 8 | Gab1 | protein_coding | -0.56791156 | 0 | -0.009487074 | -0.594228 | -1.422228 | 0.005102471 | 0.000188201 |
| ENSMUSG00000001424 | 6 | Snd1 | protein_coding | -0.578587645 | 0 | 0.576316203 | 0.466994 | -0.225283 | 0.00780816 | 0.000324365 |
| ENSMUSG00000002608 | 7 | Ccdc97 | protein_coding | -0.586848775 | 0 | -0.166614273 | -0.742869 | -1.188008 | 0.037713411 | 0.002721083 |
| ENSMUSG00000028184 | 3 | Adgrl2 | protein_coding | -0.59169705 | 0 | -0.107937354 | -0.312132 | -1.57268 | 0.007712556 | 0.000317895 |
| ENSMUSG00000024030 | 17 | Abcg1 | protein_coding | -0.594156857 | 0 | 0.335370058 | 0.372576 | -2.128156 | 1.85961E-06 | 6.93E-09 |
| ENSMUSG00000037788 | 6 | Vopp1 | protein_coding | -0.596823527 | 0 | -0.154187872 | -0.314479 | 2.95708 | 0.02627586 | 0.001634129 |
| ENSMUSG00000024500 | 18 | Ppp2r2b | protein_coding | -0.612927351 | 0 | -1.350547114 | 1.014438 | 3.816699 | 0.001568694 | 4.31902E-05 |
| ENSMUSG00000020387 | 11 | Jade2 | protein_coding | -0.616613449 | 0 | -0.603175169 | -0.672657 | -1.380309 | 0.007273596 | 0.000291556 |
| ENSMUSG00000050965 | 11 | Prkca | protein_coding | -0.617612486 | 0 | -0.004922025 | -0.481979 | -1.527101 | 0.000405682 | 8.60706E-06 |
| ENSMUSG00000052310 | 3 | Slc39a1 | protein_coding | -0.623580136 | 0 | 0.89340976 | 0.448671 | -0.408804 | 0.011107372 | 0.000513589 |
| ENSMUSG00000040725 | 7 | Hnrnpul1 | protein_coding | -0.625228337 | 0 | 0.437741199 | 0.165886 | -0.757103 | 0.002405033 | 7.18647E-05 |
| ENSMUSG00000024099 | 17 | Ndufv2 | protein_coding | -0.631416136 | 0 | 0.02775447 | -0.093208 | 0.62345 | 0.012677366 | 0.00062314 |
| ENSMUSG00000030516 | 7 | Tjp1 | protein_coding | -0.631875642 | 0 | -0.248335597 | -0.507462 | -1.038779 | 0.004482365 | 0.000160071 |
| ENSMUSG00000036327 | 2 | Qsox2 | protein_coding | -0.634923639 | 0 | -1.221058822 | -2.809893 | 0.533299 | 0.033044827 | 0.002258469 |
| ENSMUSG00000025104 | 7 | Hdgfl3 | protein_coding | -0.641491298 | 0 | 0.615498252 | 1.247725 | 3.217125 | 0.030639592 | 0.002022129 |
| ENSMUSG00000023982 | 17 | Guca1a | protein_coding | -0.643215674 | 0 | -0.243843482 | -0.233999 | 2.643852 | 0.012254346 | 0.000592567 |
| ENSMUSG00000074570 | 2 | Cass4 | protein_coding | -0.651123812 | 0 | 0.35581387 | -0.974817 | -2.018458 | 0.004471991 | 0.000159339 |
| ENSMUSG00000045193 | 10 | Cirbp | protein_coding | -0.656216942 | 0 | 0.498278555 | 0.474775 | 0.960964 | 0.022516041 | 0.001331015 |
| ENSMUSG00000074227 | 7 | Spint2 | protein_coding | -0.674520353 | 0 | -1.245271829 | -0.534288 | 3.103926 | 0.012254346 | 0.000592209 |

| | | | | | | | | | | |
|---|---|---|---|---|---|---|---|---|---|---|
| ENSMUSG00000047388 | 8 | Atmin | protein_coding | -0.676561912 | 0 | -0.31387969 | -0.836914 | -1.016642 | 0.038425856 | 0.002790472 |
| ENSMUSG00000022353 | 15 | Mtss1 | protein_coding | -0.677961458 | 0 | 0.01299672 | -0.405639 | -1.379416 | 0.002766677 | 8.60316E-05 |
| ENSMUSG00000020747 | 11 | Tmem94 | protein_coding | -0.677982501 | 0 | 0.554358346 | -0.13451 | -1.000174 | 0.037743783 | 0.002729387 |
| ENSMUSG00000031709 | 8 | Tbc1d9 | protein_coding | -0.678284359 | 0 | -0.284844965 | -0.557755 | -1.132152 | 0.038758677 | 0.002844248 |
| ENSMUSG00000028073 | 3 | Pear1 | protein_coding | -0.679854934 | 0 | 0.3099205 | -1.215354 | 0.62828 | 0.016113707 | 0.000870042 |
| ENSMUSG00000019122 | 11 | Ccl9 | protein_coding | -0.681679627 | 0 | 0.375812488 | -0.138762 | 0.401325 | 0.000645493 | 1.50364E-05 |
| ENSMUSG00000009090 | 11 | Ap1b1 | protein_coding | -0.687193423 | 0 | -0.369868166 | -0.372759 | -1.265782 | 0.000575584 | 1.30973E-05 |
| ENSMUSG00000050148 | X | Ubqln2 | protein_coding | -0.693960089 | 0 | -0.537151514 | -1.226206 | -1.03745 | 0.038425856 | 0.002792809 |
| ENSMUSG00000020865 | 11 | Abcc3 | protein_coding | -0.698592685 | 0 | -0.349780553 | -0.374426 | -1.464605 | 1.84664E-05 | 1.73E-07 |
| ENSMUSG00000025380 | 11 | Fscn2 | protein_coding | -0.699775561 | 0 | -0.511805002 | -0.608678 | 2.794136 | 0.041218429 | 0.003137527 |
| ENSMUSG00000024227 | 17 | Pdzph1 | protein_coding | -0.70015461 | 0 | -0.232434066 | -0.609057 | 3.154261 | 0.04014065 | 0.003006729 |
| ENSMUSG00000042684 | 1 | Npl | protein_coding | -0.700278426 | 0 | 0.092334267 | 0.486299 | -0.375733 | 0.005287995 | 0.000197834 |
| ENSMUSG00000031285 | X | Dcx | protein_coding | -0.701960173 | 0 | -0.100647145 | 3.118594 | 0.856052 | 0.038853506 | 0.00286633 |
| ENSMUSG00000079550 | 1 | Mpp4 | protein_coding | -0.701960173 | 0 | -0.345512706 | 0.906128 | 5.776008 | 0.000110975 | 1.72542E-06 |
| ENSMUSG00000045038 | 17 | Prkce | protein_coding | -0.702192106 | 0 | 0.137257137 | -0.607372 | -1.76646 | 0.000266042 | 5.0843E-06 |
| ENSMUSG00000001036 | 11 | Epn2 | protein_coding | -0.703678187 | 0 | 0.039273829 | -0.366694 | -1.230766 | 0.007528323 | 0.000305283 |
| ENSMUSG00000001763 | 6 | Tspan33 | protein_coding | -0.709206902 | 0 | -0.592885894 | -0.093428 | -1.182412 | 0.041668411 | 0.003192025 |
| ENSMUSG00000021281 | 12 | Tnfaip2 | protein_coding | -0.710970604 | 0 | -0.479987494 | 0.466857 | 0.566114 | 0.021246883 | 0.001238785 |
| ENSMUSG00000040543 | 11 | Pitpnm3 | protein_coding | -0.721270182 | 0 | -0.343707409 | 1.032638 | 4.757141 | 0.000342452 | 6.96052E-06 |
| ENSMUSG00000054850 | X | Smim10l2a | protein_coding | -0.727323024 | 0 | -0.051950527 | 0.317431 | -1.676082 | 0.04379575 | 0.003408188 |
| ENSMUSG00000027860 | 3 | Vangl1 | protein_coding | -0.75192033 | 0 | -0.284834335 | -1.743405 | -2.70044 | 0.023468732 | 0.001410047 |
| ENSMUSG00000030695 | 7 | Aldoa | protein_coding | -0.752394199 | 0 | -0.204093078 | -0.299495 | 0.804203 | 0.00074349 | 1.78211E-05 |
| ENSMUSG00000027488 | 2 | Snta1 | protein_coding | -0.754494363 | 0 | -0.184848376 | -0.75802 | -1.482078 | 0.010097503 | 0.000455447 |
| ENSMUSG00000025742 | X | Prps2 | protein_coding | -0.754930892 | 0 | -0.023057516 | -0.078701 | -1.097001 | 0.012853167 | 0.000640108 |
| ENSMUSG00000036192 | 19 | Rorb | protein_coding | -0.755646486 | 0 | 0.17180566 | 0.394982 | 6.019378 | 0.000165722 | 2.80475E-06 |
| ENSMUSG00000022496 | 16 | Tnfrsf17 | protein_coding | -0.758787405 | 0 | 0.249369385 | -0.150357 | -0.962937 | 0.045007913 | 0.003524387 |
| ENSMUSG00000045092 | 3 | S1pr1 | protein_coding | -0.774097895 | 0 | -0.174549389 | -0.109045 | -1.068038 | 0.011269934 | 0.000526581 |
| ENSMUSG00000032058 | 9 | Ppp2r1b | protein_coding | -0.781225873 | 0 | 0.053217439 | -0.929188 | -1.224714 | 0.02710853 | 0.001716647 |
| ENSMUSG00000032640 | 7 | Chsy1 | protein_coding | -0.789538693 | 0 | 0.231137167 | -0.379503 | -0.503725 | 0.004684277 | 0.000168041 |
| ENSMUSG00000044167 | 3 | Foxo1 | protein_coding | -0.789651978 | 0 | -0.437850571 | -0.491038 | -1.226783 | 0.021364278 | 0.001251547 |
| ENSMUSG00000020444 | 11 | Guk1 | protein_coding | -0.790851485 | 0 | -0.080866043 | -0.902565 | 1.219702 | 0.004249609 | 0.000149351 |
| ENSMUSG00000046245 | 5 | Pilra | protein_coding | -0.796760434 | 0 | -0.758683727 | -1.064367 | 1.181936 | 2.9399E-05 | 3.38E-07 |
| ENSMUSG00000047146 | 10 | Tet1 | protein_coding | -0.813643835 | 0 | 0.009014295 | -0.45485 | -2.235279 | 0.002941425 | 9.24347E-05 |
| ENSMUSG00000028125 | 3 | Abca4 | protein_coding | -0.816532382 | 0 | -1.026164137 | -0.725435 | 6.118153 | 1.29581E-06 | 4.30E-09 |
| ENSMUSG00000039529 | 18 | Atp8b1 | protein_coding | -0.82203528 | 0 | 1.041517707 | 0.510155 | -3.465504 | 0.005704513 | 0.000214875 |
| ENSMUSG00000005125 | 15 | Ndrg1 | protein_coding | -0.823169616 | 0 | -0.222676922 | 0.693813 | 3.373778 | 0.040343139 | 0.003044765 |
| ENSMUSG00000079481 | X | Nhsl2 | protein_coding | -0.830000119 | 0 | -0.27355383 | -0.547322 | -2.098704 | 6.77422E-05 | 9.55E-07 |
| ENSMUSG00000033174 | 6 | Mgll | protein_coding | -0.843299275 | 0 | 0.154646141 | -0.325471 | -1.29308 | 6.95E-07 | 2.03E-09 |
| ENSMUSG00000004798 | 11 | Ulk2 | protein_coding | -0.854677694 | 0 | -0.250185931 | -0.225765 | -1.400621 | 0.000113748 | 1.79617E-06 |
| ENSMUSG00000003418 | 2 | St8sia6 | protein_coding | -0.860328719 | 0 | -1.379167941 | 0.296435 | -0.227245 | 0.03760543 | 0.002703703 |
| ENSMUSG00000074794 | 13 | Arrdc3 | protein_coding | -0.864484665 | 0 | -0.705913465 | -0.913488 | -1.086182 | 0.044994176 | 0.003519667 |
| ENSMUSG00000033857 | 11 | Engase | protein_coding | -0.867861868 | 0 | -0.08889422 | -0.474161 | -1.766804 | 0.025751125 | 0.001584813 |
| ENSMUSG00000019818 | 10 | Cd164 | protein_coding | -0.870849163 | 0 | -0.064462417 | -0.431519 | -1.170937 | 8.2502E-06 | 5.61E-08 |
| ENSMUSG00000043832 | 6 | Clec4a3 | protein_coding | -0.87387816 | 0 | -0.578303649 | -0.195782 | -1.365576 | 0.000298452 | 5.92934E-06 |
| ENSMUSG00000003882 | 15 | Il7r | protein_coding | -0.877750635 | 0 | -0.044628012 | -0.57177 | -1.316065 | 0.000178898 | 3.0857E-06 |
| ENSMUSG00000023979 | 17 | Guca1b | protein_coding | -0.881293554 | 0 | -1.584837818 | -1.584829 | 4.320185 | 0.00164036 | 4.55619E-05 |
| ENSMUSG00000025993 | 1 | Slc40a1 | protein_coding | -0.889017235 | 0 | -0.433006353 | -0.321749 | -1.289771 | 4.91175E-06 | 2.98E-08 |
| ENSMUSG00000060550 | 17 | H2-Q7 | protein_coding | -0.905032041 | 0 | 0.191246881 | 1.18118 | 2.262091 | 0.036145432 | 0.0025699 |
| ENSMUSG00000034880 | 8 | Mrpl34 | protein_coding | -0.907548842 | 0 | -0.05127797 | -0.114315 | 0.456604 | 0.04208381 | 0.003247702 |
| ENSMUSG00000007097 | 1 | Atp1a2 | protein_coding | -0.930397426 | 0 | -0.294489037 | 0.281277 | -4.727945 | 0.000241837 | 4.54339E-06 |
| ENSMUSG00000033278 | 17 | Ptprm | protein_coding | -0.935656677 | 0 | 0.142383257 | -0.35751 | -1.470743 | 0.02244438 | 0.001323144 |
| ENSMUSG00000074305 | 9 | Peak1 | protein_coding | -0.944986998 | 0 | -0.129715027 | -0.340748 | -2.159628 | 2.62477E-05 | 2.91E-07 |
| ENSMUSG00000027784 | 3 | Ppm1l | protein_coding | -0.949932577 | 0 | -0.144080739 | -0.73349 | -1.934582 | 6.37824E-06 | 4.03E-08 |
| ENSMUSG00000047867 | 6 | Gimap6 | protein_coding | -0.956899608 | 0 | 0.143237083 | -0.623879 | -1.617711 | 0.00436641 | 0.000155223 |
| ENSMUSG00000027956 | 3 | Tmem144 | protein_coding | -0.966029658 | 0 | -0.046471541 | -0.025231 | -1.654474 | 3.53979E-05 | 4.27E-07 |
| ENSMUSG00000072966 | X | Gprasp2 | protein_coding | -0.966839249 | 0 | 0.311794267 | -0.457654 | -2.151709 | 0.000834918 | 2.02831E-05 |
| ENSMUSG00000032564 | 9 | Cpne4 | protein_coding | -0.982254864 | 0 | -1.191866618 | 2.974642 | -0.715419 | 0.004894447 | 0.000176769 |
| ENSMUSG00000017344 | 11 | Vtn | protein_coding | -0.983004698 | 0 | -1.553627023 | -0.840311 | 5.253813 | 0.000121086 | 1.95126E-06 |
| ENSMUSG00000058624 | 19 | Gda | protein_coding | -0.992438799 | 0 | -1.963532387 | 2.655506 | -2.435369 | 0.000883369 | 2.16032E-05 |
| ENSMUSG00000038463 | 1 | Olfml2b | protein_coding | -1.004985605 | 0 | -0.560915754 | 0.045064 | -1.143485 | 0.03594914 | 0.002546693 |
| ENSMUSG00000028525 | 4 | Pde4b | protein_coding | -1.005775276 | 0 | -0.70289739 | -0.754746 | -1.071108 | 0.008795351 | 0.000374634 |
| ENSMUSG00000058254 | X | Tspan7 | protein_coding | -1.029154175 | 0 | -0.408056706 | -0.131214 | -0.279305 | 0.032944695 | 0.002248958 |
| ENSMUSG00000060402 | 7 | Chst8 | protein_coding | -1.033594683 | 0 | 0.369039144 | -1.023013 | -2.669503 | 0.031249486 | 0.002092485 |
| ENSMUSG00000023009 | 15 | Nckap5l | protein_coding | -1.040176064 | 0 | -0.161664404 | -0.539301 | -1.405004 | 0.001044436 | 2.64725E-05 |
| ENSMUSG00000022836 | 16 | Mylk | protein_coding | -1.044905155 | 0 | -0.856934596 | -0.339242 | 3.070479 | 0.012075963 | 0.000574999 |
| ENSMUSG00000066800 | 1 | Rnasel | protein_coding | -1.083271016 | 0 | -0.496169063 | -0.467568 | -1.585976 | 0.000240028 | 4.48996E-06 |
| ENSMUSG00000046447 | 4 | Camk2n1 | protein_coding | -1.083764486 | 0 | -0.103266884 | -0.586936 | -0.32966 | 0.012254346 | 0.000596797 |
| ENSMUSG00000030538 | 7 | Cib1 | protein_coding | -1.083968037 | 0 | -0.521219817 | -0.655063 | -1.501983 | 0.000113748 | 1.78926E-06 |
| ENSMUSG00000030539 | 7 | Sema4b | protein_coding | -1.084898186 | 0 | 0.081191612 | -0.46745 | -2.537881 | 3.76E-08 | 4.64E-11 |
| ENSMUSG00000014164 | 13 | Klhl3 | protein_coding | -1.09481593 | 0 | -0.243201396 | 0.229923 | -0.937805 | 0.033981399 | 0.002356676 |
| ENSMUSG00000048706 | 4 | Lurap1l | protein_coding | -1.096157443 | 0 | -0.628436899 | 0.059194 | 3.426182 | 0.018861906 | 0.001055894 |
| ENSMUSG00000004814 | 5 | Ccl24 | protein_coding | -1.106653885 | 0 | 0.059130297 | 0.338197 | -3.559823 | 0.001922138 | 5.46336E-05 |
| ENSMUSG00000022197 | 15 | Pdzd2 | protein_coding | -1.108599631 | 0 | 1.091151531 | 1.304357 | 4.353291 | 0.001588638 | 4.3868E-05 |
| ENSMUSG00000052613 | 10 | Pcdh15 | protein_coding | -1.115909259 | 0 | -1.325541013 | -0.614558 | 3.653473 | 0.000547447 | 1.21911E-05 |

| | | | | | | | | | | |
|---|---|---|---|---|---|---|---|---|---|---|
| ENSMUSG00000010064 | 9 | Slc38a3 | protein_coding | -1.137479702 | 0 | -1.347111457 | -1.046383 | 3.871469 | 0.000241893 | 4.56402E-06 |
| ENSMUSG00000037337 | 7 | Map4k1 | protein_coding | -1.160980299 | 0 | 0.729515385 | 0.475443 | 2.823158 | 0.013517343 | 0.000686672 |
| ENSMUSG00000078616 | 7 | Trim30c | protein_coding | -1.163669971 | 0 | -0.449933705 | 0.385286 | -1.605632 | 0.005090231 | 0.000186726 |
| ENSMUSG00000022237 | 15 | Ankrd33b | protein_coding | -1.177063877 | 0 | -0.502652731 | 0.232919 | 3.464375 | 0.042964872 | 0.003329556 |
| ENSMUSG00000039831 | 3 | Arhgap29 | protein_coding | -1.211225149 | 0 | -0.30194787 | -1.008547 | -2.179336 | 0.012711348 | 0.00062584 |
| ENSMUSG00000025203 | 19 | Scd2 | protein_coding | -1.231261108 | 0 | -0.10016678 | 0.511211 | -1.564352 | 2.35766E-06 | 1.01E-08 |
| ENSMUSG00000015852 | 3 | Fcrls | protein_coding | -1.250312007 | 0 | -0.075736141 | -0.260219 | -2.069443 | 1.54E-15 | 1.24E-19 |
| ENSMUSG00000034906 | 2 | Ncaph | protein_coding | -1.268793932 | 0 | -0.48533262 | 0.247269 | -1.752289 | 0.012105977 | 0.000579369 |
| ENSMUSG00000021806 | 14 | Nid2 | protein_coding | -1.271852856 | 0 | -0.214453957 | -0.166718 | -0.586326 | 0.031037555 | 0.002065002 |
| ENSMUSG00000032306 | 9 | Mpi | protein_coding | -1.279073002 | 0 | -0.198948471 | 0.402219 | -1.296232 | 0.000178236 | 3.05985E-06 |
| ENSMUSG00000027210 | 2 | Meis2 | protein_coding | -1.289678284 | 0 | -0.359328774 | 3.955792 | 3.231919 | 0.000217975 | 3.95547E-06 |
| ENSMUSG00000029167 | 5 | Ppargc1a | protein_coding | -1.293061984 | 0 | -1.863813461 | 0.110438 | 2.661754 | 0.025991189 | 0.001610111 |
| ENSMUSG00000032666 | 1 | 1700025G04Rik | protein_coding | -1.29768542 | 0 | -0.287455492 | 0.006734 | -2.727231 | 0.035665659 | 0.002501131 |
| ENSMUSG00000029608 | 5 | Rph3a | protein_coding | -1.30378918 | 0 | -0.103895595 | -3.080888 | 0.98605 | 0.025922276 | 0.001601644 |
| ENSMUSG00000042616 | 4 | Oscp1 | protein_coding | -1.325303333 | 0 | -0.256870479 | -1.297384 | 0.245208 | 0.002352326 | 6.97183E-05 |
| ENSMUSG00000067889 | 19 | Sptbn2 | protein_coding | -1.327191754 | 0 | -0.379454851 | -2.686316 | -3.412986 | 0.027095369 | 0.001710397 |
| ENSMUSG00000058743 | 7 | Kcnj14 | protein_coding | -1.332527156 | 0 | -1.542158911 | -1.24143 | 4.488814 | 3.78093E-05 | 4.71E-07 |
| ENSMUSG00000024168 | 17 | Tmem204 | protein_coding | -1.357597347 | 0 | -0.35097787 | -0.503719 | -2.050185 | 0.001163259 | 3.04262E-05 |
| ENSMUSG00000061808 | 18 | Ttr | protein_coding | -1.392748712 | 0 | 2.04441968 | -2.467801 | 3.092226 | 0.029368665 | 0.001914469 |
| ENSMUSG00000035270 | 16 | Impg2 | protein_coding | -1.410868754 | 0 | -3.936872958 | -3.515477 | 1.263105 | 0.001042368 | 2.63356E-05 |
| ENSMUSG00000029343 | 5 | Crybb1 | protein_coding | -1.418152089 | 0 | -0.016043833 | -0.153748 | -1.561017 | 2.24E-07 | 4.71E-10 |
| ENSMUSG00000030772 | 7 | Dkk3 | protein_coding | -1.419345656 | 0 | 0.3793054 | -1.963076 | 2.83086 | 0.008346897 | 0.000353505 |
| ENSMUSG00000059994 | 3 | Fcrl1 | protein_coding | -1.432801699 | 0 | -0.218128307 | -0.605554 | -1.43158 | 0.009121838 | 0.000399621 |
| ENSMUSG00000053025 | 7 | Sv2b | protein_coding | -1.445886519 | 0 | -1.19388256 | -1.779049 | 4.402339 | 4.46031E-05 | 5.78E-07 |
| ENSMUSG00000020108 | 10 | Ddit4 | protein_coding | -1.45508296 | 0 | -0.389806297 | -0.643652 | -2.688328 | 0.00014757 | 2.42584E-06 |
| ENSMUSG00000041633 | X | Kctd12b | protein_coding | -1.497111182 | 0 | -0.581324323 | -0.516337 | -3.564599 | 0.001983375 | 5.66277E-05 |
| ENSMUSG00000020787 | 11 | P2rx1 | protein_coding | -1.534955449 | 0 | -0.640363056 | -0.668501 | 1.349865 | 0.005102471 | 0.000188032 |
| ENSMUSG00000002076 | 17 | Hsf2bp | protein_coding | -1.577200648 | 0 | 0.413034888 | -0.073884 | -3.609115 | 0.00106366 | 2.74765E-05 |
| ENSMUSG00000020717 | 11 | Pecam1 | protein_coding | -1.593749625 | 0 | 0.054564112 | -0.580324 | -1.019938 | 0.005102471 | 0.000188414 |
| ENSMUSG00000038569 | 5 | Rad9b | protein_coding | -1.651456351 | 0 | -0.304091559 | -1.028609 | -0.661891 | 0.027219177 | 0.001725858 |
| ENSMUSG00000038370 | 1 | Pcp4l1 | protein_coding | -1.66956346 | 0 | -1.313115994 | 4.408432 | 5.006784 | 4.97577E-05 | 6.68E-07 |
| ENSMUSG00000050556 | 2 | Kcnb1 | protein_coding | -1.701192569 | 0 | -0.391224882 | -0.362468 | 5.559979 | 7.83954E-05 | 1.13635E-06 |
| ENSMUSG00000037922 | 3 | Bank1 | protein_coding | -1.701299429 | 0 | -0.310856781 | -0.332745 | -2.015887 | 2.40E-08 | 2.72E-11 |
| ENSMUSG00000064115 | 16 | Cadm2 | protein_coding | -1.709802654 | 0 | -1.891547225 | -2.158832 | 1.570416 | 0.009150105 | 0.0004016 |
| ENSMUSG00000040820 | 16 | Hlcs | protein_coding | -1.729300551 | 0 | 1.296178936 | -2.169114 | 1.1519 | 0.025274875 | 0.001546595 |
| ENSMUSG00000022054 | 14 | Nefm | protein_coding | -1.766298664 | 0 | -0.57291344 | -1.920017 | 2.364325 | 0.033320708 | 0.002285419 |
| ENSMUSG00000028391 | 4 | Wdr31 | protein_coding | -1.81817611 | 0 | -1.645055253 | -1.767429 | 1.419138 | 0.040768044 | 0.003086396 |
| ENSMUSG00000073413 | 17 | Ly6g6d | protein_coding | -1.850063247 | 0 | -0.207697522 | 0.539627 | -4.713311 | 0.000224816 | 4.15079E-06 |
| ENSMUSG00000092035 | 6 | Peg10 | protein_coding | -1.871592832 | 0 | -0.402280148 | -1.665239 | -3.928652 | 0.008999628 | 0.000391352 |
| ENSMUSG00000028524 | 4 | Sgip1 | protein_coding | -1.877462892 | 0 | -0.9010449 | -1.785928 | 2.224787 | 0.010674836 | 0.00048671 |
| ENSMUSG00000048388 | 2 | Fam171b | protein_coding | -1.906063957 | 0 | 1.246874382 | 0.5123 | 2.956757 | 0.017575594 | 0.000969227 |
| ENSMUSG00000048126 | 1 | Col6a3 | protein_coding | -1.926373064 | 0 | -0.633227763 | -0.11634 | -5.216781 | 0.000123669 | 2.0029E-06 |
| ENSMUSG00000022537 | 16 | Tmem44 | protein_coding | -1.972087515 | 0 | -0.044253037 | -0.938983 | -3.79703 | 6.85E-07 | 1.78E-09 |
| ENSMUSG00000072720 | 5 | Myo18b | protein_coding | -1.981338586 | 0 | 0.574044917 | 0.125206 | -4.545934 | 4.35556E-05 | 5.54E-07 |
| ENSMUSG00000032530 | 9 | Lyzl4 | protein_coding | -2.027777724 | 0 | 0.801989981 | -0.505741 | -2.666789 | 0.000645493 | 1.5054E-05 |
| ENSMUSG00000071648 | 19 | Rom1 | protein_coding | -2.028317465 | 0 | -1.27373557 | -0.075151 | 8.160163 | 0.000372395 | 7.68975E-06 |
| ENSMUSG00000022055 | 14 | Nefl | protein_coding | -2.044463289 | 0 | 0.759392863 | -0.552479 | 3.348078 | 0.004129086 | 0.000142201 |
| ENSMUSG00000027200 | 2 | Sema6d | protein_coding | -2.051855078 | 0 | -0.163293496 | -0.904783 | -5.206867 | 1.22106E-06 | 3.96E-09 |
| ENSMUSG00000042589 | 5 | Cux2 | protein_coding | -2.162541678 | 0 | -0.791701626 | -1.13636 | -3.160558 | 0.046458172 | 0.00369264 |
| ENSMUSG00000078920 | 11 | Ifi47 | protein_coding | -2.203013097 | 0 | -0.497200491 | 1.668937 | 1.884008 | 0.005116918 | 0.00019044 |
| ENSMUSG00000015843 | 1 | Rxrg | protein_coding | -2.246884599 | 0 | -1.772996335 | 1.951926 | 1.473222 | 0.002253834 | 6.6243E-05 |
| ENSMUSG00000035864 | 10 | Syt1 | protein_coding | -2.280554952 | 0 | -1.537158013 | -1.091725 | 1.920703 | 0.002941425 | 9.26564E-05 |
| ENSMUSG00000031990 | 9 | Jam3 | protein_coding | -2.312341707 | 0 | -0.115604531 | -1.255647 | -2.736616 | 0.04096049 | 0.003107562 |
| ENSMUSG00000034009 | 3 | Rxfp1 | protein_coding | -2.320603697 | 0 | 0.044097678 | -2.661774 | -3.533175 | 0.011262288 | 0.000522576 |
| ENSMUSG00000050640 | 5 | Tmem150c | protein_coding | -2.346184168 | 0 | -1.840468265 | -0.063009 | 2.462616 | 0.008225206 | 0.000347082 |
| ENSMUSG00000022528 | 16 | Hes1 | protein_coding | -2.377998511 | 0 | -0.832711099 | -1.223191 | -2.014532 | 0.00012665 | 2.06143E-06 |
| ENSMUSG00000037855 | 10 | Zfp365 | protein_coding | -2.38109402 | 0 | -2.590725774 | -2.289997 | 2.27824 | 0.004799667 | 0.000172569 |
| ENSMUSG00000000861 | 11 | Bcl11a | protein_coding | -2.425218001 | 0 | -0.224710727 | 0.42346 | -1.7158 | 0.025196717 | 0.00153641 |
| ENSMUSG00000022332 | 15 | Khdrbs3 | protein_coding | -2.598291612 | 0 | 0.156629351 | -1.269997 | -2.445429 | 0.00048284 | 1.0536E-05 |
| ENSMUSG00000045103 | X | Dmd | protein_coding | -2.648142975 | 0 | -1.324567492 | -2.187351 | 1.478509 | 0.013275273 | 0.000666791 |
| ENSMUSG00000025466 | 7 | Fuom | protein_coding | -2.67109415 | 0 | -3.537881133 | -1.669 | -0.229554 | 0.008926128 | 0.00038387 |
| ENSMUSG00000037820 | 2 | Tgm2 | protein_coding | -2.699513597 | 0 | -0.331863238 | -0.702185 | -0.875321 | 0.000192069 | 3.39065E-06 |
| ENSMUSG00000036949 | 2 | Slc39a12 | protein_coding | -2.69981192 | 0 | -0.400045343 | -1.172032 | -5.290448 | 0.000117611 | 1.88575E-06 |
| ENSMUSG00000022037 | 14 | Clu | protein_coding | -2.736971275 | 0 | -0.666566441 | -3.299907 | -0.308371 | 0.010065091 | 0.000452822 |
| ENSMUSG00000024617 | 18 | Camk2a | protein_coding | -2.750274616 | 0 | 0.368834121 | -0.677145 | -2.837565 | 0.023468732 | 0.001410138 |
| ENSMUSG00000028176 | 3 | Lrrc7 | protein_coding | -2.763860652 | 0 | -0.09042325 | 1.201129 | -2.116499 | 0.038508643 | 0.002818999 |
| ENSMUSG00000003949 | 11 | Hlf | protein_coding | -2.899276298 | 0 | -0.625607142 | -1.741933 | -4.470451 | 9.72945E-06 | 7.33E-08 |
| ENSMUSG00000002341 | 8 | Ncan | protein_coding | -2.929875591 | 0 | -0.458895823 | 0.283506 | -2.665439 | 0.035778373 | 0.002516854 |
| ENSMUSG00000047342 | 11 | Zfp286 | protein_coding | -3.01753515 | 0 | -0.296685484 | -0.415653 | -2.057488 | 0.008211802 | 0.000343793 |
| ENSMUSG00000028989 | 4 | Angptl7 | protein_coding | -3.195883158 | 0 | -0.524441566 | -0.782991 | -2.827299 | 0.004927687 | 0.000179167 |
| ENSMUSG00000049436 | 16 | Upk1b | protein_coding | -3.211806654 | 0 | -0.196458759 | -0.835823 | -2.936567 | 5.18E-09 | 3.77E-12 |
| ENSMUSG00000058022 | 13 | Adtrp | protein_coding | -3.491532284 | 0 | 1.643747519 | -0.50774 | -2.134466 | 0.004041827 | 0.000136811 |
| ENSMUSG00000063354 | 15 | Slc39a4 | protein_coding | -3.513202226 | 0 | -1.436098783 | 0.930552 | 0.843997 | 0.000569095 | 1.27653E-05 |
| ENSMUSG00000041120 | 4 | Nbl1 | protein_coding | -3.576654135 | 0 | -1.440198064 | -0.175448 | -3.331524 | 0.013770631 | 0.000703641 |

| | | | | | | | | | | |
|---|---|---|---|---|---|---|---|---|---|---|
| ENSMUSG00000042371 | 11 | Slc5a10 | protein_coding | -3.652244189 | 0 | -0.150499794 | -1.18289 | -4.192649 | 0.003527542 | 0.00011569 |
| ENSMUSG00000013766 | 17 | Ly6g6e | protein_coding | -4.086322564 | 0 | -0.703052559 | -1.062197 | -2.968013 | 0.004129086 | 0.000142344 |
| ENSMUSG00000059213 | 15 | Ddn | protein_coding | -4.232323673 | 0 | 0.600453658 | -0.930605 | -3.965488 | 9.03218E-05 | 1.33848E-06 |
| ENSMUSG00000030209 | 6 | Grin2b | protein_coding | -4.290003156 | 0 | -0.332918466 | 0.237123 | -2.351698 | 0.009814736 | 0.000439513 |
| ENSMUSG00000046093 | 4 | Hpcal4 | protein_coding | -4.422100245 | 0 | -1.300343876 | -0.009516 | -2.361303 | 0.009788955 | 0.000437566 |
| ENSMUSG00000005045 | 4 | Chd5 | protein_coding | -4.438032018 | 0 | -0.635923056 | -0.042655 | -2.364939 | 0.002368171 | 7.03797E-05 |
| ENSMUSG00000048978 | 13 | Nrsn1 | protein_coding | -4.557186288 | 0 | -1.836421387 | -0.007115 | -3.058859 | 0.022519338 | 0.001334858 |
| ENSMUSG00000040016 | 3 | Ptger3 | protein_coding | -5.205717039 | 0 | -3.08597293 | -2.490741 | -3.063193 | 0.004129086 | 0.000142293 |
| ENSMUSG00000053310 | 9 | Nrgn | protein_coding | -5.265409699 | 0 | -0.996089765 | -1.041994 | -5.422834 | 9.11088E-06 | 6.64E-08 |

Genes that are related to immune functions (196)

| Gene.Name | cerebellum | cortex | hippocampus | ob | retina |
|---|---|---|---|---|---|
| Spp1 | 6.130871893 | 0 | 2.110631952 | 5.894457 | 8.079708 |
| Lgals3 | 4.128110423 | 0 | 1.28040307 | 3.426976 | 5.630613 |
| Hp | 3.951189841 | 0 | 3.044361538 | 5.169951 | 1.111205 |
| Serpinb1a | 3.933228262 | 0 | 1.025648633 | 4.345552 | 0.972788 |
| Alpk1 | 3.778678259 | 0 | 3.257317856 | 3.34867 | 3.47122 |
| Ifi206 | 3.729584851 | 0 | 0.490824938 | 4.183982 | 4.260252 |
| Tnfsf8 | 3.649454062 | 0 | 0.981869116 | -0.74267 | 5.328515 |
| Fcnb | 3.590855302 | 0 | 2.568371157 | 4.999078 | 0.769484 |
| St18 | 3.531775743 | 0 | 1.312896185 | -0.03589 | 3.07373 |
| Rsad2 | 3.434427312 | 0 | 0.959881235 | 3.13731 | 4.759717 |
| B3gnt7 | 3.386073223 | 0 | 0.819099989 | 2.51634 | 4.71227 |
| Adamts1 | 3.344208329 | 0 | -0.547392689 | -1.97909 | 1.923514 |
| Cyp4f18 | 3.256435633 | 0 | 0.354558377 | 1.021806 | 6.332474 |
| Hspa1b | 3.225907883 | 0 | 0.802603996 | -0.39581 | -0.40529 |
| Hspa1a | 3.205567175 | 0 | -0.01094095 | -0.21483 | 1.008945 |
| Ltf | 3.196058839 | 0 | 1.839351386 | 4.802103 | -1.74639 |
| Oas2 | 3.188620211 | 0 | 1.555165453 | 4.717833 | 3.852152 |
| Il18rap | 3.032073139 | 0 | 0.171030813 | 0.676228 | 3.637547 |
| 1810011O10Rik | 3.013310432 | 0 | 0.965462159 | -2.24143 | 4.188551 |
| Gch1 | 2.955644753 | 0 | 1.399473051 | 3.112621 | 2.358734 |
| Slc2a6 | 2.923032705 | 0 | 1.722958662 | 2.79965 | 5.237169 |
| Cacna2d2 | 2.885060464 | 0 | -1.05680879 | -0.75608 | 2.020937 |
| Lpl | 2.878604358 | 0 | 1.00115449 | 2.521777 | 2.413033 |
| Trem3 | 2.877504377 | 0 | 1.362237272 | 5.050046 | 1.071254 |
| H2-Aa | 2.763839343 | 0 | 0.767134826 | 2.745465 | 0.861272 |
| AB124611 | 2.7305726 | 0 | 1.730521329 | 3.492226 | 4.490252 |
| Nlrc5 | 2.677741073 | 0 | -2.007085048 | 1.880921 | 3.386167 |
| F830016B08Rik | 2.651569971 | 0 | 0.138740687 | 1.673826 | -1.45007 |
| Cd244 | 2.589690726 | 0 | 0.383560156 | 1.041788 | 4.498612 |
| Ifi207 | 2.561614701 | 0 | 0.839202023 | 2.890929 | 2.862795 |
| H2-Eb1 | 2.549252212 | 0 | 0.554757167 | 3.064072 | 1.123164 |
| Tnfsf13b | 2.485867573 | 0 | 0.736677785 | 0.631162 | 4.003227 |
| Cd93 | 2.48559373 | 0 | 0.754709423 | 2.797001 | -0.86717 |
| Gzmm | 2.427753326 | 0 | 0.589743301 | 0.18375 | 3.68567 |
| Pdcd1 | 2.380386121 | 0 | 0.217601256 | -0.91409 | 4.894188 |
| Fgr | 2.29192519 | 0 | 1.365197863 | 1.578893 | 3.242645 |
| Ifi204 | 2.253637942 | 0 | -0.604461152 | 1.840358 | 3.289616 |
| Axl | 2.220411384 | 0 | -0.791637238 | 2.306578 | 4.338706 |
| Ifitm6 | 2.203127999 | 0 | 1.646961178 | 3.573787 | -1.19861 |
| Ifi211 | 2.20227544 | 0 | 0.020324276 | 1.353624 | 3.517917 |
| Chil3 | 2.183138169 | 0 | 1.13645529 | 3.8735 | -2.45684 |
| Pbk | 2.176702503 | 0 | 3.730357443 | 4.213626 | 4.36119 |
| H2-Ab1 | 2.146275182 | 0 | 0.613631593 | 2.015808 | 1.919613 |
| Cd74 | 2.125364882 | 0 | 0.55932368 | 2.390668 | 1.489495 |
| Skint3 | 2.060287582 | 0 | -1.747842511 | -1.84397 | 0.717866 |
| Cybb | 2.052838911 | 0 | 0.797768693 | 2.290361 | 3.073937 |
| S100a9 | 2.03991683 | 0 | 0.655996476 | 3.489083 | -3.67177 |

| | | | | | |
|---|---|---|---|---|---|
| Scimp | 1.989417757 | 0 | 0.184618755 | 1.347371 | 3.633955 |
| Cd44 | 1.936389141 | 0 | -0.498217493 | 1.397863 | 3.006067 |
| Lsp1 | 1.919165727 | 0 | 0.187969374 | 0.610754 | 3.153445 |
| Ascl4 | 1.899016084 | 0 | -0.911591927 | -0.61086 | 3.041029 |
| Scg2 | 1.879521662 | 0 | 2.848658398 | 4.185677 | 4.204894 |
| Anxa1 | 1.81924572 | 0 | 0.342850856 | 3.304358 | -1.79376 |
| Chst1 | 1.815490821 | 0 | 1.437993686 | 1.925774 | 3.818197 |
| Oasl2 | 1.795715342 | 0 | -2.173779349 | 1.554389 | 3.065798 |
| Jag1 | 1.683073011 | 0 | 0.71943573 | -0.07527 | 4.254852 |
| Il1r1 | 1.673123303 | 0 | -2.119965163 | -1.18874 | 0.933983 |
| Milr1 | 1.648578416 | 0 | 1.016278025 | 1.772892 | 2.714059 |
| Cxcl10 | 1.618113008 | 0 | -0.606105827 | 3.036347 | 5.714292 |
| Nfkbiz | 1.605494279 | 0 | 0.708755375 | 1.342019 | 1.632144 |
| B430306N03Rik | 1.602981139 | 0 | 0.800115623 | 1.586079 | 2.506014 |
| Ngp | 1.596930891 | 0 | 0.709061081 | 3.281334 | -4.19013 |
| Adcy1 | 1.588379871 | 0 | -0.859806731 | -3.13542 | 0.968 |
| Zbp1 | 1.579878847 | 0 | -1.087222155 | 2.380799 | 3.499346 |
| Nefh | 1.576741291 | 0 | 0.057789566 | -0.83467 | 4.011005 |
| Ifi209 | 1.553626858 | 0 | -0.775775013 | 0.683676 | 2.340976 |
| Ccl2 | 1.523525246 | 0 | 0.665432576 | 1.767713 | 1.964214 |
| Oasl1 | 1.47976849 | 0 | -0.781369304 | 1.201743 | 3.597087 |
| H2-Q6 | 1.424454766 | 0 | -0.590746461 | 2.285191 | 2.683226 |
| Mylpf | 1.400308735 | 0 | 2.663944733 | 2.316952 | 4.215127 |
| Ccl4 | 1.362406896 | 0 | 0.66604151 | 1.539092 | 1.835519 |
| Ccl3 | 1.35485491 | 0 | 0.880118146 | 1.736307 | 2.622812 |
| Sp100 | 1.337678102 | 0 | -0.799619011 | 1.019911 | 1.987718 |
| Ccl12 | 1.316764421 | 0 | -0.234321432 | 2.163093 | 3.337355 |
| Il2rg | 1.277908709 | 0 | 0.086697919 | 2.671912 | 3.134673 |
| Camp | 1.248168749 | 0 | 0.399393367 | 3.432306 | -4.05325 |
| Lyz2 | 1.245187206 | 0 | 0.298421084 | 1.860094 | 1.192964 |
| S100a8 | 1.242062331 | 0 | 0.742322929 | 3.466899 | -4.35304 |
| Pkib | 1.238783605 | 0 | 0.489241208 | 0.633901 | 1.739605 |
| Csf2ra | 1.23037946 | 0 | 0.42888742 | 0.444896 | 1.632927 |
| Tnf | 1.203019381 | 0 | -0.08397666 | 0.722149 | 3.369352 |
| Phf11a | 1.172637684 | 0 | 1.063429536 | 2.702794 | 6.534529 |
| Cfp | 1.169955319 | 0 | 0.956314205 | 1.359501 | -1.59482 |
| Ifi213 | 1.143327926 | 0 | 0.010841621 | 2.231345 | 2.40534 |
| Slfn2 | 1.141011207 | 0 | -0.052820031 | 1.034402 | 1.530552 |
| Slamf6 | 1.139540744 | 0 | 0.015942786 | 0.845531 | 1.238045 |
| Pf4 | 1.133953422 | 0 | 0.671735312 | 1.615624 | -0.79004 |
| Ly9 | 1.115620493 | 0 | -1.315567587 | 0.91389 | 1.686722 |
| Cd163 | 1.106020813 | 0 | 0.39783313 | 1.580175 | -2.09005 |
| Ifit3b | 1.085227233 | 0 | -2.054939488 | 1.574942 | 3.565212 |
| Ptger4 | 1.084989857 | 0 | 2.175186386 | 4.036334 | 2.534857 |
| Clec10a | 1.047857592 | 0 | 0.084126464 | 1.185873 | -3.71284 |
| Ifit3 | 1.045014111 | 0 | -0.872782344 | 0.181265 | 2.152417 |
| Lat2 | 1.038443376 | 0 | 0.44965854 | 0.376814 | 1.846815 |
| Emilin1 | 1.012446138 | 0 | 1.184717129 | 1.978568 | 3.594512 |
| S1pr4 | 1.001386124 | 0 | 1.259438073 | 2.650417 | -1.85773 |

| | | | | | |
|---|---|---|---|---|---|
| Ifitm3 | 0.957247399 | 0 | -0.547974772 | 2.031251 | 2.568357 |
| Irf7 | 0.919813938 | 0 | -0.341467784 | 1.076964 | 1.932754 |
| Clec4e | 0.845707485 | 0 | 0.611680164 | -0.03589 | 4.45315 |
| Pla2r1 | 0.845707485 | 0 | -0.336623544 | 0.578671 | 4.639159 |
| Phf11d | 0.803753403 | 0 | 0.058266484 | 0.406721 | 2.100324 |
| H2-D1 | 0.79872045 | 0 | -0.235613737 | 1.016082 | 1.902674 |
| Cd22 | 0.751934928 | 0 | -0.574999916 | 1.744083 | 0.647736 |
| Bcl2a1a | 0.702951996 | 0 | -0.058413094 | 0.600727 | 1.672781 |
| Klhl6 | 0.698811276 | 0 | 0.385204044 | 0.593132 | 1.117261 |
| Ifit2 | 0.684880985 | 0 | -1.154984955 | 1.888731 | 2.6972 |
| Mbp | 0.677748461 | 0 | 0.99818058 | 0.05094 | -1.63372 |
| Cst7 | 0.654528805 | 0 | 0.289194958 | 2.325981 | 2.062429 |
| Ifit1bl1 | 0.649709867 | 0 | 0.229455678 | 1.078606 | 3.48942 |
| Cd300lf | 0.642235207 | 0 | -0.013678992 | 1.175455 | 2.449149 |
| Ccdc115 | 0.633556021 | 0 | -0.032932861 | -0.00829 | 1.134484 |
| Cd180 | 0.627737375 | 0 | 0.317508014 | 0.409941 | 1.165195 |
| Cxcl13 | 0.627406106 | 0 | 0.425068795 | 2.545504 | 4.326694 |
| Rnase6 | 0.60285831 | 0 | -0.233127751 | 0.180905 | 1.542308 |
| C4b | 0.582579793 | 0 | 0.429990026 | 2.133183 | 1.470262 |
| Ctsc | 0.576015024 | 0 | -0.185359303 | 0.237967 | 0.964521 |
| Pirb | 0.573465051 | 0 | -0.329036235 | -0.35581 | 1.309689 |
| Ly6a | 0.520683348 | 0 | -0.152146786 | 1.405857 | 1.789074 |
| Ifit1 | 0.517675269 | 0 | -0.121120074 | 2.663941 | 2.753053 |
| Gpnmb | 0.515763083 | 0 | 0.05678378 | 0.578671 | 7.097569 |
| H2-M2 | 0.507835985 | 0 | 0.05678378 | 0.578671 | 4.62052 |
| Casp1 | 0.47622703 | 0 | -0.287461567 | 0.318614 | 0.776138 |
| Fth1 | 0.472328094 | 0 | 0.033033417 | 0.557483 | 1.562967 |
| Ly86 | 0.427764806 | 0 | 0.313323432 | 0.499933 | 1.09809 |
| Acer3 | 0.415399633 | 0 | 0.225065175 | 0.251045 | 1.144731 |
| Hsph1 | 0.404573921 | 0 | -0.216568392 | -0.4751 | -1.17204 |
| Tlr2 | 0.393167344 | 0 | -0.281924404 | 0.438684 | 0.857982 |
| Tpd52 | 0.370471123 | 0 | -0.286318917 | 0.330052 | 0.723325 |
| Fcgr4 | 0.349477778 | 0 | -0.17116871 | 1.120745 | 2.748538 |
| Evi2 | 0.33963025 | 0 | -1.411684819 | -0.23641 | 1.320355 |
| Hcar2 | 0.336676113 | 0 | 1.039856169 | 1.058514 | 3.845916 |
| Mgl2 | 0.327123415 | 0 | -0.016004941 | 0.460365 | -3.38056 |
| Cd48 | 0.318499186 | 0 | 0.034334065 | 0.222058 | 1.40397 |
| H2-K1 | 0.301667965 | 0 | -0.521808484 | 0.573946 | 1.046261 |
| Ctla2a | 0.297267829 | 0 | -0.336623544 | 1.144864 | 3.770672 |
| Usp18 | 0.27746345 | 0 | -0.829992592 | 1.157908 | 1.863214 |
| Ldlr | 0.273286774 | 0 | -0.242299548 | -0.08628 | 2.261462 |
| Patz1 | 0.255602458 | 0 | 0.434276189 | -0.59849 | -1.2644 |
| Cd34 | 0.233080064 | 0 | 0.133968749 | 0.071852 | 1.037873 |
| H2-M3 | 0.179648385 | 0 | -0.271894115 | 0.045861 | 0.819357 |
| Ccl5 | 0.167008878 | 0 | 0.030109871 | 1.415699 | 4.942751 |
| Tsc22d3 | 0.155586266 | 0 | -0.580080354 | -0.838 | 0.173534 |
| Scarb1 | 0.140533447 | 0 | 0.013377229 | 0.571991 | 1.279092 |
| Lgals3bp | 0.128810758 | 0 | -0.641760447 | 0.41642 | 0.836254 |
| Pdc | 0.100910842 | 0 | 0.157472865 | -0.43082 | 9.154045 |

| | | | | | |
|---|---|---|---|---|---|
| Itgax | 0.089890996 | 0 | -0.085712028 | 1.562794 | 1.307967 |
| B2m | 0.08324408 | 0 | -0.203467387 | 0.574726 | 1.028936 |
| C5ar1 | 0.062302046 | 0 | -0.182550591 | -0.152 | 0.932074 |
| Ccl6 | 0.038924659 | 0 | 0.449401429 | 0.64 | 1.540994 |
| Isg15 | -0.02504931 | 0 | -0.980244131 | 1.396788 | 2.194882 |
| Fgl2 | -0.054346679 | 0 | -0.721130258 | 1.92159 | 2.508752 |
| Cd9 | -0.065588009 | 0 | -0.126196089 | -0.06827 | 1.049212 |
| Prdx1 | -0.0740433 | 0 | -0.129899574 | 0.375362 | 0.995949 |
| Mrc1 | -0.075354294 | 0 | -0.524356777 | 0.571175 | -2.61238 |
| Gbp7 | -0.097860128 | 0 | -0.153205672 | 0.065278 | -1.16621 |
| Mxra8 | -0.101243797 | 0 | 0.208355044 | 0.394797 | 3.580575 |
| Tulp1 | -0.126991789 | 0 | 0.251714957 | 0.342106 | 8.706932 |
| Sh2d1a | -0.126991789 | 0 | -0.336623544 | 0.818492 | 3.902507 |
| Pla2g5 | -0.126991789 | 0 | -0.336623544 | -0.03589 | 3.837718 |
| Cd81 | -0.144840008 | 0 | -0.132772424 | -0.17577 | 0.878565 |
| Adora1 | -0.151941938 | 0 | 0.256231611 | 0.83927 | 3.40565 |
| Slc12a2 | -0.157180509 | 0 | -0.294506587 | 0.272393 | -1.08307 |
| Itga9 | -0.183911758 | 0 | 0.096385878 | 0.074736 | -0.92319 |
| Tlr5 | -0.204779762 | 0 | 0.155867205 | -0.99023 | -3.38283 |
| Tspan32 | -0.208327982 | 0 | -0.315328226 | 0.288123 | 2.043549 |
| Stab1 | -0.24430581 | 0 | 0.008143334 | -0.06552 | -1.51446 |
| Cd59a | -0.245947561 | 0 | 1.057156061 | 0.404428 | 3.526814 |
| Ykt6 | -0.270950347 | 0 | 0.52350826 | -0.32792 | -0.66394 |
| Dpf3 | -0.277700555 | 0 | -0.100647145 | -0.61086 | 3.906054 |
| Paip1 | -0.28690937 | 0 | 0.978034053 | 0.557741 | -0.16887 |
| Ebi3 | -0.287936006 | 0 | -0.783571238 | -0.14577 | 0.420567 |
| Pkm | -0.292542855 | 0 | -0.091168031 | -0.08377 | 1.005809 |
| Cln6 | -0.297890626 | 0 | -2.115566945 | 0.621759 | 0.2619 |
| Srgn | -0.321064633 | 0 | -0.161382087 | -0.10051 | 0.829235 |
| Kdm6b | -0.441030231 | 0 | 0.741606812 | 0.076136 | -0.80442 |
| Abcg1 | -0.594156857 | 0 | 0.335370058 | 0.372576 | -2.12816 |
| Prkca | -0.617612486 | 0 | -0.004922025 | -0.48198 | -1.5271 |
| Pear1 | -0.679854934 | 0 | 0.3099205 | -1.21535 | 0.62828 |
| Ccl9 | -0.681679627 | 0 | 0.375812488 | -0.13876 | 0.401325 |
| Ubqln2 | -0.693960089 | 0 | -0.537151514 | -1.22621 | -1.03745 |
| Prkce | -0.702192106 | 0 | 0.137257137 | -0.60737 | -1.76646 |
| Tnfrsf17 | -0.758787405 | 0 | 0.249369385 | -0.15036 | -0.96294 |
| S1pr1 | -0.774097895 | 0 | -0.174549389 | -0.10905 | -1.06804 |
| Pilra | -0.796760434 | 0 | -0.758683727 | -1.06437 | 1.181936 |
| Il7r | -0.877750635 | 0 | -0.044628012 | -0.57177 | -1.31606 |
| H2-Q7 | -0.905032041 | 0 | 0.191246881 | 1.18118 | 2.262091 |
| Pde4b | -1.005775276 | 0 | -0.70289739 | -0.75475 | -1.07111 |
| Ccl24 | -1.106653885 | 0 | 0.059130297 | 0.338197 | -3.55982 |
| Map4k1 | -1.160980299 | 0 | 0.729515385 | 0.475443 | 2.823158 |
| Trim30c | -1.163669971 | 0 | -0.449933705 | 0.385286 | -1.60563 |
| Pecam1 | -1.593749625 | 0 | 0.054564112 | -0.58032 | -1.01994 |
| Ifi47 | -2.203013097 | 0 | -0.497200491 | 1.668937 | 1.884008 |
| Jam3 | -2.312341707 | 0 | -0.115604531 | -1.25565 | -2.73662 |
| Clu | -2.736971275 | 0 | -0.666566441 | -3.29991 | -0.30837 |

| | | | | | |
|---|---|---|---|---|---|
| Nbl1 | -3.576654135 | 0 | -1.440198064 | -0.17545 | -3.33152 |
| Klf2 | 5.04871599 | 0 | 1.44755254 | 2.851827 | 5.431431 |
