## Supplemental file 3 for "Regional and sub-regional microglial heterogeneity in the steady-state mouse brain and retina"

| Metrics | Olfactory bulb | Cortex | Hippocampus | Cerebellum | Retina |
|---|---|---|---|---|---|
| Estimated number of cells | 4,949 | 4,563 | 6,855 | 5,724 | 5,004 |
| Mean. Reads per Cell | 63,145 | 88,598 | 65,631 | 66,753 | 74,054 |
| Median Genes per Cell | 2,526 | 2,613 | 2,417 | 2,171 | 2,155 |
| Number. Of. Reads | 312,505,632 | 404,273,630 | 449,900,554 | 382,096,199 | 370,567,108 |
| Valid Barcodes | 98.20% | 98.20% | 98.10% | 98.20% | 98.10% |
| Sequencing Saturation | 75.90% | 82.70% | 77.70% | 78.80% | 83.20% |
| Q30.Bases in Barcode | 96.60% | 96.60% | 96.80% | 96.70% | 96.70% |
| Q30.Bases.in.RNA.Read | 92.60% | 92.10% | 93.50% | 92.90% | 92.80% |
| Q30.Bases.in.UMI | 96.10% | 96.20% | 96.40% | 96.20% | 96.30% |
| Reads Mapped to Genome | 95.90% | 95.60% | 94.60% | 95.60% | 96.20% |
| Reads Mapped Confidently to Genome | 94.10% | 93.90% | 92.80% | 93.80% | 94.40% |
| Reads Mapped Confidently to Intergenic Regions | 3.50% | 3.50% | 3.50% | 3.70% | 3.60% |
| Reads Mapped Confidently to Intronic Regions | 40.40% | 39.70% | 42.10% | 46.40% | 42.80% |
| Reads Mapped Confidently to Exonic Regions | 50.20% | 50.70% | 47.20% | 43.80% | 47.90% |
| Reads Mapped Confidently to Transcriptome | 47.00% | 47.50% | 44.20% | 40.60% | 44.50% |
| Reads Mapped Antisense to Gene | 2.00% | 2.00% | 2.00% | 2.00% | 2.20% |
| Fraction Reads in Cells | 96.50% | 94.70% | 96.00% | 95.80% | 95.80% |
| Total Genes Detected | 17,828 | 18,101 | 18,488 | 18,411 | 18,268 |
| Median UMI Counts per Cell | 6,303 | 6,491 | 5,699 | 4,757 | 4,849 |
